# Individual word representations dissociate from linguistic context along a cortical unimodal to heteromodal gradient

**DOI:** 10.1101/2023.04.25.538257

**Authors:** Susanne Eisenhauer, Tirso Rene del Jesus Gonzalez Alam, Piers L. Cornelissen, Jonathan Smallwood, Elizabeth Jefferies

## Abstract

Language comprehension involves multiple hierarchical processing stages across time, space, and levels of representation. When processing a word, the sensory input is transformed into increasingly abstract representations that need to be integrated with the linguistic context. Thus, language comprehension involves both input-driven as well as context-dependent processes. While neuroimaging research has traditionally focused on mapping individual brain regions to the distinct underlying processes, recent studies indicate that whole-brain distributed patterns of cortical activation might be highly relevant for cognitive functions, including language. One such pattern, based on resting-state connectivity, is the ‘principal cortical gradient’, which dissociates sensory from heteromodal brain regions. The present study investigated the extent to which this gradient provides an organizational principle underlying language function, using a multimodal neuroimaging dataset of functional magnetic resonance imaging (fMRI) and magnetoencephalography (MEG) recordings from 102 participants during sentence reading. We found that the brain response to individual representations of a word (word length, orthographic distance and word frequency), which reflect visual, orthographic, and lexical properties, gradually increases towards the sensory end of the gradient. Although these properties showed opposite effect directions in fMRI and MEG, their association with the sensory end of the gradient was consistent across both neuroimaging modalities. In contrast, MEG revealed that properties reflecting a word’s relation to its linguistic context (semantic similarity and position within the sentence) involve the heteromodal end of the gradient to a stronger extent. This dissociation between individual word and contextual properties was stable across earlier and later time windows during word presentation, indicating interactive processing of word representations and linguistic context at opposing ends of the principal gradient. To conclude, our findings indicate that the principal gradient underlies the organization of a range of linguistic representations while supporting a gradual distinction between context-independent and context-dependent representations. Furthermore, the gradient reveals convergent patterns across neuroimaging modalities (similar location along the gradient) in the presence of divergent responses (opposite effect directions).

## 1 Introduction

Language comprehension involves multiple hierarchical processing stages that are organised across time, space, and levels of representation (Carreiras et al., 2014; Dikker et al., 2020; Hickok and Poeppel, 2007). During the processing of a word, its cognitive representations are thought to become increasingly abstract over time: the initial perceptual representation of the sensory input (i.e., individual letters/phonemes) is transformed into sub-lexical orthographic and phonological representations (letter/sound combinations) and into lexical-semantic representations (word meaning) that are integrated with the preceding context (e.g., Grainger and Ziegler, 2011). The brain regions that represent visual, orthographic, phonological and semantic information about words are thought to be situated on ‘lower’ to ‘higher’ regions along a cortical hierarchy (Carreiras et al., 2014; Friederici, 2011). However, there is still controversy about how these representations are organised on the cortical surface and accessed through time.

Some researchers have proposed that the focus of processing during visual word recognition moves systematically along this representational hierarchy through time: initial visual processing of the input is associated with early activation (∼ 100 ms) in the occipital cortex, followed by orthographic processing in the inferior temporal cortex (∼150 to 300 ms), and then lexical-semantic processing in a network of regions including the angular gyrus, anterior temporal, and inferior frontal cortex (∼ 300 to 500 ms; e.g., Eisenhauer et al., 2019; Simon et al., 2012; Vartiainen et al., 2011; for reviews, see Carreiras et al., 2014; Dikker et al., 2020). Other researchers have identified early lexical-semantic responses in prefrontal and anterior temporal cortex (Clarke et al., 2011; 2013; Cornelissen et al., 2009; Eisenhauer et al., 2022; Kaestner et al., 2021; Mollo et al., 2017; 2018; Teige et al., 2019), suggesting that visual word recognition is highly interactive and visual feedforward processes are shaped by early top-down predictions (Price and Devlin, 2011). By this second view, there may be little systematic change in the timing of responses along the cortical hierarchy, although there might still be a systematic shift in the information represented in different brain areas.

In addition, while the occipital, inferior and anterior temporal, inferior prefrontal and inferior parietal regions associated with different levels of language processing are widely distributed across the left hemisphere, other researchers have proposed that these areas are organized on the cortical surface in a systematic fashion that reflects a visual to semantic processing hierarchy (Lambon Ralph et al., 2017). Previous studies have identified gradual shifts in the location of brain activation that reflect the abstractness and complexity of linguistic representations: for example, the gradual change from letter to orthographic subunit to whole-word representations is associated with a posterior-to-anterior shift in activation in occipital-temporal cortex (e.g., Dehaene et al., 2005; Lerma-Usabiaga et al., 2018; Vinckier et al., 2007). In addition to this local gradient, global gradients, i.e., gradual transitions in brain activation patterns between neighbouring regions across the whole brain, have also been linked to language processing: Gradual transitions between brain regions responding most strongly to perceptual semantic features versus more abstract (e.g., emotional/social) features were identified across the whole brain (e.g., Huth et al., 2012; 2016; Popham et al., 2021). These findings indicate that gradual transitions in the complexity of linguistic representations might involve corresponding gradual shifts in the implicated brain regions along the cortex.

In recent work examining the topographical organization of cortex, variance decomposition of intrinsic connectivity was used to reveal a whole-brain ‘principal gradient’ accounting for the most variance in intrinsic connectivity derived from resting-state functional magnetic resonance imaging (fMRI) across individuals (Margulies et al., 2016). This gradient captures gradual changes in connectivity that are maximally different between sensory/unimodal (visual, somatosensory and motor) and heteromodal regions including the default mode network. Meta-analytic decoding indicates that this principal gradient dissociates sensory, input-driven, or novelty-based processes from abstract, integration- or memory-based processes (Margulies et al., 2016). The spatial tiling of brain regions provided by this principal gradient might therefore capture sensory-to-abstract transitions during language processing. Previous fMRI studies could link this gradient to effects of coherency and sentence-level characteristics in speech comprehension and production (Morales et al., 2022; Wu et al., 2022). Further studies have demonstrated the significance of this gradient for semantic processing: Individual differences in semantic performance are linked to variation in gradient connectivity patterns (Gonzalez Alam et al., 2021; Shao et al., 2022). Furthermore, fMRI reveals stronger involvement of the heteromodal end of the gradient for semantic decisions about word pairs that share more semantic features (Wang et al. 2020), as well as for stronger semantic associations (Gao et al., 2022). Thus, consistent with meta-analytic decoding, the gradient dissociates between semantic relations that are more novel or unexpected, compared to well-known and strongly instantiated in memory.

Although this gradient has been linked to the organization of semantic cognition, its recruitment for word representations beyond the semantic level, as well as in sentence reading, which requires continuous integration of incoming information with a linguistic context, has not been explored. The temporal dynamics of functional recruitment along the principal gradient in language processing have also barely been investigated. To address these gaps, we investigated whether the principal gradient aligns with different stages of language processing based on a publicly available neuroimaging dataset of 102 participants during sentence reading (Schoffelen et al. 2019, Arana et al. 2020). This multimodal dataset enabled us to assess the association of the principal gradient to linguistic processing in both fMRI data with high spatial resolution, and in time-sensitive magnetoencephalographic (MEG) recordings. fMRI is mainly sensitive to effects that are robust across time, including both synchronized and desynchronized neural activation that may not be time- and phase-locked to the onset of processing (e.g., Singh, 2012). Evoked responses in MEG, in contrast, are able to investigate how language processing and its relation to the gradient changes across time, while reflecting synchronized neural activation only (e.g., Singh, 2012). By including both of these metrics, we can establish the extent to which systematic variation in the response to sentences along the principal gradient is robust across imaging modalities. Evoked MEG and fMRI are often sensitive to different aspects of the brain’s response (e.g., Babajani et al., 2005; Hall et al., 2014; Singh, 2012), potentially inverting the effects of linguistic variables (Vartiainen et al., 2011). A linguistic parameter may elicit *stronger* responses in brain regions supporting particular language representations because these representations are more easily accessed, or *weaker* brain responses because processing is faster and easier; for example, this pattern has been found for word frequency (e.g., Embick et al., 2001; Faisca et al., 2019; Hauk et al., 2008; Schuster et al., 2016; Yarkoni et al., 2008b). In both situations, however, the effect of the linguistic variable might be located at a similar position on the principal gradient - and at a different position from the effects of other linguistic variables - if the gradient underpins the neural organization of language.

We explored whether the principal gradient aligns with the brain’s response to the psycholinguistic characteristics of the sentences’ individual words, which differentially load on different representational levels, as well as contextual measures reflecting integrative processes as each sentence unfolded. Word-level characteristics included visual properties (word length), orthographic properties (orthographic distance, reflecting familiarity with orthographic subunits), and word frequency (reflecting familiarity with the lexical form of the word). Contextual measures included semantic similarity, reflecting the semantic ‘fit’ of each word with the preceding sentence context, and the position of each word in the sentence, reflecting the amount of contextual information available. These parameters influence reading times (e.g., Pynte et al., 2008a; Hawelka et al., 2013; Dufau et al., 2015; Eisenhauer et al., 2022) and modulate brain activation in various language-relevant regions (e.g., Frank and Willems, 2017; Gagl et al., 2022; Hauk et al., 2008; Schuster et al., 2016; 2020; Yarkoni et al., 2008b) and at various points in time, including earlier and later time windows often associated with visual/orthographic and lexical-semantic and integrative processing (e.g., Dambacher et al., 2006; Dufau et al., 2015; Eisenhauer et al., 2022; Frank and Willems, 2017; Hauk et al., 2006; Laszlo and Federmeier, 2014; Yan and Jaeger, 2020). By investigating each parameter’s brain response along the gradient across time, the present study enables us to uncover the potential relevance of the principal gradient for the organization of language processing through time and across imaging modalities. This will extend our current understanding of the neurobiology of language by (i) moving beyond investigations of the functional contributions of discrete regions to show the relevance of graded whole-brain activation patterns to the emergence of linguistic effects, (ii) placing previously established process-region associations in the broader macroscale organization of the cortex, and (iii) offering an explicitly quantifiable hierarchy of the brain network for language that might inform future computational modelling approaches.

## 2 Methods

### 2.1 MRI and MEG data

We used publicly available MRI and MEG data from the open MOUS (‘mother of unification studies’) dataset (Schoffelen et al., 2019). We focused our investigation on the data from 102 Dutch participants who performed the visual version of the task in an fMRI as well as in an MEG experiment. Further data used in the present study include anatomical T1-weighted MRI data as well as resting state MRI data. During the fMRI and MEG tasks, participants read Dutch sentences, half of which contained a complex relative clause. In addition, participants read scrambled word list versions of the sentences. Out of a total of 360 sentences and their corresponding word lists, each participant was presented with 180 sentences and 180 non-corresponding word lists. 60 sentences and word lists each were presented in the fMRI task, while the remaining 120 sentences and word lists each were presented in the MEG task. The sentences and word lists consisted of nine to 15 words with variable word presentation durations (range: 160 to 1400 ms for fMRI; 300 to 1400 ms for MEG) depending on word length (Schoffelen et al., 2019). Following 20% of the sentences and word lists, a yes/no question on the content of the sentence or the wording of the sentence/word list was presented. We excluded three participants from fMRI analyses, given their data was accidentally recorded with a different phase encoding direction (Schoffelen et al., 2019), which resulted in limited coverage of the temporal cortex. MEG analysis was based on a publicly available already preprocessed version of the MEG data from Arana et al. (2020), who had excluded two participants due to technical issues. Data were collected by the original researchers under approval of the local ethics committee (CMO – the local “Committee on Research Involving Human Subjects” in the Arnhem-Nijmegen region) and following guidelines of the Helsinki declaration.

### 2.2 MRI preprocessing

MRI data were preprocessed using FMRIB’s Software Library (FSL) version 5.0.11. Preprocessing of the task fMRI data involved motion correction using MCFLIRT (Jenkinson et al., 2002), slice-timing correction, brain extraction, spatial smoothing with a six mm full-width-half-maximum (FWHM) Gaussian kernel and high-pass filtering at 100 s. Linear registration of the fMRI data was performed using FLIRT (Jenkinson and Smith, 2001; Jenkinson et al., 2002). Due to the partial field of view which might limit registration accuracy, the task fMRI data was first registered with six degrees of freedom to a brain-extracted slice from the resting state fMRI data which had full field of view. This step was omitted for two participants with missing resting state data. Subsequently, the task fMRI data was registered to the extracted T1-weighted anatomical brain images via linear boundary-based registration, which were registered to the MNI152 standard space with twelve degrees of freedom.

### 2.3 MEG preprocessing and source localization

MEG preprocessing steps previously performed by Arana et al. (2020) involved band-pass filtering the data between 0.5 and 20 Hz to increase signal-to-noise ratio, visually identifying samples contaminated with eye movement, muscle, or sensor jump artifacts based on the variance within each sensor and marking them as missing data to exclude them from analyses, and down-sampling the data to 120 Hz. We performed all subsequent analyses using FieldTrip, (version 2021 11-21; http://fieldtrip.fcdonders.nl; Oostenveld et al., 2011) under MATLAB (version 2019a, The MathWorks Inc.). We subtracted the mean signal during the baseline period from −500 to −50 ms prior to sentence onset from the sentence epochs before arranging the trials into 600 ms epochs aligned to the onset of each individual word.

Source localization partly followed the procedure of Arana et al. (2020), using their pre-computed forward models as well as analysis scripts (https://data.donders.ru.nl/collections/di/dccn/DSC_3011020.09_410?3). The data covariance matrix was estimated using all trials and time points from the sentence epochs and regularized using Lavrentiev regularization (Lavrentiev, 1967) with a regularization parameter of 100%, i.e., the unscaled mean was added to the covariance matrix (Arana et al., 2020). The forward models were based on individual participants’ structural magnetic resonance images coregistered to a cortical surface with 8196 vertices based on the Conte69 atlas (Van Essen et al., 2011). For each vertex, linearly constrained minimum variance (LCMV) spatial filters (Van Veen et al., 1997) were computed based on all trials and time points from the sentence epochs and normalized using unit-noise gain. Spatial filters were computed based on the optimal dipole orientation that maximized spatial filter output, which was identified using singular value decomposition. Single-trial source power based on the precomputed spatial filters was then estimated for five consecutive 100 ms time windows of each individual word. Source power was averaged across vertices for each of 400 surface parcels from the functional connectivity-based Schaefer parcellation (Schaefer et al., 2018). Fifty parcels along the midline were excluded from further analyses due to the relative insensitivity of MEG to deep neural sources (e.g., Hillebrand & Barnes, 2002; Piastra et al., 2021).

### 2.4 Analysis of the association of word and sentence level parameters with the principal gradient in fMRI and MEG data

#### 2.4.1. General approach

To assess the potential involvement of the principal connectivity gradient (Margulies et al., 2019) in different processing stages during reading, we used a three-step approach. First, we determined how strongly cortical surface regions were modulated by multiple word parameters, reflecting different stages of word processing, as well as contextual parameters, reflecting the integration of meaning across the whole sentence. We used general linear models (GLMs) for fMRI data and linear mixed models (LMMs) for MEG data to estimate the effects of each of these parameters on brain activation. This resulted in cortical maps of the effect strength for each parameter. In the second step, we asked whether the strength of each linguistic effect covaried with the organization of the brain captured by the principal gradient (Margulies et al., 2016), using the maps from BrainSpace (Vos De Wael et al., 2020). We assessed the dependence of the GLM or LMM estimates of language variables on gradient values across all cortical regions, using linear models. Parameters closely tied to visual information in the input, such as word length, are expected to modulate the sensory end of the gradient to a greater extent than the heteromodal end, which would be evident from a gradual decline of word length estimates with increasing principal gradient values. On the other hand, parameters that reflect more abstract information, such as the semantic similarity between a word and the previous context, should involve the heteromodal end of the gradient to a greater extent than the sensory end, and this would be reflected by a gradual increase of semantic similarity estimates with increasing principal gradient values. While previous studies used similar linear regression approaches in the same dataset (Huizeling et al., 2022; King et al., 2020), our investigation of the association with the principal gradient is novel. As a third step, we assessed if a parameter’s significant association with the gradient observed in the previous step follows a gradual or discrete pattern by comparing if the association with the gradient shows a better fit to the linear function or to a step function with three steps, based on Akaike information criterion (AIC) as well as Bayesian information criterion (BIC).

We also investigated the associations of the parameters of interest with Gradients 2 and 3, i.e., the gradients which explained the second and third most variance in resting state fMRI data (Margulies et al., 2016; Vos De Wael et al., 2020). Gradient 2 dissociates visual from motor cortex, while Gradient 3 dissociates the default mode network from the multiple demand network (Supplementary Fig. S3). In addition, we investigated quadratic associations with the principal gradient, which might partly capture similar effects as Gradient 3, which tends to assign higher values to brain regions located in the middle of the principal gradient and correlates with the squared values of the principal gradient at an r of .68. Results for gradients 2 and 3 as well as quadratic associations with the principal gradient are included in the Supplement.

#### 2.4.2 Word and contextual parameters of interest

Parameters of interest were chosen to reflect different processing stages of word recognition (cf. Dufau et al., 2015; Eisenhauer et al., 2022): At the individual word level, *word length*, i.e., the number of letters, primarily reflected visual processing, *Orthographic Levenshtein Distance 20 (OLD20)* reflected orthographic familiarity (Yarkoni et al., 2008a), while *word frequency* reflected lexical familiarity. At the contextual level, we investigated the *semantic similarity* between a word and the preceding sentence context, as well as the *position* of each word in a sentence, reflecting the amount of contextual information available. OLD20 was estimated based on Subtlex-NL (Keuleers et al., 2010) using the R package vwr (Keuleers, 2013). Word frequency was obtained from the Subtlex-NL database and log-transformed per million. The semantic similarity between each word and the preceding sentence context was based on a computational language model trained on a large text corpus that represents words and phrases as vectors dependent on the context they appear in (e.g., Mikolov et al., 2013). Words which appear in similar contexts have more similar vector representations, so that the distance between the vectors of two words reflects their semantic similarity. In the present study, semantic similarity corresponded to the cosine similarity between a word’s vector representation and the vector representation of the preceding five-word phrase (cf. De Boom et al., 2015; Wang et al., 2014). The vector representations were estimated based on the ELMo (‘Embeddings from Language Models’) language model (Peters et al., 2018), using the pretrained Dutch model from ‘ELMo for many languages’ (Che et al., 2018; hosted at the NLPL Vectors Repository; Fares et al., 2017). ELMo is a contextualized model, i.e., vector representations are not static but can be adjusted based on the context a word appears in (Peters et al., 2018). Accordingly, the five preceding words were taken into account as context when estimating each word’s vector representation. As the estimation of semantic similarity was based on the context from five preceding words, it was not possible to obtain semantic similarity for the first five words from the sentences. To allow for equivalent data support for all parameters, we only included words starting from position six in the investigation of all parameters. We investigated the effects of the parameters of interest only for content words in sentences, i.e., excluding function words or other types of words such as names, as well as the word lists. This resulted in an average of 240 words per participant (range: 230 to 249) being included in the analysis. See Supplementary Figure S1 for parameter correlations.

#### 2.4.3 fMRI analysis

##### General linear modelling

For each participant and voxel, we estimated a GLM including explanatory variables (EVs) at the individual word or sentence/word list level using FSL, version 5.0.11. As the presentation duration of the words was confounded with word length, as well as being relatively short for an fMRI study such that linguistic processing will likely last beyond this presentation duration, word-level EVs were modelled from the onset of the respective word with a fixed duration of 1 s. This procedure is in line with a previous study using the same dataset (King et al., 2020). Our conclusions from this model were confirmed in a supplementary analysis that used presentation time as the duration (see Supplementary Fig. S7). The main word-level EVs of interest for this study, i.e., *word length, OLD20, word frequency, semantic similarity* and *position* were only included for content words in sentences. Given how semantic similarity was estimated (see section 2.4.2), all parameters of interest were only included starting from the sixth word in each sentence. Part of speech was included as an additional word-level EV for both sentences and word lists, i.e., *adjective, noun, verb,* or *other type of word* (comprising all remaining words such as function words and names). EVs at the sentence/word list level were modelled from the onset and duration of the respective sentence/word list: these included *sentence with a complex relative clause, sentence without a complex relative clause, word list,* and *presentation order*. A final EV comprised the *fixation time window (before each sentence/ word list) plus cues* (indicating if the next block would consist of sentences or word lists). Two second blank periods between the sentences/word lists provided an implicit baseline. All parametric EVs were centred and scaled. EV time courses were convolved with a hemodynamic response function and their temporal derivatives were included in the GLM. Variance inflation factors were below ten for all EVs of interest and their derivatives. To account for potential motion artefacts, the GLM also included a total of 24 standard and extended motion parameters, as well as voxels that were motion outliers based on framewise displacement.

At the group level, we used GLMs to obtain two-tailed z values for the effects of word length, OLD20, word frequency, semantic similarity and position at each voxel. We converted the z value maps for these contrasts from volumetric to surface space (FreeSurfer fsaverage5) using the vol_2_surf function from the python package nilearn (https://nilearn.github.io/stable/index.html). As interpolation to the surface induced noise by reducing the numerical values of z values at the borders of the field of view, we excluded surface vertices if z values were diminished by more than 25%. These vertices were identified based on a surface interpolation of a binary volumetric mask that had a value of one for voxels included in the GLM analysis and a zero for all other voxels. Vertices were excluded if they had a value below .75 after interpolation of the mask to the surface. Z values of the remaining surface vertices were then averaged for each of the 400 surface parcels from the functional connectivity-based Schaefer parcellation (Schaefer et al., 2018). We included only parcels for which at least 25% of the corresponding surface vertices were available given the limited field of view of the fMRI data, resulting in 269 parcels used for further analysis.

##### Associations of word and contextual parameters with the principal gradient in fMRI data

For each linguistic parameter of interest, we used linear models in R (version 3.5.2; 2018-12-20; R Development Core Team, 2008) to compute the dependence of the GLM z values on the principal gradient across the whole brain (Margulies et al., 2016; Vos De Wael et al., 2020), i.e., across 269 Schaefer parcels. Given the left-hemisphere dominance of language processing (e.g., Binder et al., 2009), which may also contribute to hemispheric differences in the gradient’s contribution to semantic cognition (Gonzalez Alam et al., 2022), we included hemisphere in an interaction term with gradient in the linear models. For main effects of the gradient or interactions with hemisphere which had a p-value < .05 in the linear models, we assessed statistical significance controlling for spatial covariance and type one error rate using a spin permutation procedure (Alexander-Bloch et al., 2018). The permutation procedure relied on 1000 permuted versions of the principal gradient which were constructed in BrainSpace. A parameter’s dependence on the gradient (as a main effect or in an interaction with hemisphere) was considered significant if the actual t-value exceeded the 95th percentile of the t-value distribution from the permuted gradients.

For each parameter showing a significant linear association with the gradient, we investigated if the association with the gradient shows a better fit to the linear function or to an alternative three-step function. This analysis aimed at dissociating if the brain response to the parameter changes in a continuous or discrete manner along the gradient. As the original linear model, the step function included interactions with hemisphere. Goodness of fit was determined using both AIC and BIC. AIC or BIC differences greater than four are considered to provide less support for the model with higher AIC/BIC (e.g., Burnham and Anderson, 2004; Raftery, 1996). If the difference in AIC as well as BIC of the step model minus the linear model was greater than four, we concluded that the linear model was favoured, whereas for a difference smaller than minus four, we concluded that the step model was favoured. For AIC or BIC differences between −4 and 4, no model was clearly favoured over the other. In case the linear model was not clearly favoured, we confirmed if the association with the step model was still significant when using the spin permutation procedure described above.

##### Dissociating effects of word and contextual parameters along the gradient in fMRI data

In order to investigate whether word and contextual parameters of interest fell at distinct positions along the principal gradient, we assessed whether the linear association with the gradient differed significantly between parameters and hemispheres. First, an analysis of variance (ANOVA) examined the interaction between gradient value, parameter (five levels corresponding to five linguistic/context effects) and hemisphere, taking z values from the group level GLM as the dependent variable. A positive relationship between a parameter and the gradient can either indicate that a positive effect of the parameter is increasing towards the heteromodal end of the gradient, and/or that a negative effect of the parameter is decreasing towards the heteromodal end of the gradient. To facilitate comparability of the relationship with the gradient between parameters, for this analysis, we inverted all z values of a parameter if they were negative for the majority of parcels. As a result, positive associations with the gradient always corresponded to a stronger effect of the language variable at the heteromodal end of the gradient, whereas negative associations with the gradient always corresponded to a stronger effect at the sensory end of the gradient. For significant gradient by parameter interactions, follow-up analyses tested for significant differences between pairs of linguistic parameters, computing the interaction between gradient and parameter (two levels) using linear models. For significant three-way interactions (for gradient by parameter by hemisphere), the follow-up analyses were performed separately for each hemisphere. Again, for all effects with a p-value < .05 in the ANOVAs/linear models, we assessed statistical significance using a spin permutation procedure (Alexander-Bloch et al., 2018) based on 1000 permuted versions of the principal gradient. Effects were considered significant if the actual F/t-value exceeded the 95th percentile of the F/t-value distribution from the permuted gradients.

#### 2.4.4 MEG analysis

##### Linear mixed modelling

As the high temporal resolution of MEG data allowed us to assess the temporal dynamics of gradient associations, we investigated five consecutive 100 ms time windows, starting from the onset of each word up until 500 ms. In Step 1 of our analysis, linear mixed models (LMMs) estimated the effects of multiple word and contextual parameters on source activation for each of 400 Schaefer parcels and each of the five time windows, using the lmerTest package (Kuznetsova et al., 2017) of the statistical software package R, version 3.5.2 (2018-12-20; R Development Core Team, 2008). Only content words of the sentences were included in the analysis. In addition, the first five words were excluded from analysis based on how the semantic similarity parameter was estimated (see section 2.4.2). Thus, our analysis focuses on word reading after an initial sentence context has been established. On average, 409 words per participant and time window (range: 241 to 478) were included in the analysis.

We included the same word and contextual parameters of interest as in the fMRI data, i.e., *word length, OLD20, word frequency, semantic similarity* and *position.* Additionally, the LMMs included the following fixed effects of no interest: *part of speech* (i.e., noun, adjective or verb), *syntactic complexity* (sentences containing a more complex relative clause vs. sentences of lower syntactic complexity), and *sentence presentation order*. All numeric fixed effects were centred and scaled. Initially, p*articipant* and *item* were included as random effects on the intercept. However, this resulted in a high proportion of overfitted models (as evident from ‘singular fit’ warning messages), and we consequently excluded the random effect of item (cf. Eisenhauer et al., 2022) based on the rationale of parsimonious mixed modelling (Bates et al., 2018), which resolved this issue for all models. For each time window and parcel, the LMM estimates of each parameter were extracted for the subsequent investigation of a potential association with the principal gradient.

##### Associations of word and contextual parameters with the principal gradient in MEG data

For each of the investigated parameters of interest and for each time window, we used linear models to compute the dependence of the LMM estimates on the principal functional connectivity gradient (Margulies et al., 2016) obtained from the BrainSpace toolbox (Vos De Wael et al., 2020) across the whole brain (350 Schaefer parcels). The linear models included the interaction of gradient with hemisphere. For main effects of the gradient or interactions with hemisphere which had a *p*-value < .05 in the linear models, we assessed statistical significance using a spin permutation procedure (Alexander-Bloch et al., 2018) as described in section 2.4.3. Associations between language parameters and gradient values (as a main effect or in an interaction with hemisphere) were considered to be significant if the actual *t*-value exceeded the 95^th^ percentile of the *t*-value distribution from the permuted gradients.

For each significant linear association with the gradient, we investigated if the association with the gradient shows a better fit to the linear function or to alternative step functions with two or three steps. As the original linear model, step functions included interactions with hemisphere. If absolute AIC and BIC differences between the two models were greater than four, we favoured the model with the lower AIC/BIC (cf. Burnham and Anderson, 2004; Raftery, 1996). For AIC or BIC differences between −4 and 4, no model was clearly favoured over the other. In case the step model was favoured, we confirmed if the association was still significant when using the spin permutation procedure described above.

##### Dissociating effects of word and contextual parameters along the gradient in MEG data

In order to investigate whether distinct gradient locations were associated with the different word and contextual parameters, for each time window, an ANOVA assessed the three-way interaction between parameter (five levels), gradient value and hemisphere on the parameter’s estimates on brain activation as obtained from LMMs. If a parameter’s LMM estimates were negative for the majority of parcels at a certain time window, these estimates were inverted for this analysis. This ensured that positive associations with the gradient always corresponded to a stronger effect at the heteromodal end of the gradient, whereas negative associations with the gradient corresponded to a stronger effect at the sensory end of the gradient. When the gradient by parameter interaction was significant, follow-up linear models investigated the interaction between gradient value and parameter (two levels) for each pair of parameters. For significant three-way interactions with hemisphere, these follow-up tests were performed separately for each hemisphere.

In addition, for each parameter, we investigated if the association with the gradient changes across time. First, we used an ANOVA to investigate the three-way interaction between gradient value, time window (five levels) and hemisphere on the estimates from LMMs as dependent variable. As the amplitude of event-related magnetic fields develops non-linearly across time, time window was entered as a discrete rather than a continuous factor. Again, predominantly negative LMM estimates were inverted. For significant gradient by time window interactions, follow-up models investigated the interaction between gradient value and time window (two levels) for each pair of time windows. For three-way interactions with hemisphere, these follow-up tests were performed separately for each hemisphere.

For all effects with a p-value < .05 in the ANOVAs/linear models, we assessed statistical significance using a spin permutation procedure (Alexander-Bloch et al., 2018) controlling for spatial correlation and type one error rate as described in section 2.4.3.

#### 2.4.5 Control analyses

The presentation duration of the words was varied according to word length. Due to the high correlation between word length and presentation duration (r = .88), we refrained from including both parameters in the GLMs/LMMs simultaneously. To control for potential influences from presentation duration on the other linguistic parameters, we re-estimated the GLMs/LMMs after replacing word length with presentation duration. For all parameters except word length, we only interpreted a parameter’s association with the gradient if it reached significance based on the estimates from *both* GLM/LMM versions, i.e., when including word length *and* when including presentation duration. Differences in gradient location between these parameters were only considered significant when the effects were found in both analyses. In case of the fMRI GLM, we also performed an additional analysis that included word length and the other four parameters of interest and control parameters while modelling the duration of each word level EV as the actual word presentation duration instead of a fixed 1s duration. While this analysis can resolve the confound between presentation duration and word length, this comes at the disadvantage that the model will likely underestimate the duration for which each linguistic parameter influences the BOLD response.

### 2.5 Behavioural data and analysis

To assess the effects of our parameters of interest on reading behaviour, we used publicly available data from a self-paced reading study including a subset of 160 sentences from the MOUS dataset in a separate sample of 73 participants (Kapteijns and Hintz, 2021). Data collection was approved by the ethics board of the Faculty of Social Sciences at Radboud University. In this study, sentences were presented word by word until button press. 20% of the sentences were followed by a yes/no question about the content or wording of the sentence, and two participants were excluded based on a performance threshold of 80% accuracy (Kapteijns and Hintz, 2021). We used LMMs to assess the effects of each of our parameters of interest, i.e., word length, OLD20, word frequency, semantic similarity, and position, on log-transformed self-paced reading response times. The effects of word length, word frequency, and position on reading times have been investigated previously in this dataset (Kapteijns and Hintz, 2021), while the effects of OLD20 and semantic similarity have not. Again, we included part of speech, syntactic complexity, and sentence order as additional control variables. All continuous variables were centred and scaled. Participant and item were included as random effects on the intercept. The step function from the lmerTest package in R was used to reduce fixed effects from the original LMMs in a stepwise procedure based on Akaike information criterion and an alpha threshold of .05, resulting in a model most optimally explaining the data. All fixed effects remaining in the optimal model were considered significant.

## 3 Results

### 3.1 Behavioural results

First, behavioural data from a self-paced reading study indicated the effects of the parameters of interest on reading speed (see Supplementary Table S1 for detailed statistics). Self-paced reading response times were significantly modulated by word length, OLD20, and position (Fig 1). Response times were faster for shorter words, orthographically more familiar words (i.e., words of lower OLD20), and words occurring earlier in the sentence.

**Figure 1.**
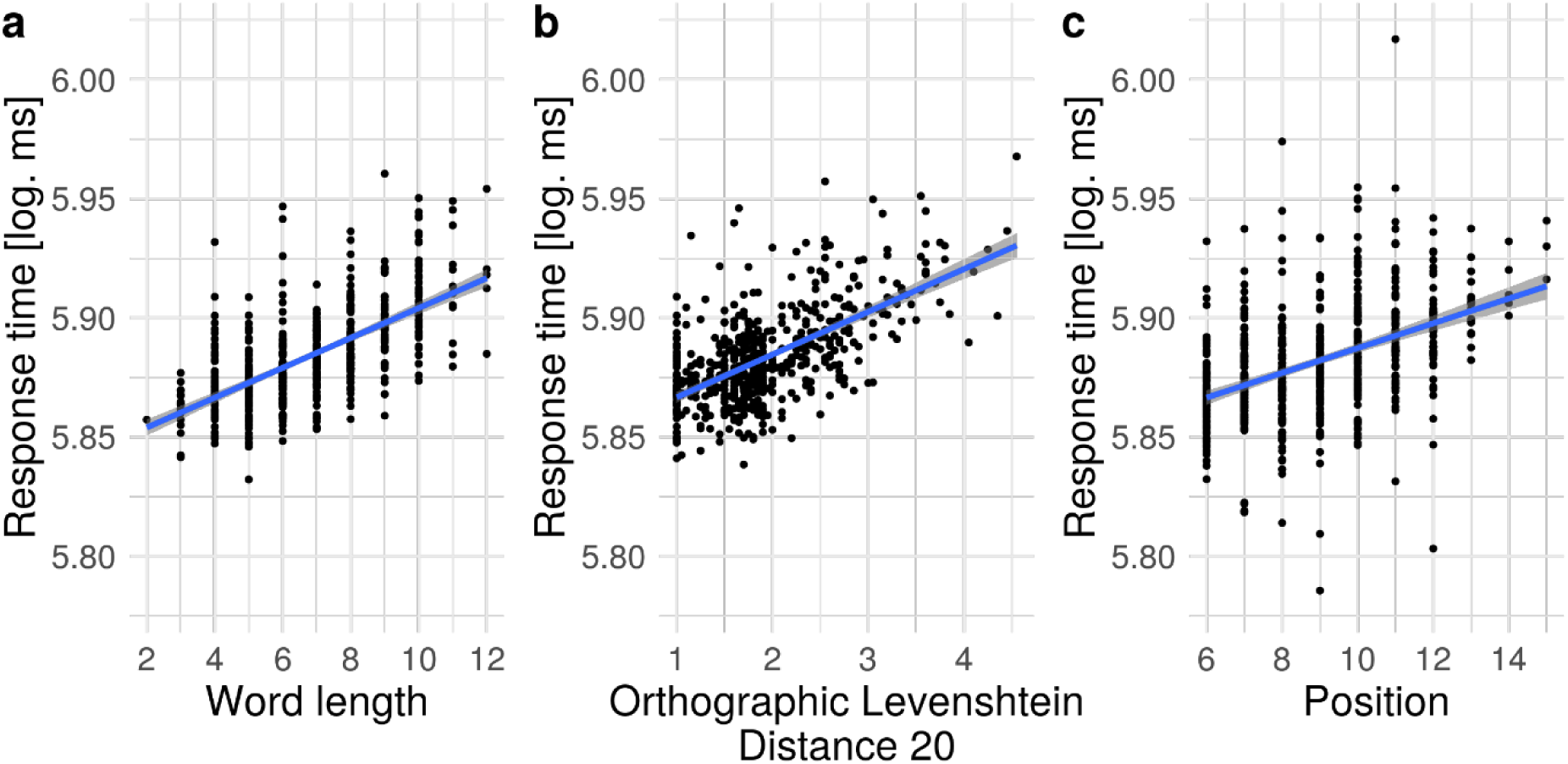
Significant effects of a) word length, b) orthographic Levenshtein distance 20, and c) position on log-transformed self-paced reading response times. Partial effects from linear mixed models are shown. Each dot represents a word averaged across participants.

### 3.2 fMRI results

#### 3.2.1 Associations of word and sentence level parameters with the principal gradient in fMRI data

Having established the behavioural effects of our parameters of interest on reading speed, we investigated the linear association of the different word and contextual parameters with the principal functional connectivity gradient in fMRI BOLD activation. Significant effects of each parameter on brain activation irrespective of the principal gradient are shown in Supplementary Figure S2. Significant linear associations of the parameters’ effects with gradients 2 and 3 are shown in Supplementary Figures S4 and S5, while results for quadratic associations with the principal gradient are shown in Supplementary Figure S6. Due to the variable presentation duration of the words, which correlated strongly with word length, we performed an additional control analysis that estimated the effect of presentation duration, rather than word length, as well as the other parameters’ effects on brain activation. For all parameters except length, we will report only those associations with the principal gradient that also reached significance when presentation duration was controlled for (see Supplementary Tables S2 and S3 for detailed results of both GLM versions). In addition, results when modelling the actual presentation duration of the words instead of a fixed duration are shown in Supplementary Figure S7. Linear models were used to investigate the relationship between a parameter’s effect on brain activation and the principal gradient. For effects with a p-value < .05 in linear models, a spin permutation procedure was used to assess if the association with the gradient was significant in comparison to permuted versions of the principal gradient; significant effects based on spin permutation are described in the following.

Significant relationships with the gradient (Fig. 2a) based on whole-brain fMRI BOLD activation effects were found for three investigated word parameters, i.e., word length (p_spin_ = .008), OLD20 (p_spin_ < .002) and word frequency (p_spin_ = .006; Fig. 2b-d). For the majority of parcels, BOLD activation increased with decreasing word length, increasing OLD20 (i.e., decreasing orthographic familiarity), and decreasing word frequency. Absolute effects of each of these three parameters were stronger at the sensory as opposed to the heteromodal end of the gradient, indicated by a significant increase (from stronger negative values to values around zero, for word length and word frequency) or decrease (from stronger positive values to values around zero, for OLD20) of z values with increasing gradient values. No significant interactions with hemisphere were found. Contextual effects, i.e., semantic similarity and position, did not differ significantly along the principal gradient (Fig. 2e,f). A model comparison contrasting the significant linear models with a three-step function indicated that the linear model was favoured for OLD20 (difference in score for step minus linear model: AIC = 6.1; BIC = 13.3); however, the step function was favoured for word length (difference in score for step minus linear model: AIC = −29.9; BIC = −22.7, p_spin_ first vs. second step: .24; first vs. third step: < .002, second vs. third step: < .002) and word frequency (difference in score for step minus linear model: AIC = −13.6; BIC = −6.4, p_spin_ first vs. second step: .50; first vs. third step: < .002, second vs. third step: < .002), indicating that negative word length and word frequency effects are relatively weaker at the heteromodal end of the principal gradient in comparison to intermediate and unimodal gradient locations (Supplementary Fig. S8). Significant quadratic associations with the gradient for word length and word frequency also indicated relatively stronger negative effects at an intermediate gradient location (Supplementary Fig. S6).

**Figure 2.**
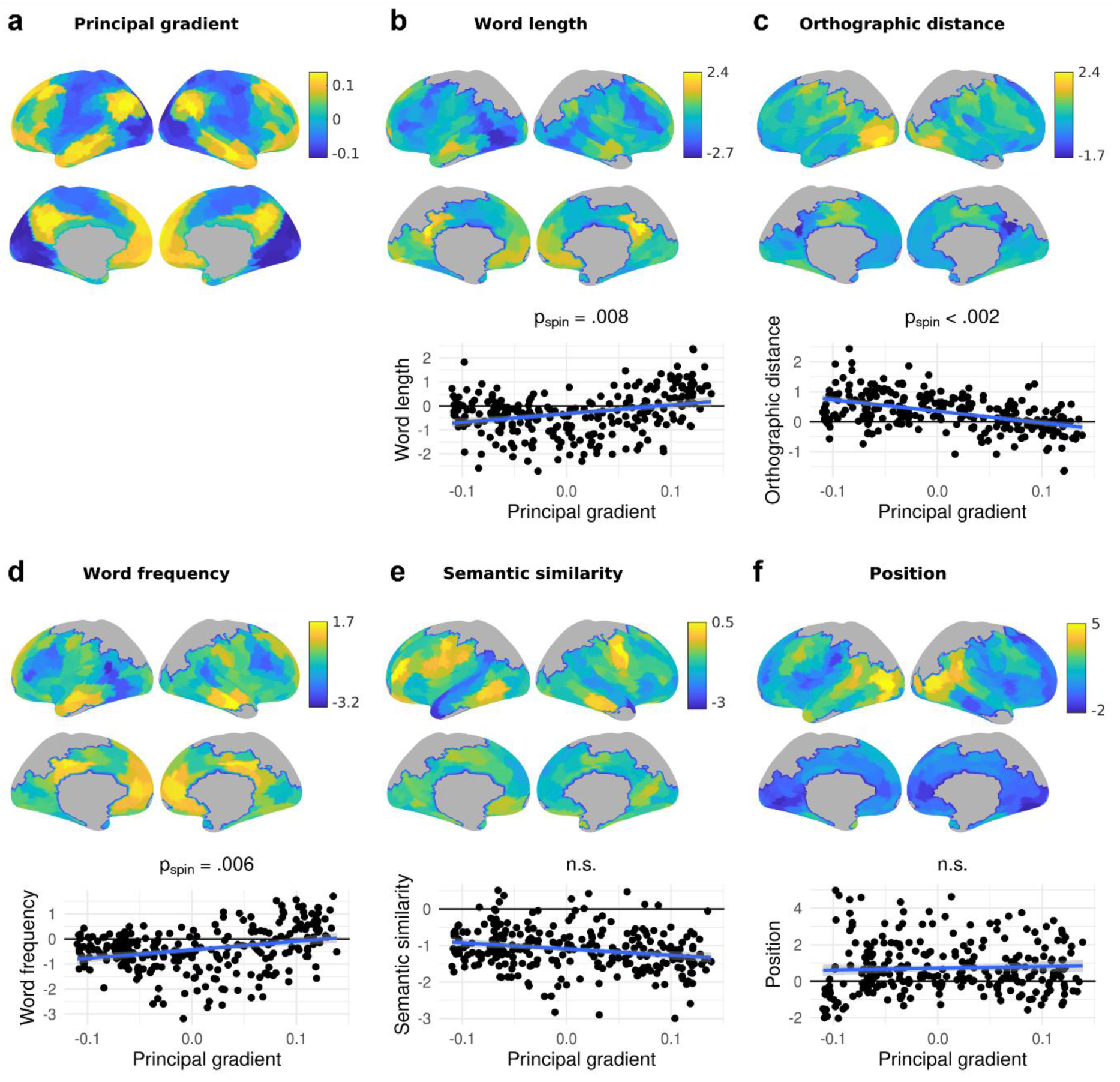
The principal connectivity gradient (a) as well as parameter effects on brain activation (b-f; top: lateral view, middle: medial view) and their dependence on the principal gradient (bottom) across the whole brain. In b-f, the top and middle panels show the effect of the respective word parameter on fMRI BOLD activation across 269 cortical parcels, indicated by the z values from the group-level general linear model. Positive z values indicate an increase in brain activation with increasing parameter values, while negative z values indicate a decrease in brain activation with increasing parameter values. Grey parcels were not analysed due to limited field of view during data acquisition. The bottom panels show the dependence of these effects on the principal gradient across parcels, i.e., the z values are plotted against the principal gradient values from Margulies et al. (2016). The lowest gradient values are associated with sensory cortices, while the highest gradient values are associated with heteromodal cortices. p_spin_ indicates p values from spin permutation; n.s. = not significant.

#### 3.2.2 Gradient location of word and contextual parameters’ effects in fMRI data

In order to assess whether principal gradient location dissociates between the different word and contextual parameters, we computed the interaction between gradient value, parameter, and hemisphere with the parameter’s effect on brain activation as dependent variable, indicated by each parcel’s z value from group level general linear models. Respective results for Gradients 2 and 3 are shown in Supplementary Figures S9 and S10. For all parameters except word length, we will only consider those effects as significant that also reached significance when presentation duration was controlled for.

We found a significant interaction between gradient value and parameter that was robust using spin permutation (p_spin_ < .001), which did not significantly interact with hemisphere (Supplementary Tables S4 and S5). Post hoc tests for pairwise differences between parameters based on spin permutation revealed that semantic similarity as well as position effects show a significantly different association with the gradient in comparison to word length (semantic similarity: p_spin_ < .002; position: p_spin_ = .002), OLD20 (semantic similarity: p_spin_ < .002; position: p_spin_ = .01) and word frequency effects (semantic similarity: p_spin_ < .002; position: p_spin_ < .002). This was driven by negative associations with the gradient for word length, OLD20, and word frequency, in contrast to a positive association for semantic similarity and position (Fig. 3a).

**Figure 3.**
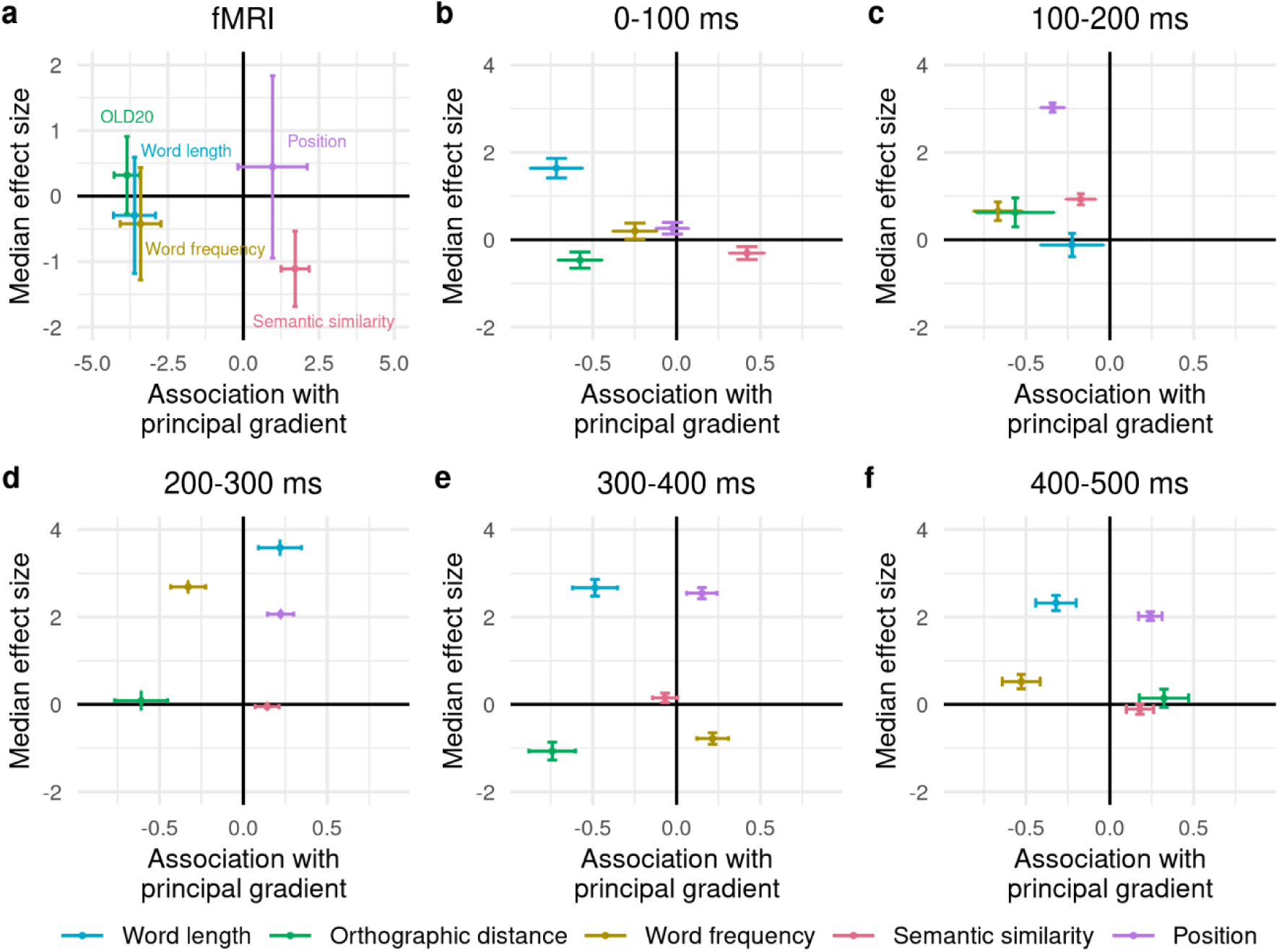
Association (i.e., linear model estimate) of each parameter with the connectivity gradient as indicated on the x axis in a) fMRI BOLD activation and b-f) MEG-measured brain activation across five time windows. Negative values on the x axis indicate a stronger effect of the parameter towards the sensory end of the gradient, while positive values indicate a stronger effect towards the heteromodal end of the gradient. Horizontal error bars indicate standard errors from linear models. The median effect size (i.e., general linear model z value in fMRI data; linear mixed model estimate in MEG data) of each parameter on brain activation across parcels is indicated on the y axis. Negative values indicate a negative effect direction (i.e., decreasing brain activation with increasing parameter values) for most parcels, while positive values indicate a positive effect direction for most parcels. Vertical error bars indicate the standard deviation of the parameter’s effect size across parcels.

### 3.3 MEG results

#### 3.3.1 Associations of word and sentence level parameters with the principal gradient in MEG data

As for the fMRI data, we investigated the linear association of the different word and contextual parameters with the principal gradient in MEG-measured brain activation. Significant effects of the parameters on brain activation irrespective of the principal gradient are shown in Supplementary Figure S11. Linear associations of the parameters with Gradients 2 and 3 are shown in Supplementary Figures S4 and S5, while results for quadratic associations with the principal gradient are shown in Supplementary Figure S12. Again, we report only those associations with the principal connectivity gradient that were significant both based on the original LMM estimates, including word length, as well as based on the control analysis including presentation duration (see Supplementary Tables S6 and S7 for detailed results of both LMM versions). The relationship between a parameter’s effect on brain activation and the principal gradient was considered significant if the p-value from linear models was below .05 and the effect also reached significance using spin permutation.

The effects of all investigated parameters on MEG-measured brain activation were significantly associated with the principal gradient (Fig. 4a) in particular time windows. MEG source activation increased with longer word length for most parcels. In the earliest time window from 0 to 100 ms, these effects were particularly strong at the sensory end of the gradient in the left hemisphere, as evident from a decline in word length estimates with an increase in gradient values (gradient by hemisphere interaction: p_spin_ = .02; Fig. 4b).

**Figure 4.**
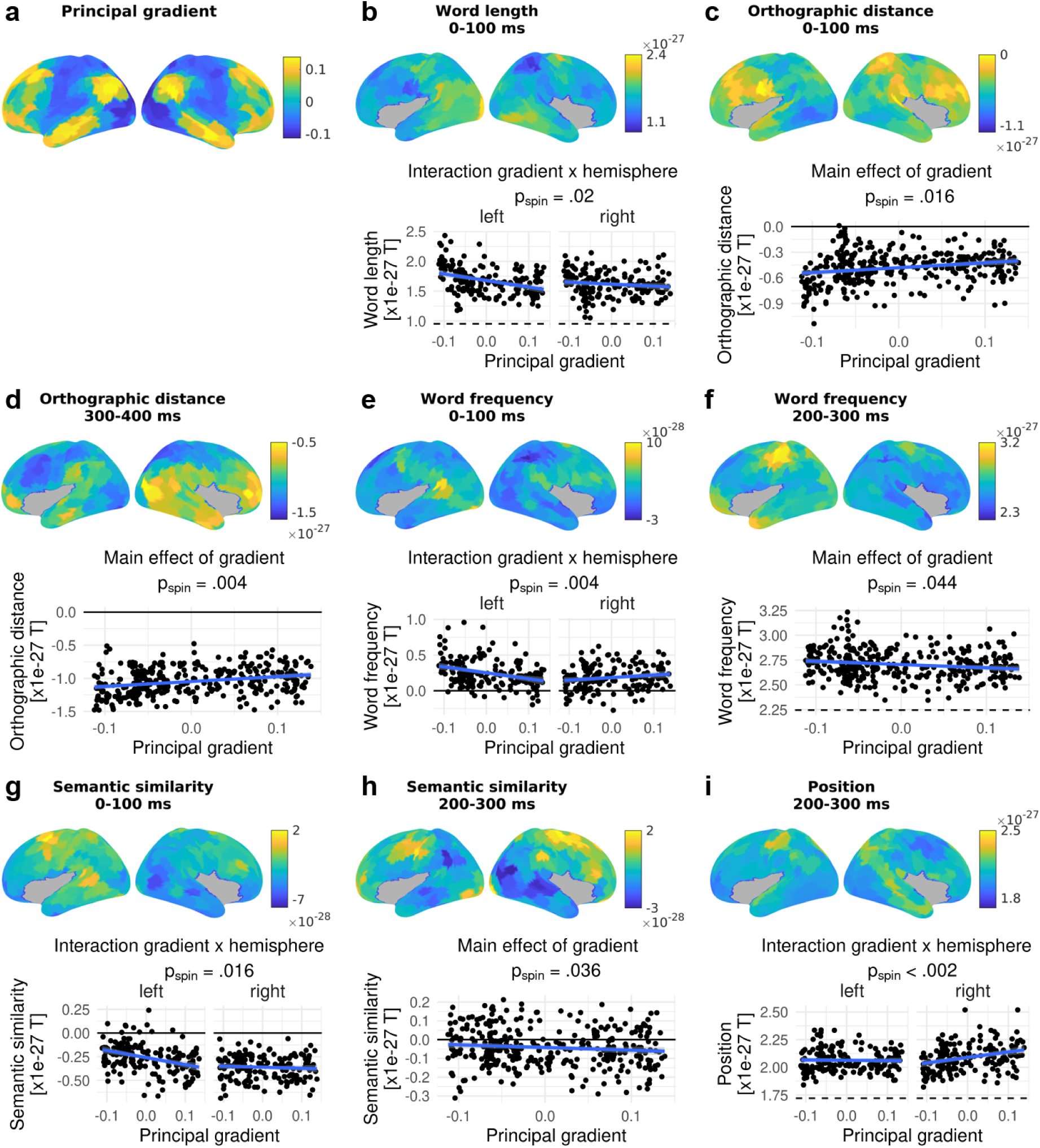
The principal connectivity gradient (a) as well as parameter effects on brain activation (b-i; top) and their dependence on the principal gradient (bottom) across the whole brain. In b-i, the top panels show the effect of the respective word or contextual parameter on MEG source activation across 350 cortical parcels, indicated by the estimates from linear mixed models. Positive estimates indicate an increase in brain activation with increasing parameter values, while negative estimates indicate a decrease in brain activation with increasing parameter values. Grey parcels were not analysed. The bottom panels show the dependence of these effects on the principal gradient across parcels, i.e., the linear mixed model estimates are plotted against the principal gradient values from Margulies et al. (2016). The lowest gradient values are associated with sensory cortices, while the highest gradient values are associated with heteromodal cortices. Only the parameters and time windows which were significantly dependent on the principal gradient are shown. For parameters and time windows that only showed a significant main effect of the gradient, the bottom panels show the association between parameter estimates and gradient values combined across the left and right hemisphere, as in a, b, c, e, and g. In case there was a significant interaction between gradient and hemisphere, the bottom panels show the association between parameter estimates and gradient values separately for the left and right hemisphere, as in d, f, and h. p_spin_ indicates p values from spin permutation for main effects or interactions.

At the orthographic level, source activation decreased with OLD20 at most parcels, i.e., the evoked response increased with orthographic familiarity. This effect was more pronounced at the sensory end of the gradient, resulting in a decrease of the negative OLD20 effect along the gradient from 0 to 100 ms (p_spin_ = .016; Fig. 4c) as well as from 300 to 400 ms (p_spin_ = .004; Fig. 4d).

Concerning word frequency, source activation was stronger for more frequent words in most parcels, in particular at the sensory end of the gradient, resulting in a decrease in word frequency estimates along the gradient. These associations between word frequency and the principal gradient first reached significance in the 0 to 100 ms time window in the left hemisphere (gradient by hemisphere interaction: p_spin_ = .004; Fig. 4e). Further significant associations emerged from 200 to 300 ms bilaterally (p_spin_ = .044; Fig. 4f).

Significant associations of semantic similarity with the gradient occurred from 0 to 100 ms in the left hemisphere (gradient by hemisphere interaction: p_spin_ = .016; Fig. 4g) as well as from 200 to 300 ms bilaterally (p_spin_ = .036; Fig. 4h). In these time windows, source activation was lower when words were more semantically similar with the preceding sentence context in the majority of parcels, and this negative effect increased along the gradient, i.e., absolute semantic similarity effects were stronger at the heteromodal end of the gradient.

Finally, position effects indicated a higher activation for later vs. earlier words in the sentence. These position effects increased along the gradient from 200 to 300 ms in the right hemisphere (gradient by hemisphere interaction: p_spin_ < .002; Fig. 4i), i.e., position effects were stronger at the heteromodal end of the gradient.

For the majority of significant associations with the gradient, model comparisons favoured the original linear models over three-step functions (difference in AIC and BIC scores for step minus linear models > 4). The only exception was the word frequency effect from 200 to 300 ms (Supplementary Fig. S13), for which neither model could be favoured based on AIC (difference in score for step minus linear model: 2.2), while BIC favoured the linear model (difference in score for step minus linear model: 9.9; step model p_spin_ values: first vs. second step: .20, first vs. third step: .044, second vs. third step: .6).

#### 3.3.2 Gradient location of word and contextual parameters’ effects in MEG data

As for fMRI, we investigated whether the word and contextual parameters were associated with significantly different principal gradient locations in the evoked responses measured with MEG. Respective results for Gradients 2 and 3 are shown in Supplementary Figures S9 and S10. Effects were considered significant if this was confirmed when presentation duration was controlled for. ANOVAs examined each parameter’s effect on brain activation (i.e., linear mixed model estimates) as a dependent variable and considered interactions between gradient value, linguistic parameter, and hemisphere for each time window (Supplementary Tables S8 and S9). In each time window, we found significant gradient by parameter interactions that remained robust using spin permutation (0 to 100 ms: p_spin_ = .024; 100 to 200 ms: p_spin_ = .01; 200 to 300 ms: p_spin_ = .048; 300 to 400 ms: p_spin_ = .022; 400 to 500 ms: p_spin_ = .036) and were not further modulated by hemisphere. The significant pairwise parameter differences driving this interaction are described below. The effect direction for most parameters was variable across time windows - effects were negative in some time windows and positive in the other time windows, as indicated on the y axes in Figure 3. Word length effects were mainly positive: brain activation increased with increasing length, except for the 100 to 200 ms time window that showed predominantly negative word length effects across parcels. OLD20 effects were mainly positive: brain activation was weaker with more orthographic familiarity, except for the 0 to 100 and 300 to 400 ms time windows that had predominantly negative OLD20 effects. Word frequency effects were mainly positive, indicating stronger evoked responses for more frequent words; however, the opposite pattern was found in the 300 to 400 ms time window. Semantic similarity effects were mainly negative, indicating weaker brain activation with increasing similarity, although semantic similarity effects were predominantly positive in the time windows 100 to 200 and 300 to 400 ms. Position effects were predominantly positive, i.e., brain activation increased for later words in the sentence.

Significant differences in the association with the gradient were observed between the following parameters using spin permutation: From 0 to 100 ms after word onset, word length (p_spin_ = .004), OLD20 (p_spin_ = .004), and word frequency effects (p_spin_ = .026) showed a negative association with the gradient (stronger effects at the sensory end), in contrast to semantic similarity showing a positive association with the gradient (stronger effects at the heteromodal end; Fig. 3b). From 100 to 200 ms, word frequency showed a more negative association with the gradient than semantic similarity (p_spin_ = .016; Fig. 3c). From 200 to 300 ms, OLD20 and word frequency effects were negatively associated with the gradient in contrast to semantic similarity (OLD20: p_spin_ = .012; word frequency: p_spin_ = .036) and position effects (OLD20: p_spin_ = .004; word frequency: p_spin_ = .014) which were positively associated (Fig. 3d). From 300 to 400 ms, word length and OLD20 were more negatively associated with the gradient than semantic similarity (which was also negatively associated; word length: p_spin_ = .05; OLD20: p_spin_ = .024), word frequency (word length: p_spin_ = .006; OLD20: p_spin_ = .002) and position (word length: p_spin_ = .002; OLD20: p_spin_ < .002), which were positively associated (Fig. 3e). From 400 to 500 ms, word frequency was negatively associated with the gradient in contrast to OLD20 (p_spin_ = .006), semantic similarity (p_spin_ = .012) and position (p_spin_ = .02), which were positively associated (Fig. 3f). In addition, word length was negatively associated with the gradient in contrast to position (p_spin_ = .034; Fig. 3f). Other pairwise comparisons between the parameter’s association with the gradient were non-significant.

In summary, most time windows showed a similar pattern to the fMRI data (Fig. 3a), with semantic similarity showing a more positive association with the gradient than at least one of the word-level parameters (i.e., word length, OLD20, and word frequency). Furthermore, a pattern found consistently during the three latest time windows (from 200 to 500 ms) was that position showed a more positive association with the gradient in comparison to at least two of the word parameters.

In addition to investigating the differences between parameters in each time window, we also investigated differences between time windows for each parameter (Supplementary Tables S10 and S11). However, only word frequency showed a significant change in its association with the gradient across time (p_spin_ = .022): While word frequency showed a negative association with the gradient in four time windows (Fig. 3b-d,f), it showed a positive association with the gradient in the 300 to 400 ms time window (Fig. 3e; p_spin_ vs. 0 to 100 ms: .052; 100 to 200 ms: .01; 200 to 300 ms: .022; 400 to 500 ms: .004). Of note, this was also the only time window in which word frequency effects were negative (stronger brain activation for less frequent words), while the remaining time windows showed the opposite word frequency effect.

### 3.4 Summary of fMRI and MEG results

We observed that the principal connectivity gradient was significantly associated with the effects of several word and contextual parameters on brain activation (see overview in Figure 5). Parameters reflecting individual word characteristics, i.e., word length, OLD20 and word frequency, showed stronger effects on brain activation towards the sensory end of the principal gradient, both in fMRI data as well as in particular time windows in MEG data. While the effect directions of word parameters in the MEG data changed across the five time windows (cf. Fig. 2), the effect directions in the time windows showing significant associations with the gradient were *opposite* to the effect directions of these parameters found in the fMRI analysis. Thus, while fMRI and MEG data did not always converge on the direction of an effect for these word parameters, both neuroimaging modalities agreed on the association of these effects with the sensory end of the principal gradient.

**Figure 5.**
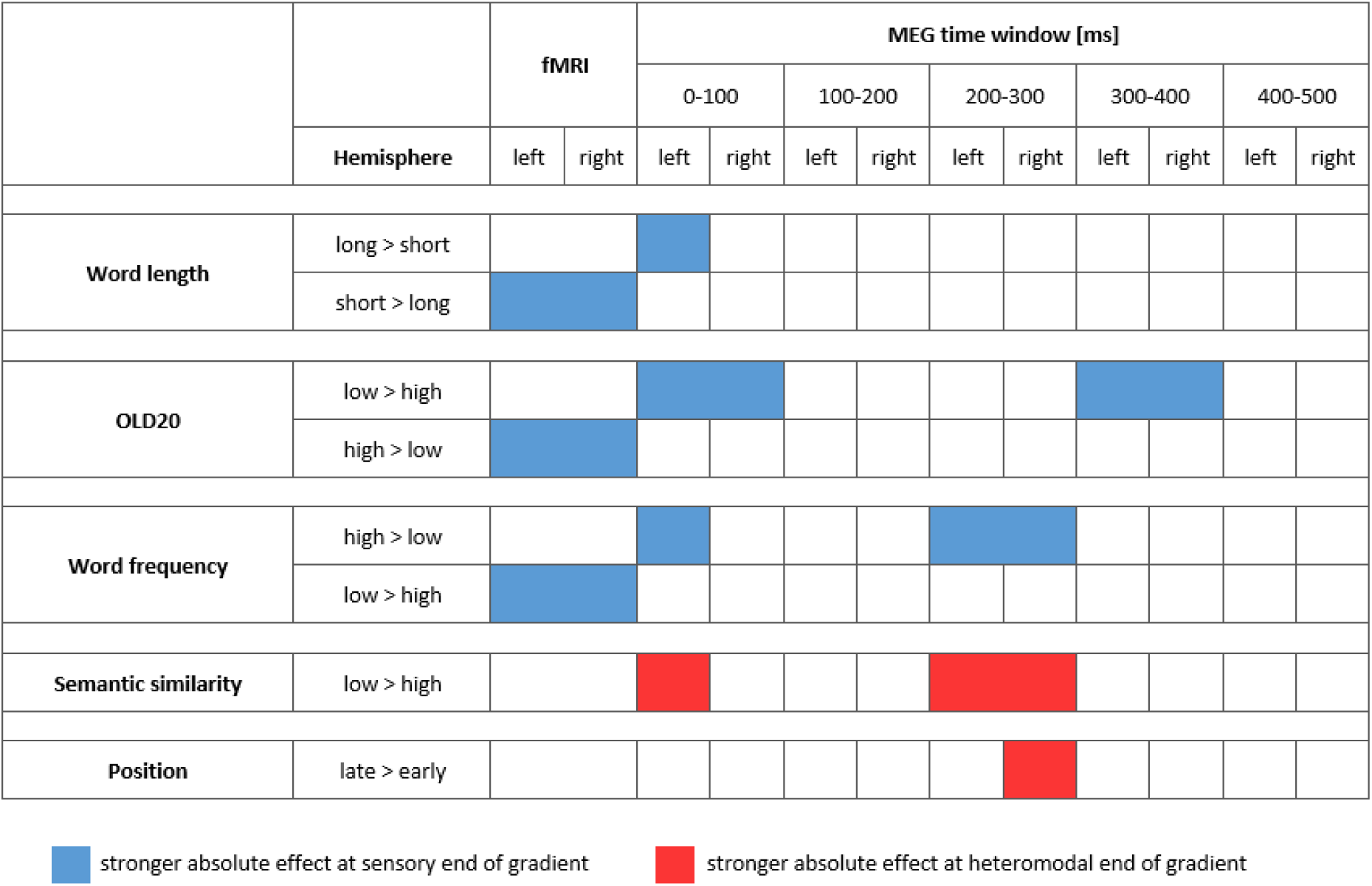
Overview of significant associations between the investigated word and contextual parameters with the principal gradient in fMRI data as well as in the five time windows investigated using MEG. The first column indicates the investigated parameter in bold and the second column the predominant direction of its effect on brain activation. For example, Word length long > short indicates that brain activation was higher for longer than shorter words, i.e., linear models indicated an increase in brain activation with increasing word length. Blue indicates the time windows (in case of MEG data) and hemispheres for which the absolute effect of the parameter was stronger at the sensory end of the gradient (i.e., the absolute effect decreases with increasing gradient value). Red indicates absolute effects of the parameter being stronger at the heteromodal end of the gradient (i.e., the absolute effect increases with increasing gradient value). In case of significant main effects of the gradient, both left and right hemisphere are coloured, whereas in case of significant gradient by hemisphere interactions, the hemisphere showing the stronger gradient effect is coloured.

In contrast to individual word parameters, contextual parameters, i.e., semantic similarity and position, showed stronger effects on brain activation towards the heteromodal end of the gradient. These associations with the gradient reached significance in the MEG data only (Fig. 5).

The observation that word parameters are associated with the sensory end while contextual parameters are associated with the heteromodal end of the gradient was further supported by the observed interactions between parameters and the gradient (cf. Fig. 2). In fMRI, we found significant differences between the three word parameters, associated with the sensory end of the gradient, in contrast to semantic similarity, which tended to be stronger towards the heteromodal end of the gradient. In MEG, the most consistent differences also occurred between the three word parameters, which were associated with the sensory end of the gradient, and the two contextual parameters, which were associated with responses towards the heteromodal end of the gradient. Finally, changes across time windows in a parameter’s association with the gradient were only found for word frequency (cf. Fig. 2). Most associations between linguistic variables and the principal gradient tended to be relatively stable through time.

## 4 Discussion

Language comprehension involves representing words in a visual to semantic pathway that contains increasingly abstract levels of processing and then integrating these representations with the linguistic context. We investigated the extent to which a cortical gradient that dissociates input-driven processes from abstract, memory-based processes (Margulies et al., 2016) can capture the neural organization of language comprehension. The brain response to different word and contextual parameters was robustly associated with this principal gradient. Linear models indicated that parameters reflecting linguistic representations of individual words affected neural responses at the sensory end of the principal gradient to a stronger extent, and this effect was consistent across two neuroimaging modalities, fMRI and MEG. These parameters included word length reflecting the visual complexity of the sensory input; Orthographic Levenshtein Distance (OLD) 20 reflecting familiarity with orthographic subunits; and word frequency reflecting familiarity with the lexical word form. MEG data revealed that in contrast to these individual word parameters, contextual parameters that reflect integrative processes across a sentence affect the heteromodal end of the gradient to a stronger extent. This dissociation between individual word representations and linguistic context was further supported when characterizing the gradient associations of each parameter: Reliable differences in gradient location were observed between contextual parameters (semantic similarity and word position) in comparison to individual word representations (word length, OLD20, and word frequency) in both MEG and fMRI data.

The observed dissociation between word representations and context is in line with the proposed distinction along the gradient between largely input-driven processes vs. abstract, integrative processes (Margulies et al., 2016): Word length and OLD20, in particular, describe representational characteristics of a word (the amount of visual information to be processed and the familiarity with orthographic subunits) and were previously shown to modulate the earliest processing stages of word recognition (∼100 ms; Assadollahi and Pulvermueller, 2003; Hauk and Pulvermueller, 2004; Hauk et al., 2009). Semantic similarity is associated with the ease of integrating a word with the preceding context, or the ease of predicting a word based on the context (e.g., Hofmann et al., 2022; Mitchell and Lapata, 2010; Mitchell et al., 2010; Pynte et al., 2008a; 2008b; Salicchi et al. 2023; Wang et al., 2010), while position reflects the amount of contextual information available in working memory (e.g., Dambacher et al., 2006; Kuperman et al., 2010). The distinction between these variables along the gradient indicates that brain regions proximal to sensory brain areas are predominantly involved in the processing of word representations, while heteromodal areas involved in mnemonic processes and integration are crucial for context-dependent linguistic processing. This finding places links between linguistic processes and brain regions in the broader macroscale organization of the cortex which supports higher-level functions in brain regions maximally distant from sensory cortices (Margulies et al., 2016; Wang et al., 2023).

Having established that the brain response to various linguistic processes linearly aligns with the hierarchical structure provided by the principal gradient, we assessed if this alignment is continuous or discrete by comparing the linear model to a three-step function. This comparison indicated that for many of the effects, the brain response changed continuously along the gradient, while some effects reflected discrete changes in brain activation. In the fMRI analysis, only OLD20 showed a continuous linear association with the gradient. In contrast, discrete changes were observed for word length and word frequency, suggesting that these parameters differentially recruit brain areas at the middle and the unimodal end of the principal gradient in comparison to brain areas at the heteromodal end. This is also indicated by their quadratic (u-shaped) association with the principal gradient as well as their linear association with Gradient 3, which differentiates the default mode network from the multiple demand network located in between the two ends of the principal gradient (discussed further below). The time-resolved MEG analysis, however, consistently indicated continuous rather than stepwise changes in brain activation along the principal gradient for all parameters, with only one effect (word frequency from 200 to 300 ms) showing comparable plausibility for stepwise changes. This suggests that at certain points in time, the brain response during linguistic processing differentiates along the whole brain in a predominantly continuous fashion aligned with the principal gradient, rather than differentially recruiting discrete processing regions that fall in a systematic sequence. These findings might support the fine-tuning of future computational models of language to adopt graded activation patterns between processing units (cf. Plaut, 2002; Ueno et al., 2011) and to explicitly quantify the location of each unit along the cortical hierarchy based on its location along the principal gradient.

Our study also provides important insights into the role of the significance of the principal gradient for human cognition. Previous evidence implicated the principal gradient for semantic cognition by demonstrating its link to individual differences in semantic task performance (Gonzalez Alam et al. 2021, Shao et al. 2022) as well as its association to fMRI responses during decisions on the semantic similarity of visual word pairs that varied in feature overlap (Wang et al., 2020) or association strength (Gao et al., 2022). A further fMRI study indicated the principal gradient reflected the brain response to linguistic properties of sentences during speech comprehension, including a ‘vocabulary’ component receiving high loadings from sentence-average word frequency and semantic diversity (U-shaped relation with the gradient), as well as a ‘sensory-motor’ component receiving high loadings from sentence-average concreteness and perception strength (linear relation with the gradient; Wu et al., 2022). Our findings extend this evidence by demonstrating that brain responses along the gradient capture a wide range of linguistic features, and that the gradient explicitly dissociates individual word representations from contextual information during sentence reading. More generally, the finding that both contextual parameters were associated with the heteromodal end of the gradient in MEG is in line with previous fMRI studies beyond the language domain that implicated this organizational pattern in memory-guided cognition (Smallwood et al., 2021) across a broad range of functions (e.g., motor control, Gale et al., 2022; automated rule-based behaviour, Vatansever et al., 2017; working memory, Murphy et al., 2018; 2019). The present MEG findings complement the previous fMRI evidence by showing that this organizational motif can impact on rapid features of information processing.

The additional use of MEG in the present study allowed us to investigate the temporal dynamics of the principal gradient’s association to linguistic and contextual effects in language processing, contributing to the debate about whether linguistic processing unfolds in a serial, predominantly feedforward manner, e.g., from visual to orthographic to semantic processing (cf. Carreiras et al., 2014; Dikker et al., 2020), or if processing across these levels occurs more interactively and in parallel (cf. Grainger and Ziegler, 2011; Price and Devlin, 2011). OLD20, word frequency, and semantic similarity were significantly associated with the gradient in both the earliest time window (i.e., 0 to 100 ms) and in a later time window (between 200 and 400 ms). Previous EEG studies of single-word reading that used multiple linear regression simultaneously controlling for visual/orthographic parameters reported the earliest onsets for lexical-semantic parameters in the time window between 100 and 200 ms (Dufau et al., 2015; Hauk et al., 2006) or even later after 250 ms (Laszlo and Federmeier, 2014). In constrast, a previous MEG study assessing word recognition in a repetition priming context found effects in the pre-100 ms time window (Eisenhauer et al., 2022) similar to the present study. Another study examining semantic associations for pairs of words presented serially uncovered effects of the type of semantic relationship within the first 100 ms of the onset of the second word (Teige et al., 2019). Potentially, in context-dependent word reading, an anticipation of upcoming words (e.g., Federmeier, 2007; Kuperberg and Jaeger, 2016) might result in an earlier onset of lexical-semantic processing.

The direction of each investigated parameter’s effect on the evoked response across the surface parcels changed over time; yet, for most parameters, the linear relation to the gradient (i.e., whether a parameter’s absolute effect was strongest at the sensory or heteromodal end of the gradient) did not change. The fact that most parameters were associated with the gradient in both early and later time windows, and the absence of explicit evidence for temporal changes for most parameters, is compatible with an interactive account of parallel processing across multiple linguistic levels (e.g., Clarke et al., 2011; 2013; Cornelissen et al., 2009; Eisenhauer et al., 2022; Kaestner et al., 2021; Mollo et al., 2017; 2018; Teige et al., 2019). This interactive processing is also reflected along the principal gradient: both ends of the gradient are activated in similar time windows but represent different aspects of linguistic information.

Only word frequency showed a changing relation with the principal gradient over time: while word frequency affected brain activation to a stronger extent at the sensory end of the gradient in most time windows (reaching significance from 0 to 100 and 200 to 300 ms), word frequency effects in the 300 to 400 ms window tended more towards the heteromodal end of the gradient. This change along the gradient co-occurred with a change in the direction of word frequency effects: The evoked brain response was higher for low in contrast to high frequency words from 300 to 400 ms, while the reverse pattern was found in the remaining time windows. A possible explanation for the change in effect direction as well as in the relation to the gradient across time might be that word frequency reflects word familiarity across multiple linguistic levels, and therefore differentially modulates temporally segregated stages during language processing which involve a different set of sensory vs. heteromodal brain regions.

Besides the relative temporal stability of the association with the principal gradient for most parameters, supplementary investigations of the present MEG data indicated that effects of individual word characteristics also align with connectivity Gradients 2 (capturing the distinction between visual and motor cortex) and 3 (distinguishing the multiple demand and default mode network) – exclusively in time windows in which no significant associations with the principal gradient were found. Additionally, quadratic associations with the principal gradient, which correlate with Gradient 3 (r = .68), also almost exclusively occurred in time windows not showing linear associations, with the exception of the early word length effect (0 to 100 ms). This indicates that processing of individual word characteristics shows highly dynamic fluctuations across time between different brain states reflected in dimensions of cortical connectivity at rest. The fMRI analysis, which is not able to tease apart these fluctuations across time, indicated that effects of individual word characteristics were aligned both with the principal gradient and Gradient 3, and partly showed quadratic associations with the principal gradient, potentially reflecting a sum of the temporal dynamics between the gradients observed in the MEG analysis. While the principal gradient predominantly reflects the continuum from input-driven to contextual processing, Gradient 2 might reflect a continuum from the input-driven visual code to articulatory codes represented in motor cortex (e.g., Kaestner et al., 2021; Wheat et al., 2010), and Gradient 3 might reflect a continuum from automatic to more demanding processes that require control from executive brain regions (cf. Duncan, 2010; 2013). Organizing the brain response to individual word characteristics during reading in a continuous fashion along each of these three dimensions at different points in time might ensure maximal engagement of currently appropriate cortical networks while maximally down-regulating the networks involved in opposing functions. These assumptions should be tested in future studies explicitly manipulating the mapping between visual and articulatory codes as well as reading demands.

When comparing linear associations with the principal gradient between fMRI and MEG, there was a high convergence regarding the principal gradient location most strongly affected by each parameter – namely, the sensory end for word length, OLD20, and word frequency, and the heteromodal end for semantic similarity. However, fMRI showed opposite effect directions for word length, OLD20, and word frequency compared with effects during the time windows that showed significant associations with the principal gradient in MEG data. While the majority of previous studies found stronger brain responses for less frequent words (e.g., Embick et al., 2001; Fiebach et al., 2002; Hauk and Pulvermueller, 2004; Hauk et al., 2008; Protopapas et al., 2016; Schuster et al., 2016), several studies also reported the reverse pattern. For example, using fMRI, Kronbichler et al. (2004) and Zhuang et al. (2011) found higher activation for low frequency words in the left middle temporal and inferior frontal gyri, while Yarkoni et al. (2008b) found the opposite pattern. Likewise, electrophysiological and MEG studies show opposing directions for word frequency effects in time windows predominantly associated with semantic processing (∼ 200 to 400 ms; e.g., Embick et al., 2001; Faisca et al., 2019; Hauk and Pulvermueller, 2004; Simon et al., 2012). Word length and OLD20 effects can also show different effect directions depending on the time window (Assadollahi and Pulvermueller, 2004; Eisenhauer et al., 2022; Hauk and Pulvermueller, 2004). It is difficult to interpret neural effect direction for any parameter, as both stronger and weaker brain responses may relate to more efficient processing. This is further complicated by the potentially opposing effects of the parameters (see Andrews, 1997; Chetail, 2015, for reviews): On the one hand, processing may be facilitated for words that are shorter, as well as orthographically or lexically more familiar. On the other hand, shorter and more familiar words are less well distinguishable from other words, increasing competition which may hinder their recognition (cf. Chen and Mirman, 2012; Grainger and Jacobs, 1996; Hutchison et al., 2008; McClelland and Rumelhart, 1981). These opposing effects might contribute to the different effect directions in brain activation observed for these variables in the present investigation as well as in previous studies. In a separate behavioural experiment included in the present study, we observed that faster self-paced reading responses were made for words that were shorter, orthographically more similar to other words, and that occurred earlier in the sentence, suggesting more efficient processing, in line with previous findings (e.g., Dufau et al., 2015; Kuperman et al., 2010; Yarkoni et al., 2008b). Word frequency and semantic similarity did not significantly affect self-paced reading response times in the present study, potentially as self-paced reading responses were fast (mean: 380 ms) and more sensitive to measures reflecting earlier stages of word recognition. Note that a previous investigation of the same dataset with a different set of parameters that did not include OLD20 did find a significant word frequency effect (Kapteijns and Hintz, 2021). As OLD20 and word frequency are correlated (r = −.65), word frequency may not have contributed to reading times on top of OLD20 effects. Nevertheless, previous studies quite consistently provided evidence for faster reading of high vs. low frequency words (e.g., Dufau et al., 2015; Eisenhauer et al., 2022; Juhasz and Rayner, 2007) as well as high vs. low semantic similarity (e.g., Hofmann et al., 2022; Pynte et al., 2008a; 2008b; Salicchi and Lenci, 2021; Wang et al., 2010).

To some extent, the neural activation patterns observed in the present study are correlates of the observed behavioural effects. However, behavioural observations do not reflect processing dynamics across time, and include a specific decision component that is not present during natural reading. Thus, while behavioural findings add crucial information about how different word and contextual features contribute to efficient reading, they might not reflect all the processes involved at the neural level. Similarly, fMRI responses predominantly reflect a sum of temporally stable effects, while MEG may also capture processes that are more short-lived (Singh, 2012). Furthermore, the two modalities differ in their sensitivity to aspects of the neural signal; for example, fMRI is sensitive to both synchronized and desynchronized neural activation, while the evoked MEG response mainly reflects synchronized neural activation (Singh, 2012), and the correspondence between the two modalities varies across brain regions (Shafiei et al., 2022). It might therefore not be surprising that effect directions in fMRI do not correspond with effect directions in MEG in most time windows, and opposite effect directions for the processing of letter vs. symbol strings in the left occipital-temporal cortex between the two neuroimaging modalities were found previously (Vartiainen et al., 2011). Given the complex nature of neural dynamics during language comprehension, combining fMRI and MEG is a very powerful approach to reveal a more detailed picture and prevent overinterpretation of results. Based on the present findings, brain location along the principal gradient, rather than effect direction, might be a more robust neural correlate of linguistic variables.

A limitation of the present study is its focus on the investigation of evoked magnetic fields, while oscillatory responses in the MEG signal are also involved in language processing (for reviews, see Hauk et al., 2017; Meyer 2018; Prystauka and Lewis, 2019) and known to correlate with fMRI responses (for reviews, see Goense et al., 2012; Singh, 2012). The relation of neural oscillations to the principal gradient in language processing should be investigated in future studies. Future studies are also needed to confirm if the present gradient associations replicate across different input modalities (i.e., reading vs. listening; words vs. pictures) that are hypothesized to be organised in a similar way.

To conclude, the present study found that the brain’s response to different linguistic characteristics varies systematically along the principal cortical gradient while expressing opposing activation patterns: Individual word representations maximally recruit the sensory end of the gradient while showing gradually lower responses in brain regions closer to the heteromodal end. In contrast, contextual linguistic processes show the reverse relationship with the principal gradient with increasing activation towards the heteromodal end. This dissociation along the gradient persisted during multiple time windows, indicating that input-driven versus contextual linguistic processes operate in parallel, albeit focused in distinct cortical locations. The gradient approach complements investigations of individual brain regions’ contributions to language comprehension by linking linguistic processing to an organizational motif that re-emerges during different aspects of cognition. The graded brain response along this motif might contribute to efficient language comprehension by maximizing linguistic representations in functionally relevant regions while down-regulating representations in regions implicated in opposing or irrelevant processes. Furthermore, the gradient might have the potential to reveal convergent patterns across neuroimaging modalities (similar location along the gradient) in the presence of divergent aspects (opposite effect directions), and thus contributes to identifying robust neural correlates of cognitive function.

## Data and code availability statement

The neuroimaging data used in this study is publicly available at https://data.donders.ru.nl/collections/di/dccn/DSC_3011020.09_236?0. The preprocessed MEG data and part of the analysis code is publicly available at https://data.donders.ru.nl/collections/di/dccn/DSC_3011020.09_410?7. The behavioural data is publicly available at https://hdl.handle.net/1839/fa97ca32-897d-4f3b-b3f9-80cb84a5a180. New analysis code generated for this study is publicly available at https://osf.io/5fxbd/.

## Acknowledgements

Data were provided (in part) by the Donders Institute for Brain, Cognition and Behaviour, Radboud University Nijmegen. We thank Jan-Mathijs Schoffelen and Sophie Arana for sharing neuroimaging data and analysis code, Florian Hintz for sharing behavioural data and analysis code, and Jean-Rémi King for sharing analysis code. This work was supported by the European Research Council (Project ID: 771863 – FLEXSEM to EJ).

## CRediT author contributions

**Susanne Eisenhauer:** Conceptualization, Methodology, Formal analysis, Data Curation, Writing - Original Draft, Writing - Review & Editing, Visualization

**Tirso Rene del Jesus Gonzalez Alam:** Methodology, Formal analysis

**Piers L. Cornelissen:** Writing - Review & Editing, Supervision

**Jonathan Smallwood:** Conceptualization, Writing - Review & Editing

**Elizabeth Jefferies:** Conceptualization, Writing - Review & Editing, Supervision, Funding acquisition

