## Supplementary Material for "Individual word representations dissociate from linguistic context along a cortical unimodal to heteromodal gradient"

**Supplementary Materials. Correlations between stimulus parameters**

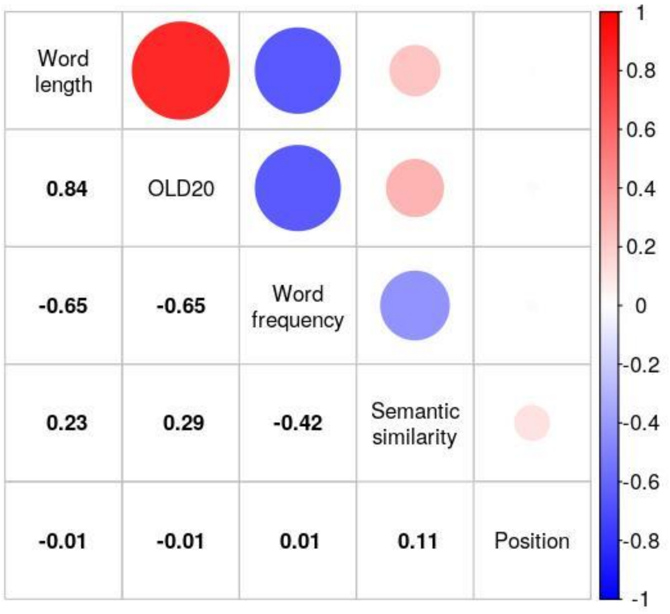


**Figure S1.** Correlations between stimulus parameters of content words included in fMRI and MEG analyses.

**Supplementary results 1. Significant effects of word and sentence level parameters on brain activation in fMRI data.**

Significant clusters of word and sentence level parameters on BOLD activation were obtained from the group level contrasts using Gaussian random-field theory at a cluster-forming threshold of $z = 3.1$. Clusters were familywise-error-corrected at a significance level of $p = .05$. Word length and word frequency negatively modulated brain activation in left posterior temporal regions (Fig. S2a,b), indicating that brain activation was higher for shorter than longer words as well as for less vs. more frequent words. Semantic similarity negatively modulated brain activation in the left anterior temporal lobe (Fig. S2c), indicating reduced brain activation for words that were more similar to the preceding five-word phrase. Position positively modulated brain activation in more distributed brain regions, including bilateral occipital and posterior temporal cortices, and left-lateralized anterior temporal cortex and precentral gyrus (Fig. S2d). No significant effects of OLD20 were found. Next, we identified the average gradient value of the parcels within the significant cluster, expressed as a percentile (converting gradient scores to percentiles from 0% at the unimodal end to 100% at the heteromodal end). This average value was at the $40^{th}$ percentile for word length, at the $48^{th}$ percentile for word position, at the $59^{th}$ percentile for word frequency, and at the $61^{st}$ percentile for semantic similarity. These percentiles indicate that brain regions at relatively intermediate locations along the principal gradient are involved in processing the associated linguistic information. Overall, the significant clusters are in well-established parts of the language network, with posterior temporal cortex implicated in lexical access (Dikker et al., 2020; Hickok and Poeppel, 2007) and semantic control (Jackson, 2021; Lambon Ralph et al., 2017), anterior temporal cortex in long-term semantic representation (Lambon Ralph et al., 2017) and composition (Pylkkänen, 2019), and precentral gyrus in articulatory code activation (Kaestner et al., 2021; Wheat et al., 2010).

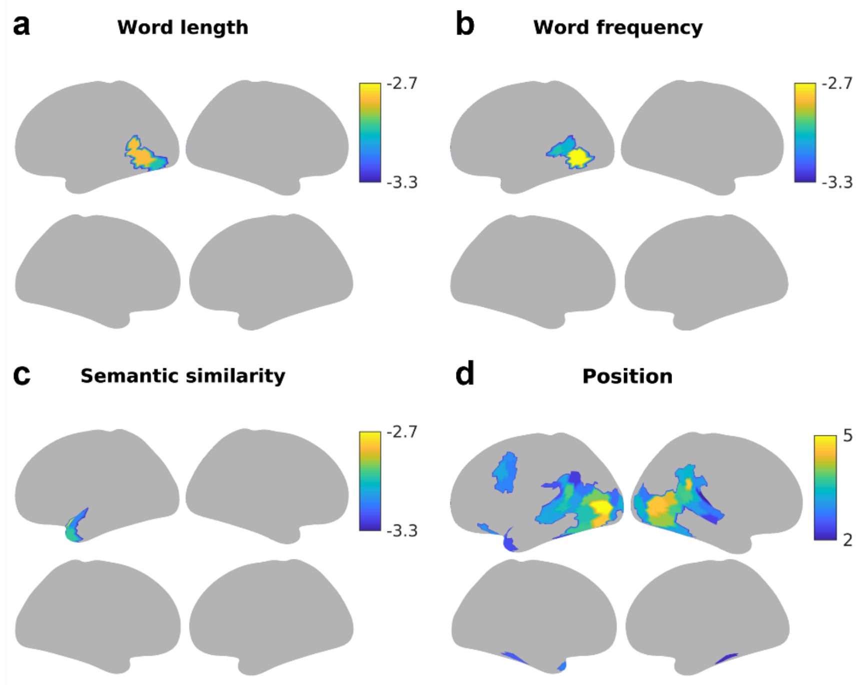


**Figure S2.** Significant effects of a) word length, b) word frequency, c) semantic similarity and d) position on fMRI BOLD activation (top: lateral view, bottom: medial view).

**Supplementary results 2. Associations of word and sentence level parameters with Gradients 2 and 3 in fMRI data**

Gradient 2, which differentiates visual from motor cortex (Margulies et al., 2016; Fig. S3a), did not show a significant relation with any of the investigated parameters (all $p > .05$; Fig. S4; Tables S12 and S13). Gradient 3 differentiates the default mode network (DMN) from the multiple demand network (MDN; Margulies et al., 2016; Fig. S3b). Effects of the individual word parameters, i.e., word length ($p_{spin} < .002$), OLD20 (stronger brain activation for higher OLD20/less orthographic familiarity; $p_{spin} = .038$), and word frequency (stronger brain activation for less frequent words; $p_{spin} < .002$), were significantly stronger at the MDN end of Gradient 3 (Fig. S5; Tables S14 and S15). Semantic similarity effects, in contrast, were significantly stronger at the DMN end of Gradient 3 in the left hemisphere (gradient by hemisphere interaction: $p_{spin} < .002$; Fig. S5). Position effects were not significantly associated with Gradient 3 ($p > .05$).

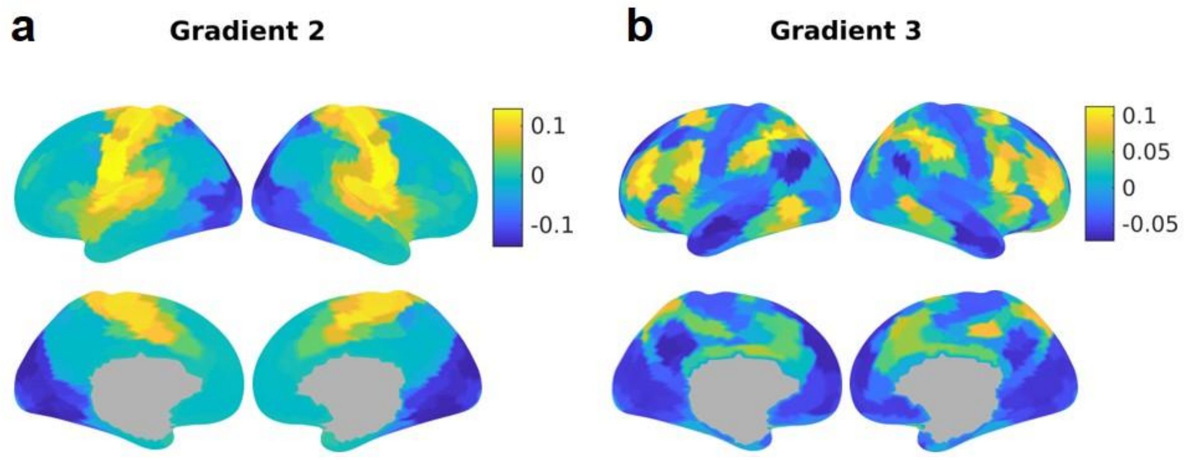


**Figure S3.** Connectivity Gradients 2 (a) and 3 (b).

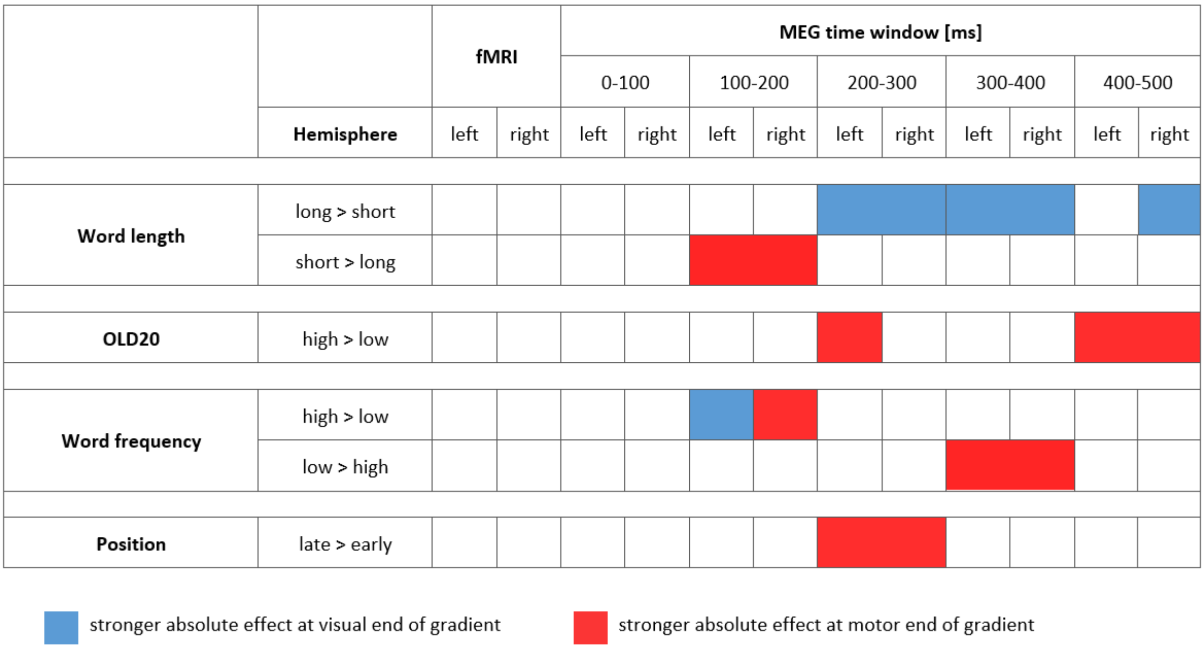


**Figure S4.** Overview of significant associations between the investigated word and contextual parameters with Gradient 2 in fMRI data as well as in the five time windows investigated using MEG. The first column indicates the investigated parameter in bold and the second column the predominant direction of its effect on brain activation. For example, Word length long > short indicates that brain activation was higher for longer than shorter words, i.e., linear models indicated an increase in brain activation with increasing word length. Blue indicates the time windows (in case of MEG data) and hemispheres for which the absolute effect of the parameter was stronger at the visual end of Gradient 2 (i.e., the absolute effect decreases with increasing gradient value). Red indicates absolute effects of the parameter being stronger at the motor end of Gradient 2 (i.e., the absolute effect increases with increasing gradient value). In case of significant main effects of Gradient 2, both left and right hemisphere are coloured, whereas in case of significant Gradient 2 by hemisphere interactions, the hemisphere showing the stronger Gradient 2 effect is coloured.

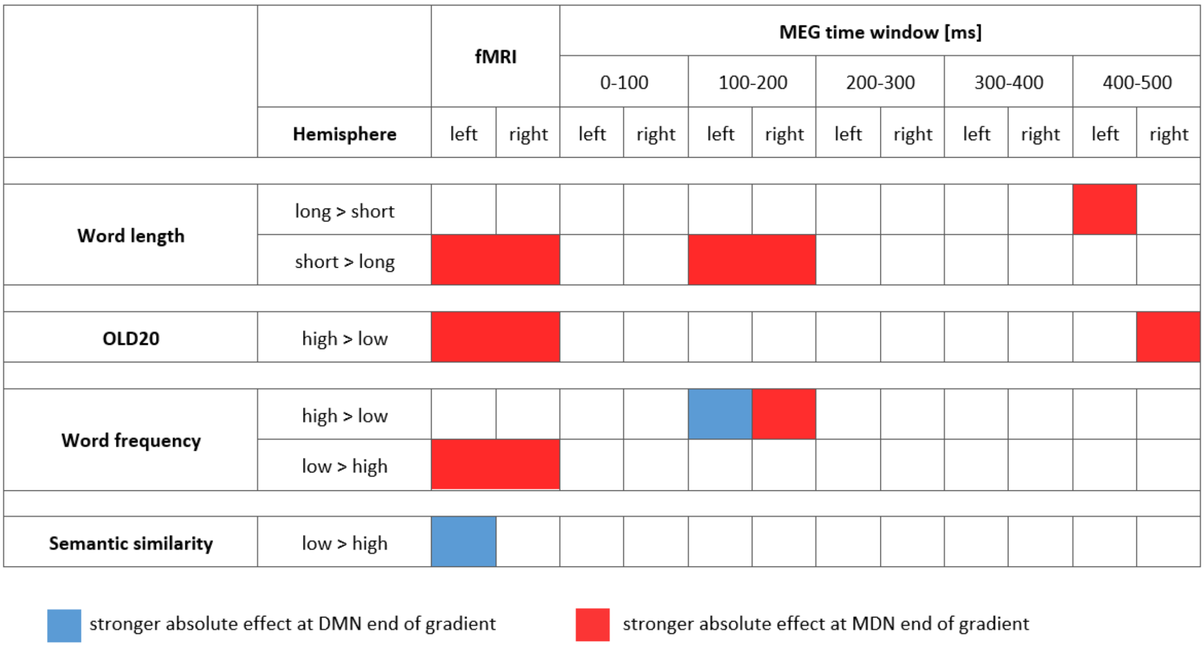


**Figure S5.** Overview of significant associations between the investigated word and contextual parameters with Gradient 3 in fMRI data as well as in the five time windows investigated using MEG. The first column indicates the investigated parameter in bold and the second column the predominant direction of its effect on brain activation. For example, Word length long > short indicates that brain activation was higher for longer than shorter words, i.e., linear models indicated an increase in brain activation with increasing word length. Blue indicates the time windows (in case of MEG data) and hemispheres for which the absolute effect of the parameter was stronger at the DMN end of Gradient 3 (i.e., the absolute effect decreases with increasing gradient value). Red indicates absolute effects of the parameter being stronger at the MDN end of Gradient 3 (i.e., the absolute effect increases with increasing gradient value). In case of significant main effects of Gradient 3, both left and right hemisphere are coloured, whereas in case of significant Gradient 3 by hemisphere interactions, the hemisphere showing the stronger Gradient 3 effect is coloured.

**Supplementary results 3. Quadratic associations of word and sentence level parameters with the principal gradient in fMRI data**

We focused our main investigations on linear associations with the principal gradient as the gradient itself provides a linear organization of brain function and simple linear models are more readily interpretable and less prone to overfitting in the face of the complex neuroimaging data than more complex functions (cf. King et al., 2018). However, in this supplementary analysis, we also assessed if a parameter's relationship to the gradient was better explained by a quadratic than a linear function by comparing the original linear model to a model that additionally contained the quadratic term of the parameter in an interaction with hemisphere. This can reveal if linguistic parameters recruit brain regions at an intermediate location along the gradient more strongly than brain regions at both ends or vice versa. Nested model comparison was based on an F-test using the anova function in R. A significant difference indicates that the more complex quadratic model should be favoured; otherwise, the linear model should be favoured. If the quadratic model was favoured, the spin permutation procedure was used to assess significance of the quadratic term. Results were considered significant if this was confirmed in the control analysis modelling presentation duration instead of word length. Significant quadratic (u-shaped) associations with the gradient were found for word length and word frequency (all $p_{spin} < .002$; Fig. S6), indicating that negative word length effects (higher brain activation for shorter words) and negative word frequency effects (higher brain activation for less frequent words) increased towards the middle of the principal gradient. No significant quadratic associations with the gradient were found for OLD20, semantic similarity and position.

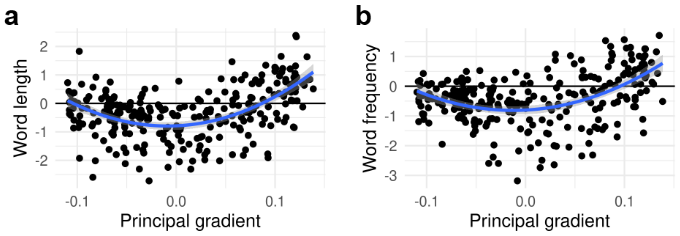


**Figure S6.** Quadratic associations with the principal connectivity gradient for a) word length and b) word frequency effects on fMRI BOLD activation across 269 cortical parcels. Z values from the group-level general linear model are plotted against the principal gradient values from Margulies et al. (2016). The lowest gradient values are associated with sensory cortices, while the highest gradient values are associated with heteromodal cortices.

**Supplementary results 4. Associations of word and sentence level parameters with the principal gradient in fMRI data when modelling the actual presentation duration of the stimuli**

When modelling the actual presentation duration of each word in the GLMs, we found similar associations with the principal connectivity gradient as in the analysis presented in the main manuscript which was based on modelling a fixed 1 s duration for each word in the GLMs (from now on referred to as 'main analysis'). That is, significant associations with the principal gradient were found for word length ($p_{spin} < .002$), OLD20 ($p_{spin} < .002$), and word frequency ($p_{spin} = .006$), but not for semantic similarity and position (Fig. S7). The relative shape of the association between these variables and the principal gradient was similar across analyses (although the intercept was shifted); while negative word length and word frequency effects increased towards the unimodal end of the gradient as in the main analysis, in this new analysis, positive word length and word frequency effects also increased towards the heteromodal end of the gradient. Given that modelling the actual presentation duration has the disadvantage of underestimating the temporal duration for which word processing influences the BOLD activation, we also performed an alternative control analysis that included the actual presentation duration as a covariate of no interest in the GLM, while assuming a fixed presentation duration of 1s and omitting word length. This also showed increasing negative word frequency effects towards the unimodal end of the principal gradient.

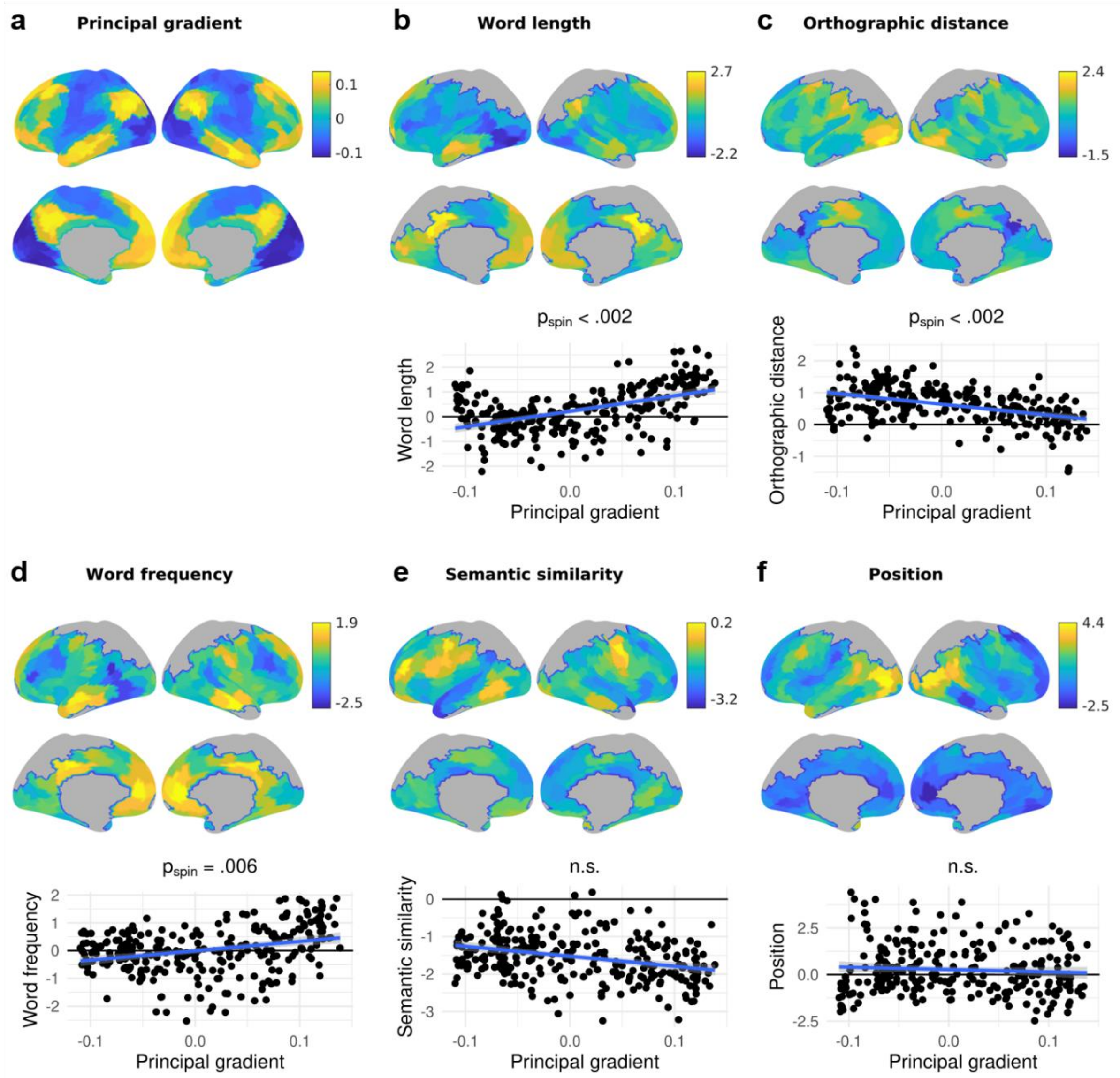


**Figure S7.** The principal connectivity gradient (a) as well as parameter effects on brain activation (b-f; top: lateral view, middle: medial view) and their dependence on the principal gradient (bottom) across the whole brain when modelling the actual word presentation duration instead of a fixed duration of 1 s. In b-f, the top and middle panels show the effect of the respective word parameter on fMRI BOLD activation across 269 cortical parcels, indicated by the z values from the group-level general linear model. Positive z values indicate an increase in brain activation with increasing parameter values, while negative z values indicate a decrease in brain activation with increasing parameter values. Grey parcels were not analysed due to limited field of view during data acquisition. The bottom panels show the dependence of these effects on the principal gradient across parcels, i.e., the z values are plotted against the principal gradient values from Margulies et al. (2016). The lowest gradient values are associated with sensory cortices, while the highest gradient values are associated with heteromodal cortices. $p_{spin}$ indicates p values from spin permutation; n.s. = not significant.

**Supplementary results 5. Stepwise associations of word level parameters with the principal gradient in fMRI data**

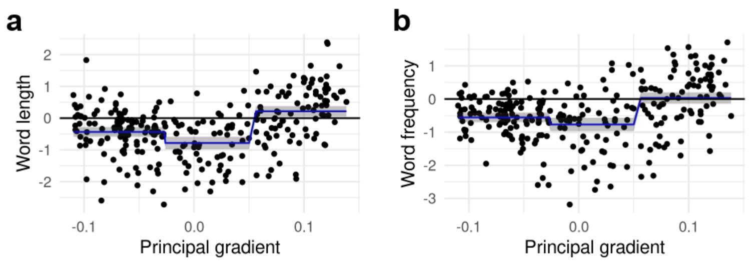


**Figure S8.** Stepwise associations with the principal connectivity gradient for a) word length and b) word frequency effects on fMRI BOLD activation across 269 cortical parcels. Z values from the group-level general linear model are plotted against the principal gradient values from Margulies et al. (2016). The lowest gradient values are associated with sensory cortices, while the highest gradient values are associated with heteromodal cortices. Results are described in the results section of the main manuscript.

**Supplementary results 6. Gradient 2 and 3 location of word and contextual parameters' effects in fMRI data**

For Gradient 2, an ANOVA computing the interaction between gradient value, parameter, and hemisphere with the parameter's effect on brain activation as dependent variable did not reveal any significant effect (Fig. S9a; Tables S16 and S17). For Gradient 3, the gradient value by parameter interaction was significant, also verified using spin permutation ($p_{spin}$ < .001), but not further modulated by hemisphere (Fig. S10a; Tables S18 and S19). Pairwise follow-up tests revealed that the association between Gradient 3 and brain activation for word length differed significantly from that of OLD20 ($p_{spin}$ < .002), semantic similarity ($p_{spin}$ < .002) and position ($p_{spin}$ = .002). In addition, word frequency differed significantly from OLD20 ($p_{spin}$ = .028) and semantic similarity ($p_{spin}$ < .002). Word length and word frequency were more strongly associated with the MDN end of Gradient 3 than the respective other parameters.

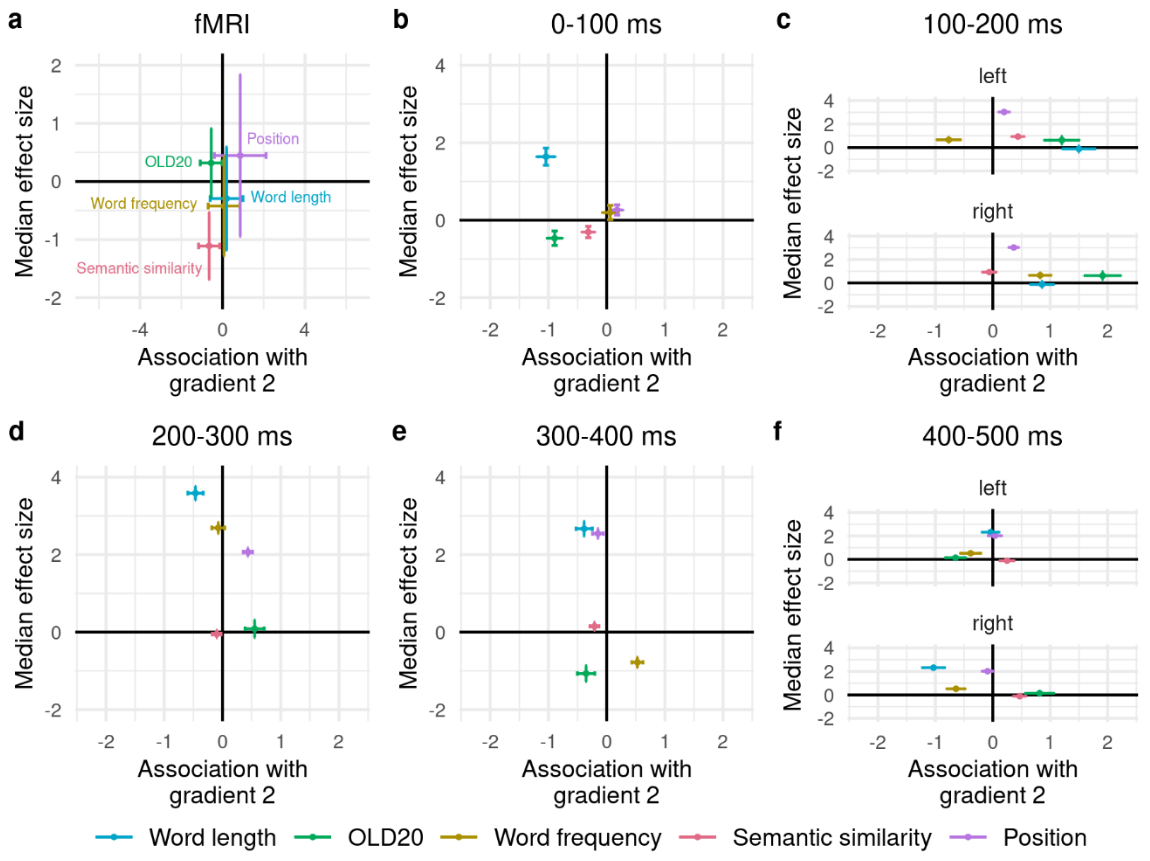


**Figure S9.** Association (i.e., linear model estimate) of each parameter with Gradient 2 as indicated on the x axis in a) fMRI BOLD activation and b-f) MEG-measured brain activation across five time windows. Negative values on the x axis indicate a stronger effect of the parameter towards the visual end of

Gradient 2, while positive values indicate a stronger effect towards the motor end of Gradient 2. Horizontal error bars indicate standard errors from linear models. The median effect size (i.e., general linear model z value in fMRI data; linear mixed model estimate in MEG data) of each parameter on brain activation across parcels is indicated on the y axis. Negative values indicate a negative effect direction (i.e., decreasing brain activation with increasing parameter values) for most parcels, while positive values indicate a positive effect direction for most parcels. Vertical error bars indicate the standard deviation of the parameter's effect size across parcels.

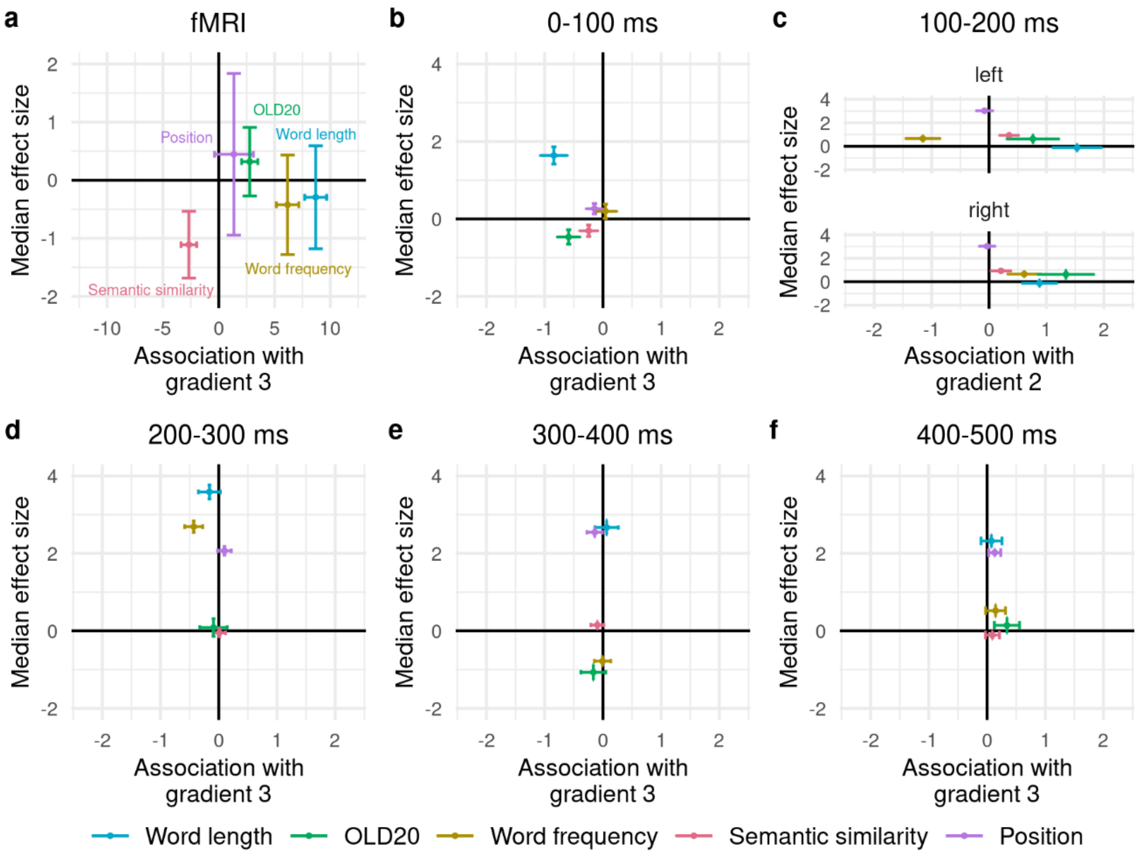


**Figure S10.** Association (i.e., linear model estimate) of each parameter with Gradient 3 as indicated on the x axis in a) fMRI BOLD activation and b-f) MEG-measured brain activation across five time windows. Negative values on the x axis indicate a stronger effect of the parameter towards the DMN end of Gradient 3, while positive values indicate a stronger effect towards the MDN end of Gradient 3. Horizontal error bars indicate standard errors from linear models. The median effect size (i.e., general linear model z value in fMRI data; linear mixed model estimate in MEG data) of each parameter on brain activation across parcels is indicated on the y axis. Negative values indicate a negative effect direction (i.e., decreasing brain activation with increasing parameter values) for most parcels, while positive values indicate a positive effect direction for most parcels. Vertical error bars indicate the standard deviation of the parameter's effect size across parcels.

**Supplementary results 7. Significant effects of word and sentence level parameters on brain activation in MEG data.**

Significant effects of word and sentence level parameters on brain activation in MEG data were based on the linear mixed model p values, Bonferroni-corrected for multiple comparisons across 350 parcels. No significant effects were found for word length, OLD20, word frequency, and semantic similarity. Potentially, cluster-based statistics might have more power to reveal significant effects as they restrict the number of significant comparisons to the cluster level (Maris and Oostenveld, 2007); however, we did not apply such statistics as the major aim of the present study was the investigation of the relation of whole-brain activation patterns with the principal gradient. Only position showed significant positive effects, i.e., greater brain activation for words later in the sentence, in the time windows between 100 and 500 ms after word onset (Fig. S11). These position effects were widely distributed across bilateral cortices from 100 to 200 ms with a peak in the right superior frontal gyrus (Fig. S11a), as well as from 300 to 400 ms with a peak in the left frontal pole (Fig. S11c). Position effects were restricted to the right precuneus from 200 to 300 ms (Fig. S11b), and to the left middle/superior frontal gyrus from 400 to 500 ms (Fig. S11d). The gradient values of these peak locations (for 100 to 200 and 300 to 400 ms) and cluster locations (for 200 to 300 and 400 to 500 ms) across these four time windows were localised as percentiles along the principal gradient (where 0% is at the unimodal end and 100% is at the heteromodal end). These effects, in temporal order (i.e., for Fig. S11 a, b, c, and d), fell at the $54^{th}$, $97^{th}$, $89^{th}$, and $68^{th}$ percentile along the gradient, respectively. Thus, for the latter three time windows, (peak) activations of significant clusters fell at a position towards the heteromodal end of the gradient, which is in line with the significant positive association of position effects with the gradient from 200 to 300 ms (cf. Fig. 4i).

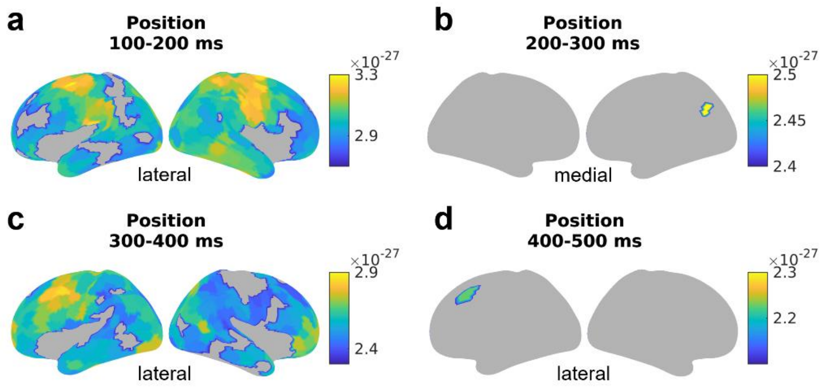


**Figure S11.** Significant effects of position on MEG source activation from a) 100 to 200, b) 200 to 300, c) 300 to 400, and d) 400 to 500 ms.

**Supplementary results 8. Associations of word and sentence level parameters with Gradients 2 and 3 in MEG data**

The effects of most investigated parameters, except for semantic similarity, on MEG-measured brain activation were significantly associated with Gradient 2 in particular time windows (Fig. S4; Tables S20 and S21). Brain activation was higher for shorter vs. longer words from 100 to 200 ms, and this word length effect was significantly stronger at the visual end of Gradient 2 ($p_{spin}$ = .042). In the time windows 200 to 300 ms, 300 to 400 ms (bilaterally) and 400 to 500 ms (right hemisphere), brain activation was higher for longer vs. shorter words, and this effect was higher at the motor end of Gradient 2 ($p_{spin}$ = .002; .032; gradient by hemisphere interaction: .022, respectively).

For OLD20, brain activation was predominantly higher for words that were less orthographically familiar, and this effect was stronger at the motor end of Gradient 2 from 200 to 300 ms in the left hemisphere (gradient by hemisphere interaction: $p_{spin} < .002$) and from 400 to 500 ms bilaterally (gradient by hemisphere interaction: $p_{spin}$ = .038; main effect: $p_{spin}$ = .054; interpreted as a main effect as only the main effect was significant when controlling for presentation duration instead of word length).

Regarding word frequency, brain activation was higher for more vs. less frequent words from 100 to 200 ms. This effect was significantly stronger at the visual end of Gradient 2 in the left hemisphere, while it was significantly stronger at the motor end of Gradient 2 in the right hemisphere (gradient by hemisphere interaction: $p_{spin}$ = .002). From 300 to 400 ms, brain activation was higher for less vs. more frequent words, and this effect was significantly stronger at the motor end of Gradient 2 ($p_{spin}$ = .01).

For position, brain activation was higher for later vs. earlier words in the sentence, and this effect was stronger at the motor end of Gradient 2 from 200 to 300 ms ($p_{spin}$ = .014).

For Gradient 3, only a few significant associations were found for the effects of word length, OLD20 and word frequency on brain activation (Fig. S5; Tables S22 and S23). The word length effect from 100 to 200 ms (short > long) was stronger at the MDN end of Gradient 3 ($p_{spin}$ = .006). The word length effect from 400 to 500 ms (long > short) was stronger at the MDN end of Gradient 3 in the left hemisphere (gradient by hemisphere interaction: $p_{spin}$ = .014). The OLD20 effect from 400 to 500 ms (high > low OLD20) was stronger at the MDN end of Gradient 3 in the right hemisphere (gradient by hemisphere interaction: $p_{spin}$ = .016). The word frequency effect from 100 to 200 ms (high > low) was significantly stronger at the

DMN end of Gradient 3 in the left hemisphere, and stronger at the MDN end of Gradient 3 in the right hemisphere (gradient by hemisphere interaction: $p_{spin} < .002$).

**Supplementary results 9. Quadratic associations of word and sentence level parameters with the principal gradient in MEG data**

As in the fMRI analysis, we also assessed if a parameter's relationship with the gradient was better explained by a quadratic than a linear function via model comparison. For effects that favoured a quadratic model, the spin permutation procedure was used to assess significance of the quadratic term. Results were considered significant if this was confirmed in the control analysis modelling presentation duration instead of word length. Significant quadratic associations with the gradient were found for the individual word characteristics, i.e., word length, OLD20, and word frequency, at different time windows (see Fig. S12) and are described in more detail below. No significant quadratic associations with the gradient were found for the contextual parameters, semantic similarity and position.

Word length effects from 0 to 100 ms showed a u-shaped relationship with the gradient, indicating that a positive word length effect (higher source activation for longer words) increased towards both ends of the principal gradient (main quadratic effect of the principal gradient: $p_{spin}$ = .006; interaction of the linear effect of the gradient with hemisphere: $p_{spin}$ = .01; Fig. S12a). A similar pattern was observed from 400 to 500 ms in the right hemisphere (interaction of the quadratic effect of the principal gradient with hemisphere: $p_{spin}$ = .022; Fig. S12c). However, in between these two time windows, from 100 to 200 ms, a u-shaped association with the gradient indicated that a negative word length effect (higher source activation for shorter words) increased towards the middle of the gradient (main quadratic effect of the principal gradient: $p_{spin}$ = .008; Fig. S12b).

OLD20 showed an inverted u-shaped association with the principal gradient from 200 to 300 ms in the left hemisphere, indicating that a positive OLD20 effect (higher source activation for less orthographically familiar words) increased towards an intermediate location along the principal gradient (interaction of the quadratic effect of the principal gradient with hemisphere: $p_{spin}$ = .05; Fig. S12d).

Word frequency effects from 100 to 200 ms showed a u-shaped association with the principal gradient in the left hemisphere, indicating that the positive word frequency effect (higher source activation for more frequent words) increased towards both ends of the principal gradient. However, in the right hemisphere, the positive word frequency effect increased towards the middle of the principal gradient, reflected by an inverted u-shaped association (interaction of the quadratic effect of the principal gradient with hemisphere: $p_{spin}$ < .002; Fig. S12e). From 400 to 500 ms, the positive word frequency effect in the right hemisphere showed a u-shaped association with the principal gradient, as the effect increased towards both ends of the principal gradient (interaction of the quadratic effect of the principal gradient with hemisphere: $p_{spin}$ = .032; Fig. S12f).

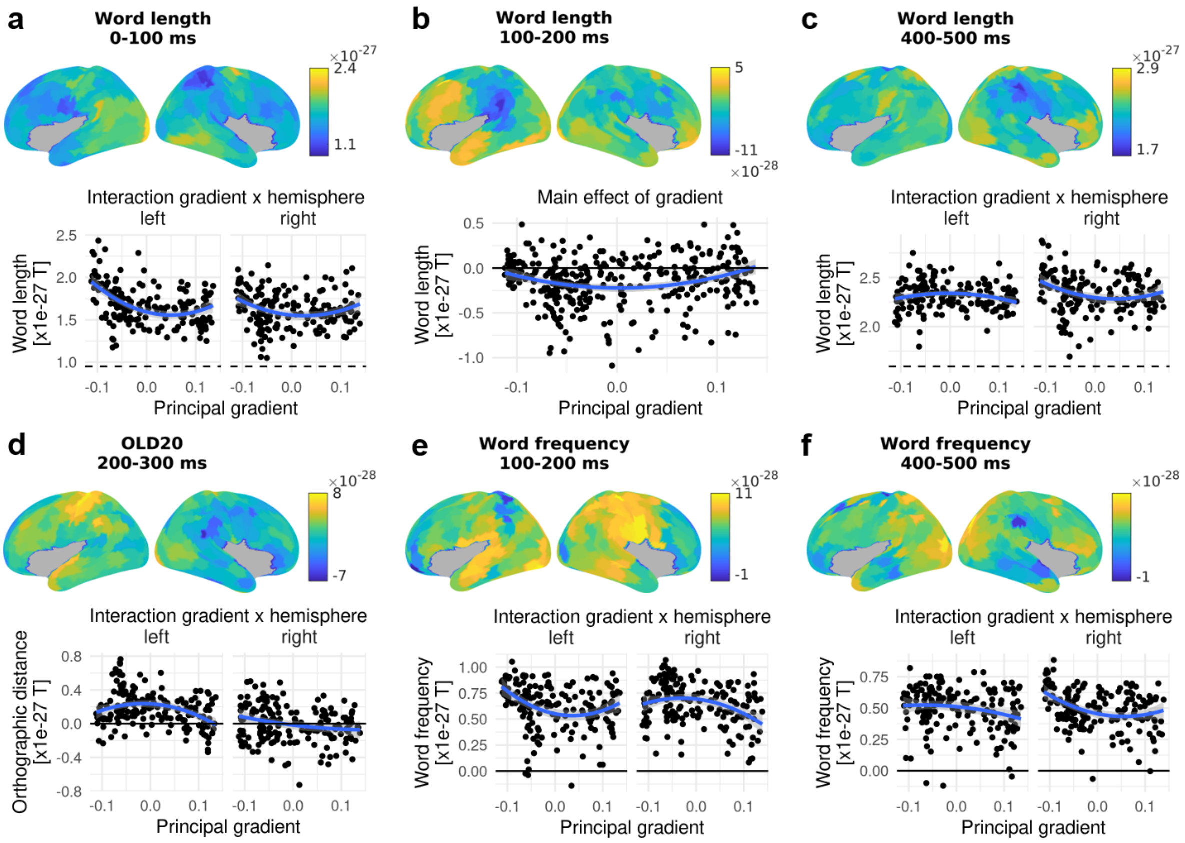


**Figure S12.** Quadratic associations with the principal connectivity gradient for word length (a-c), OLD20 (d) and word frequency effects (e-f) across the whole brain. The top panels show the effect of the respective word parameter on MEG source activation across 350 cortical parcels, indicated by the estimates from linear mixed models. Positive estimates indicate an increase in brain activation with increasing parameter values, while negative estimates indicate a decrease in brain activation with increasing parameter values. Grey parcels were not analysed. The bottom panels show the dependence of these effects on the principal gradient across parcels, i.e., the linear mixed model estimates are plotted against the principal gradient values from Margulies et al. (2016). The lowest gradient values are associated with sensory cortices, while the highest gradient values are associated with heteromodal cortices. Only the parameters and time windows which have a significant quadratic dependence on the principal gradient are shown. For parameters and time windows that only showed a significant main effect of the gradient, the bottom panels show the association between parameter estimates and gradient values combined across the left and right hemisphere, as in b. In case there was a significant interaction between gradient and hemisphere, the bottom panels show the association between parameter estimates and gradient values separately for the left and right hemisphere, as in a, c, d, e, and f.

**Supplementary results 10. Stepwise association of word frequency with the principal gradient from 200 to 300 ms in MEG data**

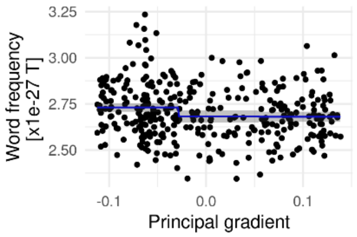


**Figure S13.** Stepwise association of the principal connectivity gradient with word frequency effects on MEG source activation in the time window from 200 to 300 ms across 350 cortical parcels. Estimates from linear mixed models are plotted against the principal gradient values from Margulies et al. (2016). The lowest gradient values are associated with sensory cortices, while the highest gradient values are associated with heteromodal cortices. Results are described in the results section of the main manuscript.

**Supplementary results 11. Gradient 2 and 3 location of word and contextual parameters' effects in MEG data**

For Gradient 2 (see Fig. 9b-f), the three-way interaction between gradient value, parameter, and hemisphere with the parameter's effect on MEG-measured brain activation as dependent variable reached significance in the time windows from 100 to 200 ms ($p_{spin}$ = .006) and 400 to 500 ms ($p_{spin}$ = .032; Tables S24 and S25; confirmed using spin permutation). Separate follow-up tests for each time window and hemisphere revealed that from 100 to 200 ms (Fig. S9c) in the left hemisphere, word length effects, which were associated with the motor end of Gradient 2, differed significantly in their association from word frequency effects, which were associated with the visual end of Gradient 2 ($p_{spin}$ = .046). In the right hemisphere, word frequency ($p_{spin}$ = .014) and OLD20 effects ($p_{spin}$ = .036), which were associated with the motor end of Gradient 2, differed significantly in their gradient association from semantic similarity effects, which did not show any tendency for an association with Gradient 2. In addition, OLD20 effects differed significantly in their Gradient 2 association from word length effects ($p_{spin}$ = .024). From 400 to 500 ms (Fig. S9f) in the left hemisphere, OLD20 effects tended more towards the visual end of Gradient 2 in contrast to semantic similarity ($p_{spin}$ = .036) and position effects ($p_{spin}$ = .05), which tended more towards the motor end. In the right hemisphere, word length effects were associated with the visual end of Gradient 2 in contrast to OLD20 ($p_{spin}$ = .028) and semantic similarity effects ($p_{spin}$ < .002), which tended towards the motor end of Gradient 2, as well as in contrast to position effects ($p_{spin}$ = .018), which did not show any tendency for an association with Gradient 2. Furthermore, word frequency effects ($p_{spin}$ = .014), which tended towards the visual end of the gradient, as well as position effects ($p_{spin}$ = .002), showed significantly different Gradient 2 associations in contrast to semantic similarity.

Differences between time windows (Tables S26 and S27) were found for word length effects ($p_{spin}$ = .011): Word length effects from 100 to 200 ms, which were stronger at the motor end (Fig. S9c), were differentially associated with Gradient 2 compared to all other time windows, in which word length effects were or tended to be stronger at the visual end (Fig. S9b,d-f; $p_{spin}$ vs. 0 to 100 ms: .014; 200 to 300 ms: .014; 300 to 400 ms: .032; 400 to 500 ms: .012). OLD20 as well showed significant changes in its association with Gradient 2 across time which differed between hemispheres (gradient by hemisphere interaction: $p_{spin}$ = .013): In the left hemisphere, OLD20 effects from 200 to 300 ms were stronger at the motor end (Fig. S9d), in contrast to the time windows 0 to 100 (Fig. S9b; $p_{spin}$ < .002) and 400 to 500 ms (Fig. S9f; $p_{spin}$ < .002), that tended to be stronger at the visual end. In the right hemisphere, OLD20 effects from 100 to 200 ms tended to be stronger at the motor end of Gradient 2 (Fig. S9c), in contrast to OLD20 effects from 0 to 100 (Fig. S9b; $p_{spin}$ = .002) and 300 to 400ms (Fig. S9e; $p_{spin}$ = .016), which tended to be stronger at the visual end.

For Gradient 3 (see Fig. S9b-f), the three-way interaction between gradient value, parameter, and hemisphere with the parameter's effect on MEG-measured brain activation as dependent variable reached significance in the time window from 100 to 200 ms ($p_{spin}$ = .006; Tables S28 and S29; confirmed using spin permutation). Separate follow-up tests for each hemisphere showed that in the left hemisphere, word frequency effects, which were associated with the DMN end of Gradient 3, differed significantly in their gradient association from word length ($p_{spin}$ < .002), OLD20 ($p_{spin}$ = .002) and semantic similarity effects ($p_{spin}$ = .002), which tended more towards the MDN end of Gradient 3, as well as from position effects ($p_{spin}$ = .016), which did not show any tendency for an association with Gradient 3 (Fig. S9c). In addition, word length effects, which were stronger at the MDN end of Gradient 3, differed significantly in their gradient association from semantic similarity ($p_{spin}$ = .01) and position effects ($p_{spin}$ = .002). No significant differences in Gradient 3 associations between any parameter pairs were found in the right hemisphere.

Differences in the association with Gradient 3 between time windows (Tables S30 and S31) were found for word length effects ($p_{spin}$ = .011), which were associated with the MDN end of Gradient 3 from 100 to 200 ms (Fig. S10c), in contrast to word length effects from 0 to 100 (Fig. S10b; $p_{spin}$ = .012) and 200 to 300 ms (Fig. S10d; $p_{spin}$ = .02), which tended to be associated with the DMN end. In addition, word length effects from 400 to 500 ms, which did not show an association with Gradient 3 (Fig. S10f), significantly differed from the association of word length effects with Gradient 3 from 0 to 100 ms (Fig. S10b; $p_{spin}$ = .026). Word frequency effects showed significant changes in their association with Gradient 3 across time which differed between hemispheres (gradient by hemisphere interaction: $p_{spin}$ = .001): In the left hemisphere, word frequency effects from 100 to 200 ms were stronger at the DMN end of Gradient 3 (Fig. S10c), in contrast to word frequency effects from 0 to 100 ($p_{spin}$ = .002), 200 to 300 ($p_{spin}$ = .04) and 400 to 500 ms ($p_{spin}$ < .002), which tended to be stronger at the MDN end (Fig. S10b,d,f). In the right hemisphere, word frequency effects from 100 to 200 ms ($p_{spin}$ = .016) and 300 to 400 ms ($p_{spin}$ = .018) tended to be stronger at the MDN end of Gradient 3 (Fig. S10c,e), in contrast to word frequency effects from 200 to 300 ms which did not show an association with the gradient (Fig. S10d).

### Supplementary results 12. Statistical tables

**Table S1.** Results of the full and winning linear model (based on stepwise model reduction using Akaike information criterion) investigating effects of linguistic parameters on self-paced reading response times

| **Full model** | | | | |
|---|---|---|---|---|
| **Parameter** | **Estimate** | **SE** | **t** | **p** |
| **Word length** | 0.013 | 0.0042 | 3.07 | 0.0022 |
| **Orthographic distance** | 0.012 | 0.0042 | 2.83 | 0.0049 |
| **Word frequency** | -0.002 | 0.0032 | -0.64 | 0.52 |
| **Semantic similarity** | -0.00028 | 0.0024 | -0.11 | 0.91 |
| **Position** | 0.01 | 0.0018 | 5.57 | 3.3e-08 |
| **Sentence order** | -0.089 | 0.0011 | -78.56 | 0 |
| **Relative clause** | 0.018 | 0.0019 | 9.51 | 1.1e-20 |
| **Verbs vs. adjectives** | -0.012 | 0.006 | -1.97 | 0.05 |
| **Verbs vs. nouns** | 0.024 | 0.0057 | 4.17 | 3.5e-05 |
| **Winning model** | | | | |
| **Parameter** | **Estimate** | **SE** | **t** | **p** |
| **Word length** | 0.013 | 0.0042 | 3.2 | 0.0014 |
| **Orthographic distance** | 0.012 | 0.004 | 3.11 | 0.002 |
| **Position** | 0.01 | 0.0018 | 5.55 | 3.6e-08 |
| **Sentence order** | -0.089 | 0.0011 | -78.56 | 0 |
| **Relative clause** | 0.018 | 0.0019 | 9.62 | 4.3e-21 |
| **Verbs vs. adjectives** | -0.012 | 0.0057 | -2.14 | 0.033 |
| **Verbs vs. nouns** | 0.024 | 0.0052 | 4.57 | 6.1e-06 |
| SE = standard error. | | | | |

**Table S2.** Results of linear models investigating effects of the principal gradient and its interaction by hemisphere on the effects of linguistic parameters on brain activation in fMRI data

| | **Principal gradient main effect** | | | | **Principal gradient by hemisphere interaction** | | | |
|---|---|---|---|---|---|---|---|---|
| **Parameter** | **Estimate** | **SE** | **t** | **p** | **Estimate** | **SE** | **t** | **p** |
| **Word length** | 3.47 | 0.97 | 3.57 | 0.00042 | 0.3 | 1.4 | 0.22 | 0.83 |
| **Orthographic distance** | -3.89 | 0.59 | -6.59 | 2.3e-10 | 0.04 | 0.85 | 0.05 | 0.96 |
| **Word frequency** | 3.72 | 0.94 | 3.98 | 8.8e-05 | -0.61 | 1.35 | -0.45 | 0.65 |
| **Semantic similarity** | -1.21 | 0.64 | -1.88 | 0.061 | -1.08 | 0.93 | -1.17 | 0.25 |
| **Position** | 2.73 | 1.57 | 1.73 | 0.084 | -3.81 | 2.28 | -1.67 | 0.095 |
| Est = estimate, SE = standard error. For p < .05, spin permutation was used to asses significance. See results section in main manuscript for significant p values from spin permutation. | | | | | | | | |

**Table S3.** Results of linear models investigating effects of the principal gradient and its interaction by hemisphere on the effects of linguistic parameters on brain activation in fMRI data when controlling for stimulus presentation duration instead of word length

| | **Principal gradient main effect** | | | | **Principal gradient by hemisphere interaction** | | | |
|---|---|---|---|---|---|---|---|---|
| **Parameter** | **Estimate** | **SE** | **t** | **p** | **Estimate** | **SE** | **t** | **p** |
| **Orthographic distance** | -3.23 | 0.62 | -5.18 | 4.5e-07 | 0.31 | 0.9 | 0.34 | 0.73 |
| **Word frequency** | 3.63 | 0.94 | 3.86 | 0.00014 | -0.61 | 1.36 | -0.45 | 0.66 |
| **Semantic similarity** | -1.26 | 0.65 | -1.93 | 0.055 | -1.31 | 0.94 | -1.38 | 0.17 |
| **Position** | 2.63 | 1.56 | 1.69 | 0.093 | -3.84 | 2.25 | -1.71 | 0.089 |
| Est = estimate, SE = standard error. For p < .05, spin permutation was used to asses significance. See results section in main manuscript for significant p values from spin permutation. | | | | | | | | |

**Table S4.** Results of the ANOVA investigating differences in linguistic parameters' associations with the principal gradient (interaction between parameters' effects on brain activation with the principal gradient and hemisphere) and post hoc linear models investigating pairwise parameter differences (interaction between parameters' effects on brain activation with the principal gradient) in fMRI data

| **ANOVA results** | | | | |
|---|---|---|---|---|
| **Parameter** | **Sum of Sqares** | **Mean Squares** | **F** | **p** |
| All (Interaction with gradient) | 43.85 | 10.96 | 14.18 | 2.5e-11 |
| All (Interaction with gradient and hemisphere) | 5.49 | 1.37 | 1.78 | 0.13 |
| **Post hoc linear model results** | | | | |
| **Parameter (Interaction with gradient)** | **Estimate** | **SE** | **t** | **p** |
| Word length vs. Orthographic distance | -0.16 | 0.84 | -0.19 | 0.85 |
| Word length vs. Word frequency | -0.18 | 0.99 | -0.18 | 0.86 |
| Word length vs. Semantic similarity | -5.5 | 0.85 | -6.5 | 1.9e-10 |
| Word length vs. Position | 4.74 | 1.37 | 3.47 | 0.00057 |
| Orthographic distance vs. Word frequency | -0.34 | 0.82 | -0.41 | 0.68 |
| Orthographic distance vs. | -5.66 | 0.63 | -8.95 | 6e-18 |

| | | | | |
|---|---|---|---|---|
| Semantic similarity | | | | |
| Orthographic distance vs. Position | 4.9 | 1.25 | 3.93 | 9.8e-05 |
| Word frequency vs. Semantic similarity | -5.32 | 0.82 | -6.45 | 2.5e-10 |
| Word frequency vs. Position | 4.56 | 1.35 | 3.37 | 0.00081 |
| Semantic similarity vs. Position | -0.76 | 1.25 | -0.61 | 0.54 |
| SE = standard error. For F or p < .05, spin permutation was used to asses significance. See results section in main manuscript for significant p values from spin permutation. | | | | |

**Table S5.** Results of the ANOVA investigating differences in linguistic parameters' associations with the principal gradient (interaction between parameters' effects on brain activation with the principal gradient and hemisphere) and follow-up linear models investigating pairwise parameter differences (interaction between parameters' effects on brain activation with the principal gradient) in fMRI data when controlling for stimulus presentation duration instead of word length

| **ANOVA results** | | | | |
|---|---|---|---|---|
| **Parameter** | **Sum of Sqares** | **Mean Squares** | **F** | **p** |
| All (Interaction with gradient) | 32.81 | 8.2 | 10.61 | 1.8e-08 |
| All (Interaction with gradient and hemisphere) | 6.02 | 1.51 | 1.95 | 0.1 |
| **Follow-up linear model results** | | | | |
| **Parameter (Interaction with gradient)** | **Estimate** | **SE** | **t** | **p** |
| Orthographic distance vs. Word frequency | 0.26 | 0.82 | 0.31 | 0.75 |
| Orthographic distance vs. Semantic similarity | -4.93 | 0.65 | -7.54 | 2e-13 |
| Orthographic distance vs. Position | 3.91 | 1.22 | 3.2 | 0.0015 |
| Word frequency vs. Semantic similarity | -5.18 | 0.83 | -6.25 | 8.2e-10 |
| Word frequency vs. Position | 4.17 | 1.33 | 3.14 | 0.0018 |

| Semantic similarity vs. Position | -1.01 | 1.23 | -0.82 | 0.41 |
|---|---|---|---|---|
| SE = standard error. For F or p < .05, spin permutation was used to asses significance. See results section in main manuscript for significant p values from spin permutation. | | | | |

**Table S6.** Results of linear models investigating effects of the principal gradient and its interaction by hemisphere on the effects of linguistic parameters on brain activation in MEG data.

| | | Principal gradient main effect | | | | Principal gradient by hemisphere interaction | | | |
|---|---|---|---|---|---|---|---|---|---|
| **Parameter** | **Time [ms]** | **Est** | **SE** | **t** | **p** | **Est** | **SE** | **t** | **p** |
| **Word length** | 0-100 | -11.04 | 2.13 | -5.19 | 3.5e-07 | 7.73 | 3.04 | 2.54 | 0.011 |
| | 100-200 | 4.89 | 2.66 | 1.84 | 0.067 | -5.43 | 3.8 | -1.43 | 0.15 |
| | 200-300 | 2.93 | 1.77 | 1.66 | 0.098 | -1.33 | 2.53 | -0.53 | 0.6 |
| | 300-400 | -4.04 | 1.85 | -2.18 | 0.03 | -1.86 | 2.65 | -0.7 | 0.48 |
| | 400-500 | -1.58 | 1.69 | -0.94 | 0.35 | -3.24 | 2.41 | -1.35 | 0.18 |
| **Orthographic distance** | 0-100 | 9.13 | 1.76 | 5.2 | 3.5e-07 | -6.66 | 2.51 | -2.65 | 0.0084 |
| | 100-200 | -6.85 | 3.28 | -2.09 | 0.037 | 2.56 | 4.68 | 0.55 | 0.58 |
| | 200-300 | -6.19 | 2.03 | -3.05 | 0.0025 | -0.35 | 2.9 | -0.12 | 0.9 |
| | 300-400 | 6.84 | 1.89 | 3.61 | 0.00035 | 1.46 | 2.71 | 0.54 | 0.59 |
| | 400-500 | 2.05 | 2.06 | 1 | 0.32 | 2.38 | 2.94 | 0.81 | 0.42 |
| **Word frequency** | 0-100 | -8.4 | 1.74 | -4.83 | 2e-06 | 11.89 | 2.49 | 4.78 | 2.6e-06 |
| | 100-200 | -6.17 | 2.07 | -2.98 | 0.0031 | -0.95 | 2.96 | -0.32 | 0.75 |
| | 200-300 | -4.31 | 1.26 | -3.42 | 0.00069 | 1.62 | 1.8 | 0.9 | 0.37 |
| | 300-400 | -0.44 | 1.29 | -0.34 | 0.74 | -3.35 | 1.84 | -1.81 | 0.07 |
| | 400-500 | -4.32 | 1.58 | -2.73 | 0.0067 | -1.97 | 2.27 | -0.87 | 0.38 |
| **Semantic similarity** | 0-100 | -7.49 | 1.33 | -5.65 | 3.4e-08 | 6.4 | 1.9 | 3.38 | 0.00081 |
| | 100-200 | -0.97 | 1.26 | -0.77 | 0.44 | -1.56 | 1.8 | -0.87 | 0.39 |
| | 200-300 | -3.11 | 1 | -3.12 | 0.002 | 3.48 | 1.42 | 2.44 | 0.015 |
| | 300-400 | 0.37 | 1.04 | 0.36 | 0.72 | -2.07 | 1.48 | -1.39 | 0.17 |
| | 400-500 | -1.69 | 1.11 | -1.53 | 0.13 | -0.03 | 1.59 | -0.02 | 0.99 |

| | | | | | | | | | |
|---|---|---|---|---|---|---|---|---|---|
| **Position** | 0-100 | -0.16 | 1.33 | -0.12 | 0.9 | -0.09 | 1.9 | -0.05 | 0.96 |
| | 100-200 | -2.6 | 1.01 | -2.58 | 0.01 | -1.55 | 1.44 | -1.07 | 0.28 |
| | 200-300 | -0.3 | 1.07 | -0.28 | 0.78 | 5.24 | 1.53 | 3.42 | 0.00069 |
| | 300-400 | -0.5 | 1.16 | -0.43 | 0.67 | 3.8 | 1.66 | 2.29 | 0.023 |
| | 400-500 | 2.81 | 0.98 | 2.88 | 0.0042 | -0.83 | 1.4 | -0.59 | 0.55 |
| Est = estimate, SE = standard error; in e-28 T. For p < .05, spin permutation was used to asses significance. See results section in main manuscript for significant p values from spin permutation. | | | | | | | | | |

**Table S7.** Results of linear models investigating effects of the principal gradient and its interaction by hemisphere on the effects of linguistic parameters on brain activation in MEG data when controlling for stimulus presentation duration instead of word length

| | | **Principal gradient main effect** | | | | **Principal gradient by hemisphere interaction** | | | |
|---|---|---|---|---|---|---|---|---|---|
| **Parameter** | **Time [ms]** | **Est** | **SE** | **t** | **p** | **Est** | **SE** | **t** | **p** |
| **Orthographic distance** | 0-100 | 7.67 | 1.39 | 5.53 | 6.5e-08 | -2.7 | 1.98 | -1.36 | 0.17 |
| | 100-200 | -5.96 | 2.78 | -2.14 | 0.033 | 3.42 | 3.97 | 0.86 | 0.39 |
| | 200-300 | -3.68 | 1.42 | -2.59 | 0.0099 | -0.93 | 2.03 | -0.46 | 0.65 |
| | 300-400 | 6.92 | 1.82 | 3.81 | 0.00017 | 1 | 2.6 | 0.39 | 0.7 |
| | 400-500 | 1.22 | 2.21 | 0.55 | 0.58 | 2.14 | 3.17 | 0.68 | 0.5 |
| **Word frequency** | 0-100 | -8.76 | 1.82 | -4.82 | 2.1e-06 | 10.97 | 2.6 | 4.22 | 3.1e-05 |
| | 100-200 | -6.1 | 2.04 | -2.99 | 0.003 | -1.77 | 2.92 | -0.61 | 0.54 |
| | 200-300 | -5.08 | 1.37 | -3.72 | 0.00023 | 1.74 | 1.95 | 0.89 | 0.37 |
| | 300-400 | -0.83 | 1.35 | -0.61 | 0.54 | -3.32 | 1.93 | -1.72 | 0.086 |
| | 400-500 | -4.12 | 1.47 | -2.8 | 0.0055 | -2.16 | 2.1 | -1.03 | 0.3 |
| **Semantic similarity** | 0-100 | -7.46 | 1.34 | -5.58 | 4.8e-08 | 6.26 | 1.91 | 3.27 | 0.0012 |
| | 100-200 | -0.99 | 1.27 | -0.78 | 0.44 | -1.62 | 1.82 | -0.89 | 0.37 |
| | 200-300 | -3.2 | 1 | -3.21 | 0.0014 | 3.5 | 1.42 | 2.46 | 0.015 |
| | 300-400 | 0.35 | 1.05 | 0.34 | 0.74 | -2.05 | 1.49 | -1.37 | 0.17 |
| | 400-500 | -1.66 | 1.11 | -1.5 | 0.13 | -0.03 | 1.58 | -0.02 | 0.98 |
| **Position** | 0-100 | 0.98 | 1.36 | 0.72 | 0.47 | -0.29 | 1.94 | -0.15 | 0.88 |
| | 100-200 | -3.04 | 1.05 | -2.91 | 0.0038 | -0.69 | 1.49 | -0.46 | 0.64 |

| | | | | | | | | | |
|---|---|---|---|---|---|---|---|---|---|
| | 200-300 | -0.18 | 1.08 | -0.17 | 0.87 | 5.3 | 1.54 | 3.44 | 0.00066 |
| | 300-400 | 0.01 | 1.15 | 0.01 | 0.99 | 3.94 | 1.64 | 2.4 | 0.017 |
| | 400-500 | 2.85 | 0.97 | 2.94 | 0.0035 | -0.49 | 1.38 | -0.35 | 0.72 |
| Est = estimate, SE = standard error; in e-28 T. For p < .05, spin permutation was used to asses significance. See results section in main manuscript for significant p values from spin permutation. | | | | | | | | | |

**Table S8.** Results of the ANOVA investigating differences in linguistic parameters' associations with the principal gradient (interaction between parameters' effects on brain activation with the principal gradient and hemisphere) and follow-up linear models investigating pairwise parameter differences (interaction between parameters' effects on brain activation with the principal gradient) in MEG data

| **ANOVA results** | | | | | |
|---|---|---|---|---|---|
| **Parameter** | **Time [ms]** | **Sum of Sqares** | **Mean Squares** | **F** | **p** |
| All (Interaction with gradient) | 0-100 | 51.17 | 5.69 | 44.48 | 3.8e-76 |
| | 100-200 | 56.87 | 6.32 | 55.53 | 5.7e-95 |
| | 200-300 | 25.87 | 2.87 | 32.46 | 3.1e-55 |
| | 300-400 | 29.33 | 3.26 | 42.05 | 6e-72 |
| | 400-500 | 22.61 | 2.51 | 31 | 1.2e-52 |
| All (Interaction with gradient and hemisphere) | 0-100 | 2.23 | 0.25 | 1.94 | 0.042 |
| | 100-200 | 1.04 | 0.12 | 1.02 | 0.42 |
| | 200-300 | 0.89 | 0.1 | 1.12 | 0.35 |
| | 300-400 | 1.52 | 0.17 | 2.18 | 0.021 |
| | 400-500 | 0.58 | 0.06 | 0.79 | 0.62 |
| **Follow-up linear model results** | | | | | |
| **Parameter** | **Time [ms]** | **Estimate** | **SE** | **t** | **p** |
| Word length vs. Orthographic distance | 0-100 | 1.41 | 2.01 | 0.7 | 0.49 |
| | 100-200 | -3.38 | 3.01 | -1.12 | 0.26 |
| | 200-300 | -8.28 | 2.03 | -4.07 | 5.2e-05 |
| | 300-400 | -2.55 | 1.93 | -1.32 | 0.19 |
| | 400-500 | 6.43 | 1.9 | 3.39 | 0.00075 |
| Word length vs. Word frequency | 0-100 | -4.68 | 2.02 | -2.32 | 0.021 |
| | 100-200 | 4.41 | 2.41 | 1.83 | 0.067 |
| | 200-300 | 5.48 | 1.65 | 3.32 | 0.00095 |
| | 300-400 | -7.01 | 1.63 | -4.29 | 2e-05 |
| | 400-500 | 2.07 | 1.65 | 1.25 | 0.21 |
| Word length vs. Semantic similarity | 0-100 | -11.36 | 1.85 | -6.12 | 1.5e-09 |
| | 100-200 | -0.52 | 2.1 | -0.25 | 0.8 |
| | 200-300 | 0.77 | 1.47 | 0.53 | 0.6 |
| | 300-400 | -4.17 | 1.53 | -2.73 | 0.0066 |
| | 400-500 | -5 | 1.46 | -3.43 | 0.00064 |
| Word length vs. Position | 0-100 | 6.91 | 1.81 | 3.81 | 0.00015 |
| | 100-200 | -1.15 | 2.03 | -0.57 | 0.57 |
| | 200-300 | 0.04 | 1.5 | 0.02 | 0.98 |
| | 300-400 | 6.37 | 1.61 | 3.95 | 8.5e-05 |

| | | | | | |
|---|---|---|---|---|---|
| | 400-500 | 5.62 | 1.39 | 4.03 | 6.1e-05 |
| Orthographic distance vs. Word frequency | 0-100 | -3.27 | 1.83 | -1.78 | 0.075 |
| | 100-200 | 1.03 | 2.76 | 0.37 | 0.71 |
| | 200-300 | -2.8 | 1.89 | -1.48 | 0.14 |
| | 300-400 | -9.56 | 1.68 | -5.69 | 1.8e-08 |
| | 400-500 | 8.51 | 1.85 | 4.59 | 5.2e-06 |
| Orthographic distance vs. Semantic similarity | 0-100 | -9.95 | 1.65 | -6.03 | 2.7e-09 |
| | 100-200 | -3.91 | 2.5 | -1.56 | 0.12 |
| | 200-300 | -7.5 | 1.73 | -4.33 | 1.7e-05 |
| | 300-400 | -6.72 | 1.58 | -4.25 | 2.4e-05 |
| | 400-500 | 1.44 | 1.68 | 0.86 | 0.39 |
| Orthographic distance vs. Position | 0-100 | 5.51 | 1.61 | 3.43 | 0.00064 |
| | 100-200 | 2.23 | 2.45 | 0.91 | 0.36 |
| | 200-300 | 8.31 | 1.76 | 4.72 | 2.9e-06 |
| | 300-400 | 8.92 | 1.66 | 5.38 | 1e-07 |
| | 400-500 | -0.81 | 1.62 | -0.5 | 0.62 |
| Word frequency vs. Semantic similarity | 0-100 | -6.68 | 1.66 | -4.03 | 6.2e-05 |
| | 100-200 | -4.93 | 1.73 | -2.86 | 0.0044 |
| | 200-300 | -4.71 | 1.26 | -3.73 | 0.00021 |
| | 300-400 | 2.84 | 1.2 | 2.36 | 0.019 |
| | 400-500 | -7.07 | 1.39 | -5.08 | 4.9e-07 |
| Word frequency vs. Position | 0-100 | 2.23 | 1.61 | 1.38 | 0.17 |
| | 100-200 | 3.26 | 1.65 | 1.98 | 0.048 |
| | 200-300 | 5.51 | 1.3 | 4.24 | 2.6e-05 |
| | 300-400 | -0.64 | 1.3 | -0.49 | 0.62 |
| | 400-500 | 7.69 | 1.33 | 5.8 | 1e-08 |
| Semantic similarity vs. Position | 0-100 | -4.45 | 1.4 | -3.18 | 0.0016 |
| | 100-200 | -1.68 | 1.16 | -1.45 | 0.15 |
| | 200-300 | 0.81 | 1.06 | 0.76 | 0.45 |
| | 300-400 | 2.19 | 1.17 | 1.87 | 0.062 |
| | 400-500 | 0.62 | 1.07 | 0.58 | 0.56 |
| SE = standard error. Sum of squares and mean squares in e-54 T. Estimate and SE in e-28 T. For F or p < .05, spin permutation was used to asses significance. See results section in main manuscript for significant p values from spin permutation. | | | | | |

**Table S9.** Results of the ANOVA investigating differences in linguistic parameters' associations with the principal gradient (interaction between parameters' effects on brain activation with the principal gradient and hemisphere) and follow-up linear models investigating pairwise parameter differences (interaction between parameters' effects on brain activation with the principal gradient) in MEG data when controlling for stimulus presentation duration instead of word length

| **ANOVA results** | | | | | |
|---|---|---|---|---|---|
| **Parameter** | **Time [ms]** | **Sum of Sqares** | **Mean Squares** | **F** | **p** |
| All (Interaction with gradient) | 0-100 | 51.55 | 5.73 | 45.23 | 2e-77 |
| | 100-200 | 58.4 | 6.49 | 60.01 | 1.8e-102 |
| | 200-300 | 26.04 | 2.89 | 33.92 | 8.7e-58 |
| | 300-400 | 29.43 | 3.27 | 42.94 | 1.7e-73 |
| | 400-500 | 22.07 | 2.45 | 29.85 | 1.2e-50 |
| All (Interaction with gradient and hemisphere) | 0-100 | 1.5 | 0.17 | 1.32 | 0.22 |
| | 100-200 | 1.28 | 0.14 | 1.31 | 0.23 |
| | 200-300 | 0.93 | 0.1 | 1.21 | 0.29 |
| | 300-400 | 1.47 | 0.16 | 2.15 | 0.023 |
| | 400-500 | 0.58 | 0.06 | 0.78 | 0.64 |
| **Follow-up linear model results** | | | | | |
| **Parameter** | **Time [ms]** | **Estimate** | **SE** | **t** | **p** |
| Orthographic distance vs. Word frequency | 0-100 | -2.88 | 1.76 | -1.63 | 0.1 |
| | 100-200 | 2.78 | 2.47 | 1.13 | 0.26 |
| | 200-300 | -0.04 | 1.57 | -0.03 | 0.98 |
| | 300-400 | -9.88 | 1.64 | -6.02 | 2.9e-09 |
| | 400-500 | 7.45 | 1.9 | 3.93 | 9.4e-05 |
| Orthographic distance vs. Semantic similarity | 0-100 | -10.37 | 1.52 | -6.84 | 1.8e-11 |
| | 100-200 | -2.47 | 2.18 | -1.13 | 0.26 |
| | 200-300 | -5.5 | 1.28 | -4.31 | 1.9e-05 |
| | 300-400 | -6.63 | 1.51 | -4.38 | 1.4e-05 |
| | 400-500 | 0.52 | 1.78 | 0.29 | 0.77 |
| Orthographic distance vs. Position | 0-100 | 6.89 | 1.48 | 4.66 | 3.9e-06 |
| | 100-200 | 0.84 | 2.12 | 0.39 | 0.69 |
| | 200-300 | 6.35 | 1.32 | 4.82 | 1.8e-06 |
| | 300-400 | 9.42 | 1.59 | 5.94 | 4.6e-09 |
| | 400-500 | 0.34 | 1.72 | 0.2 | 0.85 |
| Word frequency vs. Semantic similarity | 0-100 | -7.49 | 1.72 | -4.35 | 1.5e-05 |
| | 100-200 | -5.25 | 1.72 | -3.05 | 0.0024 |
| | 200-300 | -5.45 | 1.37 | -3.99 | 7.3e-05 |
| | 300-400 | 3.25 | 1.25 | 2.61 | 0.0092 |
| | 400-500 | -6.94 | 1.33 | -5.23 | 2.3e-07 |
| Word frequency vs. Position | 0-100 | 4.01 | 1.69 | 2.38 | 0.018 |
| | 100-200 | 3.61 | 1.64 | 2.2 | 0.028 |
| | 200-300 | 6.31 | 1.41 | 4.48 | 8.6e-06 |
| | 300-400 | -0.46 | 1.33 | -0.35 | 0.73 |
| | 400-500 | 7.79 | 1.26 | 6.2 | 9.9e-10 |
| | 0-100 | -3.48 | 1.43 | -2.43 | 0.015 |
| | 100-200 | -1.63 | 1.18 | -1.39 | 0.17 |

| | | | | | |
|---|---|---|---|---|---|
| Semantic similarity vs. Position | 200-300 | 0.86 | 1.07 | 0.8 | 0.42 |
| | 300-400 | 2.79 | 1.17 | 2.38 | 0.017 |
| | 400-500 | 0.86 | 1.07 | 0.8 | 0.42 |
| SE = standard error. Sum of squares and mean squares in e-54 T. Estimate and SE in e-28 T. For F or p < .05, spin permutation was used to asses significance. See results section in main manuscript for significant p values from spin permutation. | | | | | |

**Table S10.** Results of the ANOVA investigating differences in each linguistic parameter's association with the principal gradient across time (interaction between time, principal gradient and hemisphere) and follow-up linear models investigating pairwise time window differences for word frequency effects (interaction between time and principal gradient) in MEG data

| **ANOVA results** | | | | | |
|---|---|---|---|---|---|
| **Parameter** | **Effect** | **Sum of Squares** | **Mean Squares** | **F** | **p** |
| Word length | Principal gradient x Time | 0.95 | 0.24 | 5.63 | 0.00017 |
| | Principal gradient x Time x Hemisphere | 0.47 | 0.12 | 2.8 | 0.025 |
| Orthographic distance | Principal gradient x Time | 1.45 | 0.36 | 6.98 | 1.4e-05 |
| | Principal gradient x Time x Hemisphere | 0.2 | 0.05 | 0.94 | 0.44 |
| Word frequency | Principal gradient x Time | 0.9 | 0.22 | 8.52 | 8.3e-07 |
| | Principal gradient x Time x Hemisphere | 0.6 | 0.15 | 5.68 | 0.00015 |
| Semantic similarity | Principal gradient x Time | 0.42 | 0.11 | 7.85 | 2.9e-06 |
| | Principal gradient x Time x Hemisphere | 0.12 | 0.03 | 2.16 | 0.071 |
| Position | Principal gradient x Time | 0.46 | 0.12 | 9.22 | 2.3e-07 |
| | Principal gradient x | 0.18 | 0.04 | 3.57 | 0.0066 |

| | | | | | |
|---|---|---|---|---|---|
| | Time x Hemisphere | | | | |
| **Follow-up linear model results** | | | | | |
| **Parameter** | **Time** | **Estimate** | **SE** | **t** | **p** |
| Word frequency | 0-100 vs. 100-200 | -4.18 | 1.97 | -2.13 | 0.034 |
| | 0-100 vs. 200-300 | -0.81 | 1.67 | -0.49 | 0.63 |
| | 0-100 vs. 300-400 | 4.63 | 1.61 | 2.89 | 0.004 |
| | 0-100 vs. 400-500 | -2.8 | 1.72 | -1.62 | 0.1 |
| | 100-200 vs. 200-300 | 3.37 | 1.81 | 1.87 | 0.062 |
| | 100-200 vs. 300-400 | 8.81 | 1.75 | 5.04 | 6e-07 |
| | 100-200 vs. 400-500 | 1.38 | 1.86 | 0.75 | 0.46 |
| | 200-300 vs. 300-400 | 5.44 | 1.4 | 3.88 | 0.00011 |
| | 200-300 vs. 400-500 | -1.99 | 1.54 | -1.3 | 0.2 |
| | 300-400 vs. 400-500 | -7.43 | 1.47 | -5.05 | 5.5e-07 |
| SE = standard error. Sum of squares and mean squares in e-54 T. Estimate and SE in e-28 T. For F or p < .05, spin permutation was used to asses significance. See results section in main manuscript for significant p values from spin permutation. | | | | | |

**Table S11.** Results of the ANOVA investigating differences in each linguistic parameter's association with the principal gradient across time (interaction between time, principal gradient and hemisphere) and follow-up linear models investigating pairwise time window differences for word frequency effects (interaction between time and principal gradient) in MEG data when controlling for stimulus presentation duration instead of word length

| **ANOVA results** | | | | | |
|---|---|---|---|---|---|
| **Parameter** | **Effect** | **Sum of Squares** | **Mean Squares** | **F** | **p** |
| Orthographic distance | Principal gradient x Time | 1.09 | 0.27 | 6.79 | 2e-05 |
| | Principal gradient x Time x Hemisphere | 0.09 | 0.02 | 0.54 | 0.71 |
| Word frequency | Principal gradient x Time | 1.03 | 0.26 | 9.56 | 1.2e-07 |
| | Principal gradient x | 0.56 | 0.14 | 5.18 | 0.00038 |

| | | | | | |
|---|---|---|---|---|---|
| | Time x Hemisphere | | | | |
| Semantic similarity | Principal gradient x Time | 0.43 | 0.11 | 8.01 | 2.1e-06 |
| | Principal gradient x Time x Hemisphere | 0.11 | 0.03 | 2.03 | 0.087 |
| Position | Principal gradient x Time | 0.5 | 0.12 | 9.69 | 9.5e-08 |
| | Principal gradient x Time x Hemisphere | 0.16 | 0.04 | 3.11 | 0.015 |
| **Follow-up linear model results** | | | | | |
| **Parameter** | **Time** | **Estimate** | **SE** | **t** | **p** |
| Word frequency | 0-100 vs. 100-200 | -3.78 | 2.01 | -1.88 | 0.06 |
| | 0-100 vs. 200-300 | -0.71 | 1.8 | -0.39 | 0.7 |
| | 0-100 vs. 300-400 | 5.8 | 1.69 | 3.43 | 0.00064 |
| | 0-100 vs. 400-500 | -1.92 | 1.73 | -1.11 | 0.27 |
| | 100-200 vs. 200-300 | 3.08 | 1.87 | 1.64 | 0.1 |
| | 100-200 vs. 300-400 | 9.58 | 1.77 | 5.42 | 8.1e-08 |
| | 100-200 vs. 400-500 | 1.86 | 1.8 | 1.03 | 0.3 |
| | 200-300 vs. 300-400 | 6.5 | 1.53 | 4.26 | 2.4e-05 |
| | 200-300 vs. 400-500 | -1.22 | 1.57 | -0.78 | 0.44 |
| | 300-400 vs. 400-500 | -7.72 | 1.44 | -5.35 | 1.2e-07 |
| SE = standard error. Sum of squares and mean squares in e-54 T. Estimate and SE in e-28 T. For F or p < .05, spin permutation was used to asses significance. See results section in main manuscript for significant p values from spin permutation. | | | | | |

**Table S12.** Results of linear models investigating effects of Gradient 2 and its interaction by hemisphere on the effects of linguistic parameters on brain activation in fMRI data

| | **Gradient 2 main effect** | | | | **Gradient 2 by hemisphere interaction** | | | |
|---|---|---|---|---|---|---|---|---|
| **Parameter** | **Est** | **SE** | **t** | **p** | **Est** | **SE** | **t** | **p** |
| **Word length** | -1.4 | 1.1 | -1.27 | 0.21 | 2.5 | 1.59 | 1.57 | 0.12 |
| **Orthographic distance** | -0.24 | 0.73 | -0.33 | 0.74 | -0.64 | 1.06 | -0.61 | 0.55 |
| **Word frequency** | 0.05 | 1.07 | 0.05 | 0.96 | -0.27 | 1.54 | -0.17 | 0.86 |
| **Semantic similarity** | 0.62 | 0.72 | 0.87 | 0.39 | 0.05 | 1.03 | 0.05 | 0.96 |
| **Position** | 0.95 | 1.72 | 0.55 | 0.58 | -0.17 | 2.49 | -0.07 | 0.95 |
| Est = estimate, SE = standard error. For p < .05, spin permutation was used to asses significance. See text for significant p values from spin permutation. | | | | | | | | |

**Table S13.** Results of linear models investigating effects of Gradient 2 and its interaction by hemisphere on the effects of linguistic parameters on brain activation in fMRI data when controlling for stimulus presentation duration instead of word length

| | **Gradient 2 main effect** | | | | **Gradient 2 by hemisphere interaction** | | | |
|---|---|---|---|---|---|---|---|---|
| **Parameter** | **Est** | **SE** | **t** | **p** | **Est** | **SE** | **t** | **p** |
| **Orthographic distance** | -2.74 | 0.71 | -3.86 | 0.00014 | 1.1 | 1.03 | 1.07 | 0.29 |
| **Word frequency** | 0.63 | 1.07 | 0.59 | 0.56 | -0.74 | 1.55 | -0.48 | 0.63 |
| **Semantic similarity** | 0.92 | 0.73 | 1.26 | 0.21 | -0.06 | 1.05 | -0.05 | 0.96 |
| **Position** | 0.72 | 1.7 | 0.42 | 0.67 | -0.28 | 2.46 | -0.11 | 0.91 |
| Est = estimate, SE = standard error. For p < .05, spin permutation was used to asses significance. See text for significant p values from spin permutation. | | | | | | | | |

**Table S14.** Results of linear models investigating effects of Gradient 3 and its interaction by hemisphere on the effects of linguistic parameters on brain activation in fMRI data

| | **Gradient 3 main effect** | | | | **Gradient 3 by hemisphere interaction** | | | |
|---|---|---|---|---|---|---|---|---|
| **Parameter** | **Est** | **SE** | **t** | **p** | **Est** | **SE** | **t** | **p** |
| **Word length** | -9.77 | 1.38 | -7.1 | 1.2e-11 | 1.91 | 1.96 | 0.97 | 0.33 |
| **Orthographic distance** | 3.16 | 1.01 | 3.12 | 0.002 | -0.52 | 1.45 | -0.36 | 0.72 |
| **Word frequency** | -7.49 | 1.41 | -5.32 | 2.2e-07 | 2.29 | 2.01 | 1.14 | 0.26 |
| **Semantic similarity** | 5.38 | 0.97 | 5.57 | 6.1e-08 | -5.32 | 1.38 | -3.87 | 0.00014 |
| **Position** | 0.86 | 2.44 | 0.35 | 0.73 | 1.78 | 3.49 | 0.51 | 0.61 |

| Est = estimate, SE = standard error. For p < .05, spin permutation was used to asses significance. See text for significant p values from spin permutation. |
|---|

**Table S15.** Results of linear models investigating effects of Gradient 3 and its interaction by hemisphere on the effects of linguistic parameters on brain activation in fMRI data when controlling for stimulus presentation duration instead of word length

| | **Gradient 3 main effect** | | | | **Gradient 3 by hemisphere interaction** | | | |
|---|---|---|---|---|---|---|---|---|
| **Parameter** | **Est** | **SE** | **t** | **p** | **Est** | **SE** | **t** | **p** |
| **Orthographic distance** | -3.18 | 1.02 | -3.11 | 0.0021 | 1.58 | 1.46 | 1.08 | 0.28 |
| **Word frequency** | -5.82 | 1.45 | -4.01 | 8e-05 | 1.35 | 2.07 | 0.65 | 0.51 |
| **Semantic similarity** | 6.09 | 0.97 | 6.27 | 1.5e-09 | -5.77 | 1.39 | -4.16 | 4.3e-05 |
| **Position** | 1.33 | 2.41 | 0.55 | 0.58 | 1.49 | 3.45 | 0.43 | 0.67 |
| Est = estimate, SE = standard error. For p < .05, spin permutation was used to asses significance. See text for significant p values from spin permutation. | | | | | | | | |

**Table S16.** Results of the ANOVA investigating differences in linguistic parameters' associations with Gradient 2 (interaction between parameters' effects on brain activation with Gradient 2 and hemisphere) in fMRI data

| **ANOVA results** | | | | |
|---|---|---|---|---|
| **Parameter** | **Sum of Sqares** | **Mean Squares** | **F** | **p** |
| All (Interaction with gradient) | 1.86 | 0.47 | 0.57 | 0.69 |
| All (Interaction with gradient and hemisphere) | 1.5 | 0.38 | 0.46 | 0.77 |
| For F or p < .05, spin permutation was used to asses significance. See results section in main manuscript for significant values from spin permutation. | | | | |

**Table S17.** Results of the ANOVA investigating differences in linguistic parameters' associations with Gradient 2 (interaction between parameters' effects on brain activation with Gradient 2 and hemisphere) in fMRI data when controlling for stimulus presentation duration instead of word length

| **ANOVA results** | | | | |
|---|---|---|---|---|
| **Parameter** | **Sum of Sqares** | **Mean Squares** | **F** | **p** |
| All (Interaction with gradient) | 8.84 | 2.21 | 2.77 | 0.026 |
| All (Interaction with gradient and hemisphere) | 0.58 | 0.15 | 0.18 | 0.95 |
| For F or p < .05, spin permutation was used to asses significance. See text for significant p values from spin permutation. | | | | |

**Table S18.** Results of the ANOVA investigating differences in linguistic parameters' associations with Gradient 3 (interaction between parameters' effects on brain activation with Gradient 3 and hemisphere) and post hoc linear models investigating pairwise parameter differences (interaction between parameters' effects on brain activation with Gradient 3) in fMRI data

| ANOVA results | | | | |
|---|---|---|---|---|
| **Parameter** | **Sum of Sqares** | **Mean Squares** | **F** | **p** |
| All (Interaction with gradient) | 48.92 | 12.23 | 16.18 | 6e-13 |
| All (Interaction with gradient and hemisphere) | 6.21 | 1.55 | 2.05 | 0.085 |
| **Post hoc linear model results** | | | | |
| **Parameter (Interaction with gradient)** | **Estimate** | **SE** | **t** | **p** |
| Word length vs. Orthographic distance | -5.91 | 1.22 | -4.86 | 1.6e-06 |
| Word length vs. Word frequency | 2.52 | 1.41 | 1.79 | 0.073 |
| Word length vs. Semantic similarity | 11.37 | 1.21 | 9.42 | 1.3e-19 |
| Word length vs. Position | -7.33 | 2 | -3.66 | 0.00028 |
| Orthographic distance vs. Word frequency | -3.39 | 1.24 | -2.74 | 0.0064 |
| Orthographic distance vs. Semantic similarity | 5.46 | 1.01 | 5.41 | 9.8e-08 |
| Orthographic distance vs. Position | -1.42 | 1.89 | -0.75 | 0.45 |
| Word frequency vs. Semantic similarity | 8.85 | 1.23 | 7.2 | 2.1e-12 |
| Word frequency vs. Position | -4.81 | 2.02 | -2.39 | 0.017 |
| Semantic similarity vs. Position | 4.04 | 1.88 | 2.14 | 0.032 |
| SE = standard error. For F or p < .05, spin permutation was used to asses significance. See text for significant p values from spin permutation. | | | | |

**Table S19.** Results of the ANOVA investigating differences in linguistic parameters' associations with Gradient 3 (interaction between parameters' effects on brain activation with Gradient 3 and hemisphere) and follow-up linear models investigating pairwise parameter differences (interaction between parameters' effects on brain activation with Gradient 3) in fMRI data when controlling for stimulus presentation duration instead of word length

| **ANOVA results** | | | | |
|---|---|---|---|---|
| **Parameter** | **Sum of Sqares** | **Mean Squares** | **F** | **p** |
| All (Interaction with gradient) | 31.35 | 7.84 | 10.07 | 5e-08 |
| All (Interaction with gradient and hemisphere) | 4.26 | 1.07 | 1.37 | 0.24 |
| **Follow-up linear model results** | | | | |
| **Parameter (Interaction with gradient)** | **Estimate** | **SE** | **t** | **p** |
| Orthographic distance vs. Word frequency | -7.47 | 1.27 | -5.9 | 6.6e-09 |
| Orthographic distance vs. Semantic similarity | 0.67 | 1.02 | 0.66 | 0.51 |
| Orthographic distance vs. Position | 4.16 | 1.88 | 2.22 | 0.027 |
| Word frequency vs. Semantic similarity | 8.14 | 1.26 | 6.47 | 2.2e-10 |
| Word frequency vs. Position | -3.31 | 2.02 | -1.64 | 0.1 |
| Semantic similarity vs. Position | 4.83 | 1.87 | 2.58 | 0.01 |
| SE = standard error. For F or p < .05, spin permutation was used to asses significance. See text for significant p values from spin permutation. | | | | |

**Table S20.** Results of linear models investigating effects of Gradient 2 and its interaction by hemisphere on the effects of linguistic parameters on brain activation in MEG data.

| | | Gradient 2 main effect | | | | Gradient 2 by hemisphere interaction | | | |
|---|---|---|---|---|---|---|---|---|---|
| **Parameter** | **Time [ms]** | **Est** | **SE** | **t** | **p** | **Est** | **SE** | **t** | **p** |
| **Word length** | 0-100 | -11.5 | 2.16 | -5.32 | 1.8e-07 | 2 | 3.08 | 0.65 | 0.52 |
| | 100-200 | -14.98 | 2.62 | -5.72 | 2.3e-08 | 6.42 | 3.74 | 1.72 | 0.087 |
| | 200-300 | -8.33 | 1.79 | -4.64 | 4.8e-06 | 7.72 | 2.56 | 3.02 | 0.0028 |
| | 300-400 | -6.85 | 1.93 | -3.55 | 0.00044 | 5.91 | 2.75 | 2.14 | 0.033 |
| | 400-500 | -0.39 | 1.69 | -0.23 | 0.82 | -9.93 | 2.41 | -4.12 | 4.8e-05 |
| **Orthographic distance** | 0-100 | 8.55 | 1.77 | 4.82 | 2.2e-06 | 1.02 | 2.53 | 0.4 | 0.69 |
| | 100-200 | 12.01 | 3.22 | 3.73 | 0.00022 | 7.1 | 4.6 | 1.55 | 0.12 |
| | 200-300 | 12.05 | 2.07 | 5.82 | 1.3e-08 | -13.84 | 2.95 | -4.69 | 4e-06 |
| | 300-400 | 5.43 | 2.03 | 2.67 | 0.008 | -3.58 | 2.9 | -1.23 | 0.22 |
| | 400-500 | -6.49 | 2.08 | -3.12 | 0.002 | 14.63 | 2.97 | 4.92 | 1.3e-06 |
| **Word frequency** | 0-100 | -4.73 | 1.83 | -2.58 | 0.01 | 10.69 | 2.62 | 4.08 | 5.5e-05 |
| | 100-200 | -7.67 | 2.13 | -3.6 | 0.00036 | 15.91 | 3.04 | 5.24 | 2.9e-07 |
| | 200-300 | -0.01 | 1.34 | -0.01 | 0.99 | -1.92 | 1.91 | -1.01 | 0.31 |
| | 300-400 | -6.63 | 1.29 | -5.12 | 5.1e-07 | 2.9 | 1.85 | 1.57 | 0.12 |
| | 400-500 | -3.87 | 1.65 | -2.34 | 0.02 | -2.54 | 2.36 | -1.08 | 0.28 |
| **Semantic similarity** | 0-100 | 2.27 | 1.42 | 1.59 | 0.11 | 1.48 | 2.03 | 0.73 | 0.47 |
| | 100-200 | 4.38 | 1.29 | 3.39 | 0.00079 | -4.99 | 1.85 | -2.7 | 0.0073 |
| | 200-300 | -0.89 | 1.04 | -0.85 | 0.39 | 3.84 | 1.48 | 2.59 | 0.0099 |
| | 300-400 | -4.39 | 1.06 | -4.15 | 4.1e-05 | 4.79 | 1.51 | 3.17 | 0.0016 |
| | 400-500 | -2.48 | 1.13 | -2.19 | 0.029 | -2.21 | 1.61 | -1.37 | 0.17 |
| **Position** | 0-100 | 1.93 | 1.37 | 1.4 | 0.16 | -0.18 | 1.96 | -0.09 | 0.93 |
| | 100-200 | 1.97 | 1.06 | 1.85 | 0.065 | 1.68 | 1.52 | 1.11 | 0.27 |
| | 200-300 | 4.27 | 1.1 | 3.89 | 0.00012 | 0.3 | 1.57 | 0.19 | 0.85 |
| | 300-400 | 0.27 | 1.21 | 0.22 | 0.82 | -3.9 | 1.72 | -2.27 | 0.024 |

| | | | | | | | | | |
|---|---|---|---|---|---|---|---|---|---|
| | 400-500 | 0.36 | 1.03 | 0.35 | 0.73 | -1.27 | 1.47 | -0.86 | 0.39 |
| Est = estimate, SE = standard error; in e-28 T. For p < .05, spin permutation was used to asses significance. See text for significant p values from spin permutation. | | | | | | | | | |

**Table S21.** Results of linear models investigating effects of Gradient 2 and its interaction by hemisphere on the effects of linguistic parameters on brain activation in MEG data when controlling for stimulus presentation duration instead of word length

| | | **Gradient 2 main effect** | | | | **Gradient 2 by hemisphere interaction** | | | |
|---|---|---|---|---|---|---|---|---|---|
| **Parameter** | **Time [ms]** | **Est** | **SE** | **t** | **p** | **Est** | **SE** | **t** | **p** |
| **Orthographic distance** | 0-100 | 3.84 | 1.45 | 2.64 | 0.0086 | 4.24 | 2.08 | 2.04 | 0.042 |
| | 100-200 | 1.66 | 2.79 | 0.6 | 0.55 | 14.24 | 3.98 | 3.58 | 0.00039 |
| | 200-300 | 5.39 | 1.48 | 3.65 | 3e-04 | -7.86 | 2.11 | -3.73 | 0.00023 |
| | 300-400 | 3.66 | 1.95 | 1.88 | 0.061 | 1.69 | 2.78 | 0.61 | 0.54 |
| | 400-500 | -10.35 | 2.23 | -4.65 | 4.8e-06 | 15.44 | 3.18 | 4.86 | 1.8e-06 |
| **Word frequency** | 0-100 | -3.83 | 1.92 | -1.99 | 0.047 | 9.56 | 2.74 | 3.49 | 0.00055 |
| | 100-200 | -4.77 | 2.13 | -2.24 | 0.026 | 13.57 | 3.04 | 4.46 | 1.1e-05 |
| | 200-300 | 1.99 | 1.45 | 1.37 | 0.17 | -3.7 | 2.07 | -1.78 | 0.075 |
| | 300-400 | -6.5 | 1.35 | -4.82 | 2.1e-06 | 1.25 | 1.93 | 0.65 | 0.52 |
| | 400-500 | -2.31 | 1.55 | -1.5 | 0.14 | -3.75 | 2.21 | -1.7 | 0.091 |
| **Semantic similarity** | 0-100 | 2.44 | 1.43 | 1.7 | 0.09 | 1.34 | 2.05 | 0.65 | 0.51 |
| | 100-200 | 4.79 | 1.31 | 3.66 | 0.00029 | -5.28 | 1.87 | -2.83 | 0.0049 |
| | 200-300 | -0.63 | 1.04 | -0.6 | 0.55 | 3.61 | 1.48 | 2.43 | 0.015 |
| | 300-400 | -4.35 | 1.07 | -4.08 | 5.6e-05 | 4.59 | 1.52 | 3.02 | 0.0028 |
| | 400-500 | -2.3 | 1.13 | -2.04 | 0.042 | -2.3 | 1.61 | -1.43 | 0.15 |
| **Position** | 0-100 | 2.47 | 1.4 | 1.76 | 0.079 | 0.22 | 2 | 0.11 | 0.91 |
| | 100-200 | 1.82 | 1.09 | 1.66 | 0.097 | 2.28 | 1.56 | 1.46 | 0.15 |
| | 200-300 | 4.01 | 1.11 | 3.61 | 0.00035 | 0.5 | 1.58 | 0.32 | 0.75 |
| | 300-400 | 0.77 | 1.2 | 0.64 | 0.52 | -3.62 | 1.72 | -2.11 | 0.036 |
| | 400-500 | -0.32 | 1.03 | -0.31 | 0.76 | 0.06 | 1.47 | 0.04 | 0.96 |

| Est = estimate, SE = standard error; in e-28 T. For p < .05, spin permutation was used to asses significance. See text for significant p values from spin permutation. |
|---|

**Table S22.** Results of linear models investigating effects of Gradient 3 and its interaction by hemisphere on the effects of linguistic parameters on brain activation in MEG data

| | | **Gradient 3 main effect** | | | | **Gradient 3 by hemisphere interaction** | | | |
|---|---|---|---|---|---|---|---|---|---|
| **Parameter** | **Time [ms]** | **Est** | **SE** | **t** | **p** | **Est** | **SE** | **t** | **p** |
| **Word length** | 0-100 | -7.48 | 3.21 | -2.33 | 0.02 | 2 | 3.08 | 0.65 | 0.52 |
| | 100-200 | -15.31 | 3.82 | -4 | 7.6e-05 | 6.42 | 3.74 | 1.72 | 0.087 |
| | 200-300 | -1.5 | 2.62 | -0.57 | 0.57 | 7.72 | 2.56 | 3.02 | 0.0028 |
| | 300-400 | 2.61 | 2.78 | 0.94 | 0.35 | 5.91 | 2.75 | 2.14 | 0.033 |
| | 400-500 | 7.12 | 2.47 | 2.89 | 0.0041 | -9.93 | 2.41 | -4.12 | 4.8e-05 |
| **Orthographic distance** | 0-100 | 3.94 | 2.66 | 1.48 | 0.14 | 1.02 | 2.53 | 0.4 | 0.69 |
| | 100-200 | 7.63 | 4.8 | 1.59 | 0.11 | 7.1 | 4.6 | 1.55 | 0.12 |
| | 200-300 | 4.25 | 3.06 | 1.39 | 0.17 | -13.84 | 2.95 | -4.69 | 4e-06 |
| | 300-400 | 2.28 | 2.92 | 0.78 | 0.43 | -3.58 | 2.9 | -1.23 | 0.22 |
| | 400-500 | -5.15 | 2.97 | -1.73 | 0.084 | 14.63 | 2.97 | 4.92 | 1.3e-06 |
| **Word frequency** | 0-100 | -1.62 | 2.66 | -0.61 | 0.54 | 10.69 | 2.62 | 4.08 | 5.5e-05 |
| | 100-200 | -11.53 | 3.06 | -3.77 | 0.00019 | 15.91 | 3.04 | 5.24 | 2.9e-07 |
| | 200-300 | -1.12 | 1.87 | -0.6 | 0.55 | -1.92 | 1.91 | -1.01 | 0.31 |
| | 300-400 | 2.05 | 1.92 | 1.07 | 0.29 | 2.9 | 1.85 | 1.57 | 0.12 |
| | 400-500 | 5.17 | 2.39 | 2.16 | 0.031 | -2.54 | 2.36 | -1.08 | 0.28 |
| **Semantic similarity** | 0-100 | 2.81 | 2.03 | 1.38 | 0.17 | 1.48 | 2.03 | 0.73 | 0.47 |
| | 100-200 | 3.49 | 1.85 | 1.88 | 0.061 | -4.99 | 1.85 | -2.7 | 0.0073 |
| | 200-300 | -2.47 | 1.48 | -1.67 | 0.096 | 3.84 | 1.48 | 2.59 | 0.0099 |
| | 300-400 | -3.97 | 1.52 | -2.61 | 0.0094 | 4.79 | 1.51 | 3.17 | 0.0016 |
| | 400-500 | 0.48 | 1.64 | 0.29 | 0.77 | -2.21 | 1.61 | -1.37 | 0.17 |
| **Position** | 0-100 | -2.59 | 1.95 | -1.33 | 0.19 | -0.18 | 1.96 | -0.09 | 0.93 |
| | 100-200 | -0.79 | 1.54 | -0.51 | 0.61 | 1.68 | 1.52 | 1.11 | 0.27 |

| | | | | | | | | | |
|---|---|---|---|---|---|---|---|---|---|
| | 200-300 | 0.4 | 1.62 | 0.25 | 0.8 | 0.3 | 1.57 | 0.19 | 0.85 |
| | 300-400 | -2.66 | 1.73 | -1.54 | 0.12 | -3.9 | 1.72 | -2.27 | 0.024 |
| | 400-500 | -0.56 | 1.45 | -0.39 | 0.7 | -1.27 | 1.47 | -0.86 | 0.39 |
| Est = estimate, SE = standard error; in e-28 T. For p < .05, spin permutation was used to asses significance. See text for significant p values from spin permutation. | | | | | | | | | |

**Table S23.** Results of linear models investigating effects of Gradient 3 and its interaction by hemisphere on the effects of linguistic parameters on brain activation in MEG data when controlling for stimulus presentation duration instead of word length

| | | **Gradient 3 main effect** | | | | **Gradient 3 by hemisphere interaction** | | | |
|---|---|---|---|---|---|---|---|---|---|
| **Parameter** | **Time [ms]** | **Est** | **SE** | **t** | **p** | **Est** | **SE** | **t** | **p** |
| **Orthographic distance** | 0-100 | 3.84 | 1.45 | 2.64 | 0.0086 | -1.28 | 4.58 | -0.28 | 0.78 |
| | 100-200 | 1.66 | 2.79 | 0.6 | 0.55 | 6.52 | 5.46 | 1.2 | 0.23 |
| | 200-300 | 5.39 | 1.48 | 3.65 | 3e-04 | -0.92 | 3.73 | -0.25 | 0.8 |
| | 300-400 | 3.66 | 1.95 | 1.88 | 0.061 | -3.36 | 3.97 | -0.85 | 0.4 |
| | 400-500 | -10.35 | 2.23 | -4.65 | 4.8e-06 | -13.29 | 3.52 | -3.78 | 0.00019 |
| **Word frequency** | 0-100 | -3.83 | 1.92 | -1.99 | 0.047 | 3.16 | 3.8 | 0.83 | 0.41 |
| | 100-200 | -4.77 | 2.13 | -2.24 | 0.026 | 5.76 | 6.86 | 0.84 | 0.4 |
| | 200-300 | 1.99 | 1.45 | 1.37 | 0.17 | -8.47 | 4.37 | -1.94 | 0.053 |
| | 300-400 | -6.5 | 1.35 | -4.82 | 2.1e-06 | -2.34 | 4.16 | -0.56 | 0.57 |
| | 400-500 | -2.31 | 1.55 | -1.5 | 0.14 | 17.6 | 4.24 | 4.15 | 4.2e-05 |
| **Semantic similarity** | 0-100 | 2.44 | 1.43 | 1.7 | 0.09 | 4.94 | 3.79 | 1.3 | 0.19 |
| | 100-200 | 4.79 | 1.31 | 3.66 | 0.00029 | 17.65 | 4.37 | 4.04 | 6.5e-05 |
| | 200-300 | -0.63 | 1.04 | -0.6 | 0.55 | -4.82 | 2.67 | -1.8 | 0.072 |
| | 300-400 | -4.35 | 1.07 | -4.08 | 5.6e-05 | -4.63 | 2.74 | -1.69 | 0.091 |
| | 400-500 | -2.3 | 1.13 | -2.04 | 0.042 | -7.53 | 3.41 | -2.21 | 0.028 |
| **Position** | 0-100 | 2.47 | 1.4 | 1.76 | 0.079 | 0.42 | 2.9 | 0.15 | 0.88 |
| | 100-200 | 1.82 | 1.09 | 1.66 | 0.097 | -1.42 | 2.65 | -0.54 | 0.59 |
| | 200-300 | 4.01 | 1.11 | 3.61 | 0.00035 | 4.7 | 2.11 | 2.23 | 0.027 |

| | | | | | | | | |
|---|---|---|---|---|---|---|---|---|
| | 300-400 | 0.77 | 1.2 | 0.64 | 0.52 | 5.76 | 2.17 | 2.66 | 0.0083 |
| | 400-500 | -0.32 | 1.03 | -0.31 | 0.76 | -3.46 | 2.34 | -1.48 | 0.14 |
| Est = estimate, SE = standard error; in e-28 T. For p < .05, spin permutation was used to asses significance. See text for significant p values from spin permutation. | | | | | | | | | |

**Table S24.** Results of the ANOVA investigating differences in linguistic parameters' associations with Gradient 2 (interaction between parameters' effects on brain activation with Gradient 2 and hemisphere) and follow-up linear models for each hemisphere investigating pairwise parameter differences (interaction between parameters' effects on brain activation with Gradient 2) in MEG data

| **ANOVA results** | | | | | |
|---|---|---|---|---|---|
| **Parameter** | **Time [ms]** | **Sum of Sqares** | **Mean Squares** | **F** | **p** |
| All (Interaction with gradient) | 0-100 | 27.59 | 3.07 | 22.72 | 5.7e-38 |
| | 100-200 | 29.14 | 3.24 | 26.13 | 4.8e-44 |
| | 200-300 | 5.07 | 0.56 | 5.91 | 3.2e-08 |
| | 300-400 | 6.75 | 0.75 | 8.77 | 3.8e-13 |
| | 400-500 | 4.62 | 0.51 | 5.95 | 2.7e-08 |
| All (Interaction with gradient and hemisphere) | 0-100 | 2.29 | 0.25 | 1.89 | 0.049 |
| | 100-200 | 3.65 | 0.41 | 3.27 | 0.00057 |
| | 200-300 | 1.64 | 0.18 | 1.91 | 0.046 |
| | 300-400 | 0.86 | 0.1 | 1.11 | 0.35 |
| | 400-500 | 3.76 | 0.42 | 4.84 | 1.8e-06 |
| **Follow-up linear model results left hemisphere** | | | | | |
| **Parameter** | **Time [ms]** | **Estimate** | **SE** | **t** | **p** |
| Word length vs. Orthographic distance | 100-200 | -2.97 | 4.4 | -0.68 | 0.5 |
| Word length vs. Word frequency | | 22.66 | 3.74 | 6.05 | 3.7e-09 |
| Word length vs. Semantic similarity | | 10.6 | 3.26 | 3.25 | 0.0013 |
| Word length vs. Position | | -13.02 | 3.21 | -4.06 | 6.1e-05 |
| Orthographic distance vs. Word frequency | | 19.69 | 3.9 | 5.05 | 7.2e-07 |
| Orthographic distance vs. Semantic similarity | | 7.63 | 3.44 | 2.22 | 0.027 |

| | | | | | |
|---|---|---|---|---|---|
| Orthographic distance vs. Position | | -10.05 | 3.39 | -2.97 | 0.0032 |
| Word frequency vs. Semantic similarity | | -12.06 | 2.55 | -4.73 | 3.3e-06 |
| Word frequency vs. Position | | 9.64 | 2.48 | 3.89 | 0.00012 |
| Semantic similarity vs. Position | | -2.42 | 1.67 | -1.45 | 0.15 |
| Word length vs. Orthographic distance | 400-500 | -6.1 | 2.26 | -2.69 | 0.0074 |
| Word length vs. Word frequency | | 3.48 | 2.29 | 1.52 | 0.13 |
| Word length vs. Semantic similarity | | -2.87 | 1.89 | -1.52 | 0.13 |
| Word length vs. Position | | 0.75 | 1.85 | 0.4 | 0.69 |
| Orthographic distance vs. Word frequency | | -2.62 | 2.47 | -1.06 | 0.29 |
| Orthographic distance vs. Semantic similarity | | -8.97 | 2.1 | -4.26 | 2.6e-05 |
| Orthographic distance vs. Position | | 6.85 | 2.07 | 3.31 | 0.001 |
| Word frequency vs. Semantic similarity | | -6.35 | 2.13 | -2.98 | 0.0031 |
| Word frequency vs. Position | | 4.23 | 2.1 | 2.02 | 0.045 |
| Semantic similarity vs. Position | | -2.12 | 1.65 | -1.28 | 0.2 |
| **Follow-up linear model results right hemisphere** | | | | | |
| **Parameter** | **Time [ms]** | **Estimate** | **SE** | **t** | **p** |
| Word length vs. | 100-200 | 10.55 | 3.95 | 2.67 | 0.008 |

| | | | | | |
|---|---|---|---|---|---|
| Orthographic distance | | | | | |
| Word length vs. Word frequency | | 0.33 | 3.01 | 0.11 | 0.91 |
| Word length vs. Semantic similarity | | 9.17 | 2.57 | 3.57 | 0.00041 |
| Word length vs. Position | | -4.92 | 2.42 | -2.04 | 0.043 |
| Orthographic distance vs. Word frequency | | 10.88 | 3.89 | 2.8 | 0.0055 |
| Orthographic distance vs. Semantic similarity | | 19.72 | 3.57 | 5.53 | 6.4e-08 |
| Orthographic distance vs. Position | | -15.47 | 3.46 | -4.48 | 1e-05 |
| Word frequency vs. Semantic similarity | | 8.85 | 2.48 | 3.57 | 4e-04 |
| Word frequency vs. Position | | -4.6 | 2.31 | -1.99 | 0.048 |
| Semantic similarity vs. Position | | 4.25 | 1.71 | 2.48 | 0.014 |
| Word length vs. Orthographic distance | 400-500 | 18.46 | 3.11 | 5.94 | 6.9e-09 |
| Word length vs. Word frequency | | -3.91 | 2.48 | -1.58 | 0.12 |
| Word length vs. Semantic similarity | | -15 | 2.2 | -6.81 | 4.5e-11 |
| Word length vs. Position | | 9.41 | 2.14 | 4.4 | 1.5e-05 |
| Orthographic distance vs. Word frequency | | 14.55 | 2.89 | 5.03 | 8.1e-07 |
| Orthographic distance vs. Semantic similarity | | 3.46 | 2.66 | 1.3 | 0.19 |

| | | | | | |
|---|---|---|---|---|---|
| Orthographic distance vs. Position | | -9.05 | 2.61 | -3.47 | 0.00059 |
| Word frequency vs. Semantic similarity | | -11.09 | 1.89 | -5.86 | 1.1e-08 |
| Word frequency vs. Position | | 5.5 | 1.82 | 3.02 | 0.0027 |
| Semantic similarity vs. Position | | -5.59 | 1.42 | -3.94 | 9.8e-05 |
| SE = standard error. Sum of squares and mean squares in e-54 T. Estimate and SE in e-28 T. For p < .05, spin permutation was used to asses significance. See text for significant p values from spin permutation. | | | | | |

**Table S25.** Results of the ANOVA investigating differences in linguistic parameters' associations with Gradient 2 (interaction between parameters' effects on brain activation with Gradient 2 and hemisphere) and follow-up linear models for each hemisphere investigating pairwise parameter differences (interaction between parameters' effects on brain activation with Gradient 2) in MEG data when controlling for stimulus presentation duration instead of word length

| **ANOVA results** | | | | | |
|---|---|---|---|---|---|
| **Parameter** | **Time [ms]** | **Sum of Sqares** | **Mean Squares** | **F** | **p** |
| All (Interaction with gradient) | 0-100 | 24.87 | 2.76 | 20.49 | 5.6e-34 |
| | 100-200 | 21.16 | 2.35 | 19.45 | 3.9e-32 |
| | 200-300 | 3.87 | 0.43 | 4.64 | 4e-06 |
| | 300-400 | 7.43 | 0.83 | 9.83 | 5.3e-15 |
| | 400-500 | 3.96 | 0.44 | 5.04 | 8.9e-07 |
| All (Interaction with gradient and hemisphere) | 0-100 | 1.92 | 0.21 | 1.58 | 0.11 |
| | 100-200 | 3.43 | 0.38 | 3.15 | 0.00085 |
| | 200-300 | 1.07 | 0.12 | 1.28 | 0.24 |
| | 300-400 | 0.54 | 0.06 | 0.72 | 0.7 |
| | 400-500 | 4.22 | 0.47 | 5.38 | 2.5e-07 |
| **Follow-up linear model results left hemisphere** | | | | | |
| **Parameter** | **Time [ms]** | **Estimate** | **SE** | **t** | **p** |
| Orthographic distance vs. Word frequency | 100-200 | 6.43 | 3.55 | 1.81 | 0.071 |
| Orthographic distance vs. Semantic similarity | | -3.13 | 3.11 | -1.01 | 0.31 |

| | | | | | |
|---|---|---|---|---|---|
| Orthographic distance vs. Position | | 0.16 | 3.07 | 0.05 | 0.96 |
| Word frequency vs. Semantic similarity | | -9.56 | 2.48 | -3.85 | 0.00014 |
| Word frequency vs. Position | | 6.58 | 2.44 | 2.7 | 0.0072 |
| Semantic similarity vs. Position | | -2.97 | 1.72 | -1.72 | 0.085 |
| Orthographic distance vs. Word frequency | 400-500 | -8.04 | 2.73 | -2.95 | 0.0034 |
| Orthographic distance vs. Semantic similarity | | -12.65 | 2.48 | -5.1 | 5.6e-07 |
| Orthographic distance vs. Position | | 10.03 | 2.44 | 4.11 | 4.9e-05 |
| Word frequency vs. Semantic similarity | | -4.61 | 2.04 | -2.26 | 0.024 |
| Word frequency vs. Position | | 2 | 1.99 | 1 | 0.32 |
| Semantic similarity vs. Position | | -2.62 | 1.64 | -1.6 | 0.11 |
| **Follow-up linear model results right hemisphere** | | | | | |
| **Parameter** | **Time [ms]** | **Estimate** | **SE** | **t** | **p** |
| Orthographic distance vs. Word frequency | 100-200 | 7.1 | 3.53 | 2.01 | 0.045 |
| Orthographic distance vs. Semantic similarity | | 16.4 | 3.11 | 5.28 | 2.4e-07 |
| Orthographic distance vs. Position | | -11.81 | 2.97 | -3.98 | 8.5e-05 |
| Word frequency vs. Semantic similarity | | 9.29 | 2.56 | 3.63 | 0.00033 |

| | | | | | |
|---|---|---|---|---|---|
| Word frequency vs. Position | | -4.71 | 2.39 | -1.97 | 0.05 |
| Semantic similarity vs. Position | | 4.59 | 1.72 | 2.67 | 0.0078 |
| Orthographic distance vs. Word frequency | 400-500 | 11.16 | 2.75 | 4.06 | 6.1e-05 |
| Orthographic distance vs. Semantic similarity | | 0.49 | 2.56 | 0.19 | 0.85 |
| Orthographic distance vs. Position | | -5.34 | 2.51 | -2.13 | 0.034 |
| Word frequency vs. Semantic similarity | | -10.66 | 1.81 | -5.89 | 9.4e-09 |
| Word frequency vs. Position | | 5.81 | 1.74 | 3.33 | 0.00096 |
| Semantic similarity vs. Position | | -4.85 | 1.43 | -3.4 | 0.00075 |
| SE = standard error. Sum of squares and mean squares in e-54 T. Estimate and SE in e-28 T. For p < .05, spin permutation was used to asses significance. See text for significant p values from spin permutation. | | | | | |

**Table S26.** Results of the ANOVA investigating differences in each linguistic parameter's association with Gradient 2 across time (interaction between time, Gradient 2 and hemisphere) and follow-up linear models investigating pairwise time window differences for word frequency effects (interaction between time and Gradient 2) in MEG data

| **ANOVA results** | | | | | |
|---|---|---|---|---|---|
| **Parameter** | **Effect** | **Sum of Squares** | **Mean Squares** | **F** | **p** |
| Word length | Principal gradient x Time | 5.17 | 1.29 | 32.48 | 3.8e-26 |
| | Principal gradient x Time x Hemisphere | 1.09 | 0.27 | 6.85 | 1.8e-05 |
| Orthographic distance | Principal gradient x Time | 6.32 | 1.58 | 32.23 | 6e-26 |

| | | | | | |
|---|---|---|---|---|---|
| | Principal gradient x Time x Hemisphere | 2.05 | 0.51 | 10.46 | 2.2e-08 |
| Word frequency | Principal gradient x Time | 1.01 | 0.25 | 9.58 | 1.2e-07 |
| | Principal gradient x Time x Hemisphere | 1.43 | 0.36 | 13.58 | 6.6e-11 |
| Semantic similarity | Principal gradient x Time | 0.59 | 0.15 | 11.02 | 8e-09 |
| | Principal gradient x Time x Hemisphere | 0.31 | 0.08 | 5.77 | 0.00013 |
| Position | Principal gradient x Time | 0.4 | 0.1 | 7.99 | 2.2e-06 |
| | Principal gradient x Time x Hemisphere | 0.08 | 0.02 | 1.6 | 0.17 |
| **Follow-up linear model results** | | | | | |
| **Parameter** | **Time** | **Estimate** | **SE** | **t** | **p** |
| Word length | 0-100 vs. 100-200 | 22.23 | 2.43 | 9.14 | 6.9e-19 |
| | 0-100 vs. 200-300 | 5.79 | 2.03 | 2.85 | 0.0046 |
| | 0-100 vs. 300-400 | 6.56 | 2.09 | 3.14 | 0.0018 |
| | 0-100 vs. 400-500 | 5.12 | 1.99 | 2.58 | 0.01 |
| | 100-200 vs. 200-300 | -16.45 | 2.29 | -7.2 | 1.6e-12 |
| | 100-200 vs. 300-400 | -15.68 | 2.34 | -6.71 | 4e-11 |
| | 100-200 vs. 400-500 | -17.12 | 2.24 | -7.64 | 7.5e-14 |
| | 200-300 vs. 300-400 | 0.77 | 1.92 | 0.4 | 0.69 |
| | 200-300 vs. 400-500 | -0.67 | 1.8 | -0.37 | 0.71 |
| | 300-400 vs. 400-500 | -1.44 | 1.86 | -0.77 | 0.44 |
| **Follow-up linear model results left hemisphere** | | | | | |
| **Parameter** | **Time** | **Estimate** | **SE** | **t** | **p** |
| Orthographic distance | 0-100 vs. 100-200 | 20.56 | 3.7 | 5.56 | 5.3e-08 |

| | | | | | |
|---|---|---|---|---|---|
| | 0-100 vs. 200-300 | 20.59 | 2.63 | 7.83 | 6e-14 |
| | 0-100 vs. 300-400 | 3.12 | 2.54 | 1.23 | 0.22 |
| | 0-100 vs. 400-500 | 2.06 | 2.53 | 0.81 | 0.42 |
| | 100-200 vs. 200-300 | 0.03 | 3.71 | 0.01 | 0.99 |
| | 100-200 vs. 300-400 | -17.45 | 3.65 | -4.78 | 2.6e-06 |
| | 100-200 vs. 400-500 | -18.51 | 3.64 | -5.08 | 6.1e-07 |
| | 200-300 vs. 300-400 | -17.48 | 2.57 | -6.81 | 4.3e-11 |
| | 200-300 vs. 400-500 | -18.54 | 2.55 | -7.27 | 2.4e-12 |
| | 300-400 vs. 400-500 | -1.06 | 2.46 | -0.43 | 0.67 |
| **Follow-up linear model results right hemisphere** | | | | | |
| **Parameter** | **Time** | **Estimate** | **SE** | **t** | **p** |
| Orthographic distance | 0-100 vs. 100-200 | 28.69 | 3.72 | 7.7 | 1.5e-13 |
| | 0-100 vs. 200-300 | 7.77 | 2.87 | 2.71 | 0.0071 |
| | 0-100 vs. 300-400 | 7.71 | 2.91 | 2.65 | 0.0083 |
| | 0-100 vs. 400-500 | 17.71 | 2.99 | 5.92 | 7.8e-09 |
| | 100-200 vs. 200-300 | -20.91 | 4.01 | -5.21 | 3.3e-07 |
| | 100-200 vs. 300-400 | -20.97 | 4.04 | -5.19 | 3.6e-07 |
| | 100-200 vs. 400-500 | -10.98 | 4.1 | -2.68 | 0.0078 |
| | 200-300 vs. 300-400 | -0.06 | 3.27 | -0.02 | 0.99 |
| | 200-300 vs. 400-500 | 9.94 | 3.34 | 2.97 | 0.0032 |
| | 300-400 vs. 400-500 | 10 | 3.37 | 2.96 | 0.0033 |
| SE = standard error. Sum of squares and mean squares in e-54 T. Estimate and SE in e-28 T. For F or p < .05, spin permutation was used to asses significance. See text for significant p values from spin permutation. | | | | | |

**Table S27.** Results of the ANOVA investigating differences in each linguistic parameter's association with Gradient 2 across time (interaction between time, Gradient 2 and hemisphere) and follow-up linear models investigating pairwise time window differences for word frequency effects (interaction between time and Gradient 2) in MEG data when controlling for stimulus presentation duration instead of word length

| ANOVA results | | | | | |
|---|---|---|---|---|---|
| **Parameter** | **Effect** | **Sum of Squares** | **Mean Squares** | **F** | **p** |
| Orthographic distance | Principal gradient x Time | 2.5 | 0.63 | 16.09 | 6.1e-13 |
| | Principal gradient x Time x Hemisphere | 2.17 | 0.54 | 13.95 | 3.3e-11 |
| Word frequency | Principal gradient x Time | 0.96 | 0.24 | 8.84 | 4.6e-07 |
| | Principal gradient x Time x Hemisphere | 1.2 | 0.3 | 11.09 | 7e-09 |
| Semantic similarity | Principal gradient x Time | 0.61 | 0.15 | 11.35 | 4.3e-09 |
| | Principal gradient x Time x Hemisphere | 0.31 | 0.08 | 5.66 | 0.00016 |
| Position | Principal gradient x Time | 0.34 | 0.09 | 6.68 | 2.5e-05 |
| | Principal gradient x Time x Hemisphere | 0.08 | 0.02 | 1.65 | 0.16 |
| **Follow-up linear model results left hemisphere** | | | | | |
| **Parameter** | **Time** | **Estimate** | **SE** | **t** | **p** |
| Orthographic distance | 0-100 vs. 100-200 | -2.18 | 3.2 | -0.68 | 0.5 |
| | 0-100 vs. 200-300 | 1.55 | 2.06 | 0.75 | 0.45 |
| | 0-100 vs. 300-400 | -0.19 | 2.26 | -0.08 | 0.93 |
| | 0-100 vs. 400-500 | -14.19 | 2.63 | -5.4 | 1.2e-07 |
| | 100-200 vs. 200-300 | 3.73 | 3.18 | 1.17 | 0.24 |

| | | | | | |
|---|---|---|---|---|---|
| | 100-200 vs. 300-400 | 2 | 3.31 | 0.6 | 0.55 |
| | 100-200 vs. 400-500 | -12.01 | 3.57 | -3.36 | 0.00086 |
| | 200-300 vs. 300-400 | -1.74 | 2.23 | -0.78 | 0.44 |
| | 200-300 vs. 400-500 | -15.74 | 2.6 | -6.05 | 3.7e-09 |
| | 300-400 vs. 400-500 | -14.01 | 2.76 | -5.07 | 6.3e-07 |
| **Follow-up linear model results right hemisphere** | | | | | |
| **Parameter** | **Time** | **Estimate** | **SE** | **t** | **p** |
| Orthographic distance | 0-100 vs. 100-200 | 7.82 | 3.14 | 2.49 | 0.013 |
| | 0-100 vs. 200-300 | -10.55 | 2.12 | -4.97 | 1e-06 |
| | 0-100 vs. 300-400 | -2.73 | 2.64 | -1.03 | 0.3 |
| | 0-100 vs. 400-500 | -2.99 | 2.74 | -1.09 | 0.28 |
| | 100-200 vs. 200-300 | -18.37 | 3.19 | -5.76 | 1.9e-08 |
| | 100-200 vs. 300-400 | -10.55 | 3.56 | -2.97 | 0.0032 |
| | 100-200 vs. 400-500 | -10.81 | 3.63 | -2.98 | 0.0031 |
| | 200-300 vs. 300-400 | 7.81 | 2.7 | 2.9 | 0.004 |
| | 200-300 vs. 400-500 | 7.55 | 2.79 | 2.7 | 0.0072 |
| | 300-400 vs. 400-500 | -0.26 | 3.21 | -0.08 | 0.94 |
| SE = standard error. Sum of squares and mean squares in e-54 T. Estimate and SE in e-28 T. For F or p < .05, spin permutation was used to asses significance. See text for significant p values from spin permutation. | | | | | |

**Table S28.** Results of the ANOVA investigating differences in linguistic parameters' associations with Gradient 3 (interaction between parameters' effects on brain activation with Gradient 3 and hemisphere) and follow-up linear models for each hemisphere investigating pairwise parameter differences (interaction between parameters' effects on brain activation with Gradient 3) in MEG data

| ANOVA results | | | | | |
|---|---|---|---|---|---|
| **Parameter** | **Time [ms]** | **Sum of Sqares** | **Mean Squares** | **F** | **p** |
| All (Interaction with gradient) | 0-100 | 3.52 | 0.39 | 2.73 | 0.0036 |
| | 100-200 | 7.46 | 0.83 | 6.36 | 5.5e-09 |
| | 200-300 | 2.54 | 0.28 | 2.92 | 0.0019 |
| | 300-400 | 1.23 | 0.14 | 1.58 | 0.12 |
| | 400-500 | 1.63 | 0.18 | 2.06 | 0.03 |
| All (Interaction with gradient and hemisphere) | 0-100 | 0.52 | 0.06 | 0.4 | 0.93 |
| | 100-200 | 2.82 | 0.31 | 2.41 | 0.01 |
| | 200-300 | 0.27 | 0.03 | 0.32 | 0.97 |
| | 300-400 | 1.67 | 0.19 | 2.14 | 0.023 |
| | 400-500 | 1.98 | 0.22 | 2.5 | 0.0076 |
| **Follow-up linear model results left hemisphere** | | | | | |
| **Parameter** | **Time [ms]** | **Estimate** | **SE** | **t** | **p** |
| Word length vs. Orthographic distance | 100-200 | -7.68 | 6.44 | -1.19 | 0.23 |
| Word length vs. Word frequency | | 26.84 | 5.42 | 4.95 | 1.1e-06 |
| Word length vs. Semantic similarity | | 11.82 | 4.78 | 2.48 | 0.014 |
| Word length vs. Position | | -16.1 | 4.69 | -3.43 | 0.00067 |
| Orthographic distance vs. Word frequency | | 19.16 | 5.64 | 3.4 | 0.00076 |
| Orthographic distance vs. Semantic similarity | | 4.15 | 5.03 | 0.82 | 0.41 |
| Orthographic distance vs. Position | | -8.42 | 4.95 | -1.7 | 0.09 |
| Word frequency vs. Semantic similarity | | -15.02 | 3.63 | -4.14 | 4.4e-05 |

| | | | | | |
|---|---|---|---|---|---|
| Word frequency vs. Position | | 10.74 | 3.51 | 3.06 | 0.0024 |
| Semantic similarity vs. Position | | -4.27 | 2.41 | -1.78 | 0.077 |
| **Follow-up linear model results right hemisphere** | | | | | |
| **Parameter** | **Time [ms]** | **Estimate** | **SE** | **t** | **p** |
| Word length vs. Orthographic distance | 100-200 | 4.6 | 5.92 | 0.78 | 0.44 |
| Word length vs. Word frequency | 100-200 | 2.67 | 4.38 | 0.61 | 0.54 |
| Word length vs. Semantic similarity | 100-200 | 6.72 | 3.7 | 1.82 | 0.07 |
| Word length vs. Position | 100-200 | -9.13 | 3.51 | -2.6 | 0.0097 |
| Orthographic distance vs. Word frequency | 100-200 | 7.27 | 5.85 | 1.24 | 0.22 |
| Orthographic distance vs. Semantic similarity | 100-200 | 11.32 | 5.36 | 2.11 | 0.035 |
| Orthographic distance vs. Position | 100-200 | -13.73 | 5.23 | -2.62 | 0.0091 |
| Word frequency vs. Semantic similarity | 100-200 | 4.06 | 3.59 | 1.13 | 0.26 |
| Word frequency vs. Position | 100-200 | -6.46 | 3.39 | -1.91 | 0.058 |
| Semantic similarity vs. Position | 100-200 | -2.4 | 2.45 | -0.98 | 0.33 |
| SE = standard error. Sum of squares and mean squares in e-54 T. Estimate and SE in e-28 T. For F or p < .05, spin permutation was used to asses significance. See text for significant p values from spin permutation. | | | | | |

**Table S29.** Results of the ANOVA investigating differences in linguistic parameters' associations with Gradient 3 (interaction between parameters' effects on brain activation with Gradient 3 and hemisphere) and follow-up linear models for each hemisphere investigating pairwise parameter differences (interaction between parameters' effects on brain activation with Gradient 3) in MEG data when controlling for stimulus presentation duration instead of word length

| **ANOVA results** | | | | | |
|---|---|---|---|---|---|
| **Parameter** | **Time [ms]** | **Sum of Sqares** | **Mean Squares** | **F** | **p** |
| All (Interaction with gradient) | 0-100 | 2.63 | 0.29 | 2.05 | 0.031 |
| | 100-200 | 5.96 | 0.66 | 5.27 | 3.6e-07 |
| | 200-300 | 2.31 | 0.26 | 2.74 | 0.0034 |
| | 300-400 | 1.44 | 0.16 | 1.87 | 0.052 |
| | 400-500 | 1.6 | 0.18 | 2 | 0.035 |
| All (Interaction with gradient and hemisphere) | 0-100 | 0.54 | 0.06 | 0.42 | 0.92 |
| | 100-200 | 2.75 | 0.31 | 2.43 | 0.0093 |
| | 200-300 | 0.26 | 0.03 | 0.31 | 0.97 |
| | 300-400 | 1.51 | 0.17 | 1.96 | 0.04 |
| | 400-500 | 1.82 | 0.2 | 2.28 | 0.015 |
| **Follow-up linear model results left hemisphere** | | | | | |
| **Parameter** | **Time [ms]** | **Estimate** | **SE** | **t** | **p** |
| Orthographic distance vs. Word frequency | 100-200 | 14.7 | 4.99 | 2.95 | 0.0034 |
| Orthographic distance vs. Semantic similarity | | -0.1 | 4.42 | -0.02 | 0.98 |
| Orthographic distance vs. Position | | -3.18 | 4.36 | -0.73 | 0.47 |
| Word frequency vs. Semantic similarity | | -14.8 | 3.48 | -4.25 | 2.7e-05 |
| Word frequency vs. Position | | 11.52 | 3.4 | 3.39 | 0.00077 |
| Semantic similarity vs. Position | | -3.28 | 2.49 | -1.32 | 0.19 |
| **Follow-up linear model results right hemisphere** | | | | | |
| **Parameter** | **Time [ms]** | **Estimate** | **SE** | **t** | **p** |
| Orthographic distance vs. Word frequency | 100-200 | 5.54 | 5.28 | 1.05 | 0.29 |

| | | | | | |
|---|---|---|---|---|---|
| Orthographic distance vs. Semantic similarity | | 9.51 | 4.64 | 2.05 | 0.041 |
| Orthographic distance vs. Position | | -11.19 | 4.48 | -2.5 | 0.013 |
| Word frequency vs. Semantic similarity | | 3.96 | 3.72 | 1.06 | 0.29 |
| Word frequency vs. Position | | -5.65 | 3.52 | -1.6 | 0.11 |
| Semantic similarity vs. Position | | -1.69 | 2.47 | -0.68 | 0.49 |
| SE = standard error. Sum of squares and mean squares in e-54 T. Estimate and SE in e-28 T. For p < .05, spin permutation was used to asses significance. See text for significant p values from spin permutation. | | | | | |

**Table S30.** Results of the ANOVA investigating differences in each linguistic parameter's association with Gradient 3 across time (interaction between time, Gradient 3 and hemisphere) and follow-up linear models investigating pairwise time window differences for word frequency effects (interaction between time and Gradient 3) in MEG data

| **ANOVA results** | | | | | |
|---|---|---|---|---|---|
| **Parameter** | **Effect** | **Sum of Squares** | **Mean Squares** | **F** | **p** |
| Word length | Principal gradient x Time | 2 | 0.5 | 11.85 | 1.7e-09 |
| | Principal gradient x Time x Hemisphere | 0.24 | 0.06 | 1.4 | 0.23 |
| Orthographic distance | Principal gradient x Time | 1.42 | 0.36 | 6.74 | 2.2e-05 |
| | Principal gradient x Time x Hemisphere | 0.89 | 0.22 | 4.21 | 0.0021 |
| Word frequency | Principal gradient x Time | 0.21 | 0.05 | 1.92 | 0.1 |
| | Principal gradient x Time x Hemisphere | 0.9 | 0.22 | 8.23 | 1.4e-06 |
| Semantic similarity | Principal gradient x Time | 0.14 | 0.04 | 2.56 | 0.037 |
| | Principal gradient x Time x Hemisphere | 0.15 | 0.04 | 2.83 | 0.024 |
| Position | Principal gradient x Time | 0.06 | 0.02 | 1.2 | 0.31 |
| | Principal gradient x Time x Hemisphere | 0.02 | 0.01 | 0.43 | 0.78 |
| **Follow-up linear model results** | | | | | |
| **Parameter** | **Time** | **Estimate** | **SE** | **t** | **p** |
| Word length | 0-100 vs. 100-200 | 20.56 | 3.56 | 5.77 | 1.2e-08 |
| | 0-100 vs. 200-300 | 6.83 | 2.97 | 2.3 | 0.022 |

| | | | | | |
|---|---|---|---|---|---|
| | 0-100 vs. 300-400 | 9.07 | 3.04 | 2.98 | 0.003 |
| | 0-100 vs. 400-500 | 9.18 | 2.91 | 3.15 | 0.0017 |
| | 100-200 vs. 200-300 | -13.73 | 3.31 | -4.14 | 3.8e-05 |
| | 100-200 vs. 300-400 | -11.48 | 3.38 | -3.4 | 0.00071 |
| | 100-200 vs. 400-500 | -11.38 | 3.26 | -3.49 | 0.00051 |
| | 200-300 vs. 300-400 | 2.24 | 2.75 | 0.82 | 0.41 |
| | 200-300 vs. 400-500 | 2.35 | 2.6 | 0.9 | 0.37 |
| | 300-400 vs. 400-500 | 0.1 | 2.69 | 0.04 | 0.97 |
| **Follow-up linear model results left hemisphere** | | | | | |
| **Parameter** | **Time** | **Estimate** | **SE** | **t** | **p** |
| Word frequency | 0-100 vs. 100-200 | -9.91 | 4.22 | -2.35 | 0.019 |
| | 0-100 vs. 200-300 | 0.5 | 3.52 | 0.14 | 0.89 |
| | 0-100 vs. 300-400 | -0.43 | 3.54 | -0.12 | 0.9 |
| | 0-100 vs. 400-500 | 6.78 | 3.77 | 1.8 | 0.073 |
| | 100-200 vs. 200-300 | 10.41 | 3.78 | 2.76 | 0.0062 |
| | 100-200 vs. 300-400 | 9.48 | 3.8 | 2.49 | 0.013 |
| | 100-200 vs. 400-500 | 16.7 | 4.02 | 4.16 | 4e-05 |
| | 200-300 vs. 300-400 | -0.93 | 3 | -0.31 | 0.76 |
| | 200-300 vs. 400-500 | 6.29 | 3.27 | 1.92 | 0.055 |
| | 300-400 vs. 400-500 | 7.22 | 3.3 | 2.19 | 0.029 |
| **Follow-up linear model results right hemisphere** | | | | | |
| **Parameter** | **Time** | **Estimate** | **SE** | **t** | **p** |
| Word frequency | 0-100 vs. 100-200 | 2.79 | 3.94 | 0.71 | 0.48 |
| | 0-100 vs. 200-300 | -9.26 | 3 | -3.08 | 0.0022 |
| | 0-100 vs. 300-400 | -0.75 | 3.03 | -0.25 | 0.81 |
| | 0-100 vs. 400-500 | -5.69 | 3.42 | -1.66 | 0.097 |
| | 100-200 vs. 200-300 | -12.06 | 3.44 | -3.51 | 0.00052 |

| | | | | | |
|---|---|---|---|---|---|
| | 100-200 vs. 300-400 | -3.54 | 3.47 | -1.02 | 0.31 |
| | 100-200 vs. 400-500 | -8.48 | 3.81 | -2.23 | 0.027 |
| | 200-300 vs. 300-400 | 8.52 | 2.34 | 3.63 | 0.00032 |
| | 200-300 vs. 400-500 | 3.57 | 2.83 | 1.26 | 0.21 |
| | 300-400 vs. 400-500 | -4.94 | 2.86 | -1.73 | 0.085 |
| SE = standard error. Sum of squares and mean squares in e-54 T. Estimate and SE in e-28 T. For F or p < .05, spin permutation was used to asses significance. See text for significant p values from spin permutation. | | | | | |

**Table S31.** Results of the ANOVA investigating differences in each linguistic parameter's association with Gradient 3 across time (interaction between time, Gradient 3 and hemisphere) and follow-up linear models investigating pairwise time window differences for word frequency effects (interaction between time and Gradient 3) in MEG data when controlling for stimulus presentation duration instead of word length

| ANOVA results | | | | | |
|---|---|---|---|---|---|
| **Parameter** | **Effect** | **Sum of Squares** | **Mean Squares** | **F** | **p** |
| Orthographic distance | Principal gradient x Time | 0.89 | 0.22 | 5.48 | 0.00022 |
| | Principal gradient x Time x Hemisphere | 0.84 | 0.21 | 5.18 | 0.00038 |
| Word frequency | Principal gradient x Time | 0.26 | 0.07 | 2.34 | 0.053 |
| | Principal gradient x Time x Hemisphere | 0.9 | 0.22 | 8.03 | 2e-06 |
| Semantic similarity | Principal gradient x Time | 0.16 | 0.04 | 2.83 | 0.023 |
| | Principal gradient x Time x Hemisphere | 0.16 | 0.04 | 2.82 | 0.024 |
| Position | Principal gradient x Time | 0.04 | 0.01 | 0.78 | 0.54 |
| | Principal gradient x | 0.04 | 0.01 | 0.69 | 0.6 |

| | | | | | |
|---|---|---|---|---|---|
| | Time x Hemisphere | | | | |
| **Follow-up linear model results left hemisphere** | | | | | |
| **Parameter** | **Time** | **Estimate** | **SE** | **t** | **p** |
| Word frequency | 0-100 vs. 100-200 | -11.39 | 4.22 | -2.7 | 0.0073 |
| | 0-100 vs. 200-300 | -0.38 | 3.8 | -0.1 | 0.92 |
| | 0-100 vs. 300-400 | -0.07 | 3.74 | -0.02 | 0.98 |
| | 0-100 vs. 400-500 | 5.66 | 3.79 | 1.49 | 0.14 |
| | 100-200 vs. 200-300 | 11.01 | 3.76 | 2.93 | 0.0036 |
| | 100-200 vs. 300-400 | 11.32 | 3.69 | 3.07 | 0.0023 |
| | 100-200 vs. 400-500 | 17.05 | 3.75 | 4.55 | 7.5e-06 |
| | 200-300 vs. 300-400 | 0.31 | 3.21 | 0.1 | 0.92 |
| | 200-300 vs. 400-500 | 6.04 | 3.28 | 1.85 | 0.066 |
| | 300-400 vs. 400-500 | 5.74 | 3.2 | 1.79 | 0.074 |
| **Follow-up linear model results right hemisphere** | | | | | |
| **Parameter** | **Time** | **Estimate** | **SE** | **t** | **p** |
| Word frequency | 0-100 vs. 100-200 | 1.68 | 4.06 | 0.41 | 0.68 |
| | 0-100 vs. 200-300 | -9.79 | 3.08 | -3.17 | 0.0016 |
| | 0-100 vs. 300-400 | -8.22 | 3.11 | -2.64 | 0.0086 |
| | 0-100 vs. 400-500 | -6.54 | 3.34 | -1.96 | 0.051 |
| | 100-200 vs. 200-300 | -11.48 | 3.63 | -3.16 | 0.0017 |
| | 100-200 vs. 300-400 | -9.9 | 3.65 | -2.71 | 0.007 |
| | 100-200 vs. 400-500 | -8.23 | 3.85 | -2.14 | 0.033 |
| | 200-300 vs. 300-400 | 1.58 | 2.52 | 0.63 | 0.53 |
| | 200-300 vs. 400-500 | 3.25 | 2.8 | 1.16 | 0.25 |
| | 300-400 vs. 400-500 | 1.67 | 2.82 | 0.59 | 0.55 |
| SE = standard error. Sum of squares and mean squares in e-54 T. Estimate and SE in e-28 T. For F or p < .05, spin permutation was used to asses significance. See text for significant p values from spin permutation. | | | | | |
